# Redox regulation of the SARS-CoV-2 main protease provides new opportunities for drug design

**DOI:** 10.1101/2022.04.18.487732

**Authors:** Lisa-Marie Funk, Gereon Poschmann, Ashwin Chari, Fabian Rabe von Pappenheim, Kim-Maren Stegmann, Antje Dickmanns, Nora Eulig, Marie Wensien, Elham Paknia, Gabi Heyne, Elke Penka, Arwen R. Pearson, Carsten Berndt, Tobias Fritz, Sophia Bazzi, Jon Uranga, Ricardo A. Mata, Matthias Dobbelstein, Rolf Hilgenfeld, Ute Curth, Kai Tittmann

## Abstract

Besides vaccines, the development of antiviral drugs targeting SARS-CoV-2 is critical for stopping the current COVID-19 pandemic and preventing future outbreaks. The SARS-CoV-2 main protease (M^pro^), a cysteine protease with essential functions in viral replication, has been validated as an effective drug target. Here, we show that M^pro^ is subject to redox regulation and reversibly switches between the enzymatically active dimer and the functionally dormant monomer through redox modifications of cysteine residues. These include sulfenylation, disulfide formation between the catalytic cysteine and a proximal cysteine, and generation of an allosteric lysine-cysteine SONOS bridge that is required for structural stability under oxidative stress conditions, such as those exerted by the innate immune system. We identify homo- and heterobifunctional reagents that mimic the redox switching and possess antiviral activity. The discovered redox switches are conserved in main proteases from other coronaviruses, e.g. MERS and SARS-CoV, indicating their potential as common druggable sites.

## Introduction

The current COVID-19 pandemic, caused by the severe acute respiratory syndrome coronavirus 2 (SARS-CoV-2), constitutes the largest global health crisis in the recent past with at least six million deaths and approximately 0.5 billion cases worldwide (*1*). Although the development of vaccines has been instrumental in the reduction of severe progression and lethality of the disease, antiviral drugs are required to complement vaccination in high-risk groups, and for controlling sudden future outbreaks (*2*). Also, the genetic diversity and rapid evolution of SARS-CoV-2 has led to the emergence of virus variants, for which vaccination has reduced efficiency (*3–5*). As SARS-CoV-2 or related viruses are expected to remain a global threat in the future, the development of antiviral drugs becomes increasingly important.

Major therapeutic strategies for the treatment of COVID-19 include the application of neutralizing antibodies/nanobodies and small-molecule drugs targeting vital enzymes of the viral replication machinery (*6–9*). In the latter context, the SARS-CoV-2 main protease M^pro^ is a particularly promising drug target (*10, 11*). It proteolytically processes the viral polyproteins pp1a and pp1ab at no less than 11 cleavage sites, and thereby also ensures its own release. Its biological function in the viral replication cycle, along with the absence of a closely related human homologue, establish M^pro^ as a propitious drug target. The structure determination of SARS-CoV-2 M^pro^ sparked the development of several classes of inhibitors that bind either to the active site and covalently modify the catalytic cysteine or to allosteric sites (*11–18*). These efforts culminated in the design of Paxlovid™ (Pfizer), an orally administered FDA-approved antiviral drug which contains Nirmatrelvir, an inhibitor targeting SARS-CoV-2 M^pro^ (*19*).

M^pro^ is a cysteine protease that contains a catalytic dyad consisting of the nucleophilic cysteine 145 (C145) and histidine 41 (H41) (*11*). In total, M^pro^ contains 12 cysteine residues per chain (306 residues) (**Fig. 1A**, **SI Fig. 1**), which amounts to ~4% cysteines. This is a statistically unusual high cysteine abundance for a viral protein (*20*). The involvement of catalytic cysteine residues is a potential Achilles heel for viral replication, as oxidative stress exerted by the host innate immune system in response to viral infection may irreversibly oxidize the cysteines and thus inactivate the enzyme and block replication (*21,22*). Although M^pro^ resides in the cytoplasm, which is typically considered to be of reducing nature, it has been established that oxidative bursts or even physiological redox signaling based on enzymatic production of reactive oxygen species (ROS) such as H_2_O_2_ leads to local oxidizing conditions and subsequent oxidation of protein thiols in the cytosol (*23*). We recently reported the discovery of lysine-cysteine redox switches in proteins consisting of NOS (nitrogen-oxygen-sulfur) and SONOS (sulfur-oxygen-nitrogen-oxygen-sulfur) bridges (*24,25*). Interestingly, M^pro^ is amongst this class of proteins suggesting the possibility that it is redox regulated. Specifically, an allosteric SONOS bridge consisting of two cysteines (C22, C44) and one lysine (K61) within one protein chain was detected (**SI Fig. 2**). C44 is located on a flexible loop (residues 44-53) and points either towards the active site and contacts Y54 (“in” conformation) or towards the protein surface (“out” conformation). The flexibility of the loop is also confirmed by molecular dynamics (MD) simulations (**SI Fig. 3**) in good agreement with temperature-dependent structural data of M^pro^ (*26*). Independent structural studies have indicated that the catalytic cysteine C145 of M^pro^ is susceptible to oxidation and forms various oxidation products including mono-oxidized (sulfenic acid) and di-oxidized (sulfinic acid) species (**SI Fig. 4**), which rapidly interconvert to the tri-oxidized (sulfonic acid) form. The latter two forms are considered to be irreversible modifications and would lead to a dysfunctional enzyme and an arrest of viral replication (*27*).

**Figure 1.**
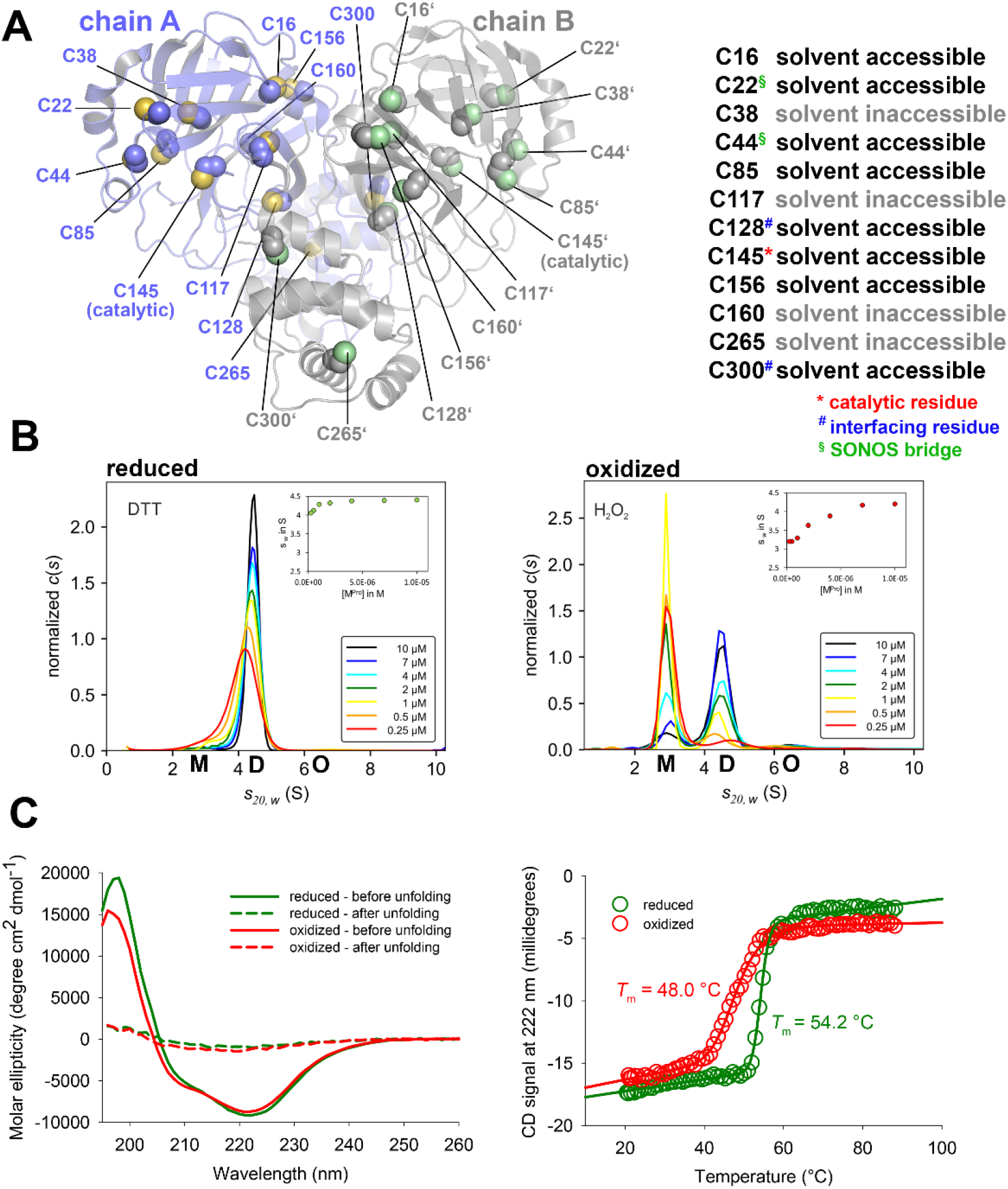
Structure and redox properties of SARS-CoV-2 main protease (M^pro^). (**A**) Structure of the M^pro^ dimer (pdb code 7KPH) highlighting the positions, structural properties and functions of cysteine residues. The two monomers of the functional dimer and corresponding cysteines are colored individually. A close up of the active site and proximal cysteines is shown in SI Fig. 1. (**B**) Sedimentation velocity analysis of SARS-CoV-2 M^pro^ in a concentration range from 0.25 to 10 μM under either reducing (left panel, 1 mM DTT) or oxidizing (right panel, 1 mM H_2_O_2_) conditions indicate a redox-dependent monomer ⇔ dimer equilibrium with apparent equilibrium constants of *K_app_* <0.25 μM for the reduced enzyme and of about 2.5 μM for the oxidized enzyme. Insets show s_w_ binding isotherms, as calculated from the corresponding c(s) distributions. Abbreviations: M, monomer (S_20,w_ = 2.9 S); D, dimer (S_20,w_ = 4.5 S); O, oligomers ((S_20,w_ = 6.3 S). (**C**) Secondary structure (left panel) and thermal unfolding (right panel) analysis of M^pro^ by far-UV CD spectroscopy under reducing and oxidizing conditions. Note the slightly reduced helical content (lower signal at 222 nm) and the decreased melting temperature of the oxidized enzyme (*T_m_* = 48.0 °C) versus the reduced counterpart (*T*_m_ = 54.2 °C). Further note the decreased cooperativity of unfolding (decreased steepness of transition) of the oxidized enzyme.

## Results and discussion

In order to test for potential redox regulation of M^pro^, we subjected the protein to different levels of oxidative insult using either 1 mM H_2_O_2_ as an upper limit for physiologically relevant oxidative stress conditions or 20 mM H_2_O_2_ as a superphysiological concentration (*28*). For 1 mM H_2_O_2_, we observed a progressive loss of enzymatic activity over time that correlates with a dissociation of the functional dimer and formation of the monomer as observed by size exclusion chromatography experiments (**SI Fig. 5**). Exposure of the oxidized protein to reducing conditions fully restores enzymatic activity and leads to the reconstitution of the dimer suggesting the existence of a redox switch. At 20 mM H_2_O_2_, however, enzymatic activity is irreversibly lost and almost no dimer dissociation is observed. To quantitatively assess the oligomeric equilibrium between monomer and dimer under reducing and oxidizing conditions, we performed analytical ultracentrifugation experiments (**Fig. 1B, SI Fig. 6**). Under reducing conditions, the M^pro^ dimer hardly dissociates even at the lowest protein concentration tested (0.25 μM) suggesting a *K*_D_^app^ of <250 nM. In contrast, under oxidizing conditions – either in the presence of H_2_O_2_ or, alternatively, O_2_ in a non-reducing buffer – M^pro^ exists in a monomer-dimer equilibrium. From the transition range of the s_w_ binding isotherms an apparent *K*_D_^app^ of about 2.5 μM could be estimated as reported before, however, an exact determination is prevented by the inability to fit the isotherms to a momer dimer two-state model (*11*). That notwithstanding the data clearly indicate that the equilibrium constant is at least one order of magnitude larger compared to the reduced protein. The shift of the oligomeric equilibrium under changing redox conditions is fully reversible (**SI Fig. 6**). The dissociation of a protein oligomer under oxidizing conditions is intriguing, typically, the reverse effect is observed for redox sensitive proteins where reduction of interchain disulfide bridges leads to deoligomerization (*29*). Secondary structure analysis of M^pro^ by far-UV circular dichroism (CD) spectroscopy under oxidizing and reducing conditions indicates structural differences between the two states (**Fig. 1C**). Upon oxidation, the fraction of α-helices slightly decreases (lower signal at 222 nm), while the fraction of ß-strands increases (higher signal at 210 nm). Also, the oxidized and reduced protein exhibit different thermal stabilities and different cooperativities of unfolding. The melting temperature of oxidized M^pro^ (*T_m_* = 48.0 °C) is 6 °C lower than that of the reduced enzyme (*T*_m_ = 54.2°C). The transition from the folded to the unfolded state is less steep in case of the oxidized protein indicating a reduced cooperativity of unfolding.

In order to identify the cysteine residues, which are part of the redox switch, we generated single-site variants with cysteine-to-serine substitutions for all 12 cysteines. In addition, we produced single, double and triple mutants with individual and combined exchanges of the SONOS bridge residues C22, C44 and K61 including residue Y54 that forms a hydrogen bond with C44. First, we analyzed the enzymatic activity of all variants under reducing and oxidizing conditions and tested whether a putative redox-induced change in activity is reversible (**Fig. 2A**, **SI Table 1**). As expected, variant C145S, in which the catalytic cysteine has been replaced, exhibits no measurable enzymatic activity. Most cysteine variants are almost as active as the wild-type protein with variants C117S and C44S being notable exceptions. Variant C44S exhibits the lowest residual activity (16%), while variant C117S is slightly more active (42% residual activity). A full kinetic analysis indicates that the reduction in activity is due to a decreased catalytic constant (*k*_cat_) rather than impaired substrate binding (**SI Fig. 7**). Double and triple variants with multiple exchanges of SONOS residues lead to enzyme variants with almost abolished enzymatic activity. Interestingly, variant Y54F, in which the tyrosine that interacts with SONOS residue C44 is replaced, shows a similar catalytic deficiency (19%) as variant C44S suggesting that both residues are required for full catalytic competence of the active site. To define the structural basis of the compromised enzymatic activity in variants C44S and Y54F, we crystallized both variants and compared the atomic structure with the known structure of the wild-type enzyme (*30*). In the case of variant C44S, we were able to obtain a structural snapshot of the covalent acyl intermediate formed with the C-terminal glutamine of a symmetry-related M^pro^ molecule (**SI Fig. 8**). This structural analysis reveals in both cases small structural changes mostly confined to the active site, notably of residue H41 that forms the catalytic dyad with C145 (**SI Fig. 8, 9**).

**Figure 2.**
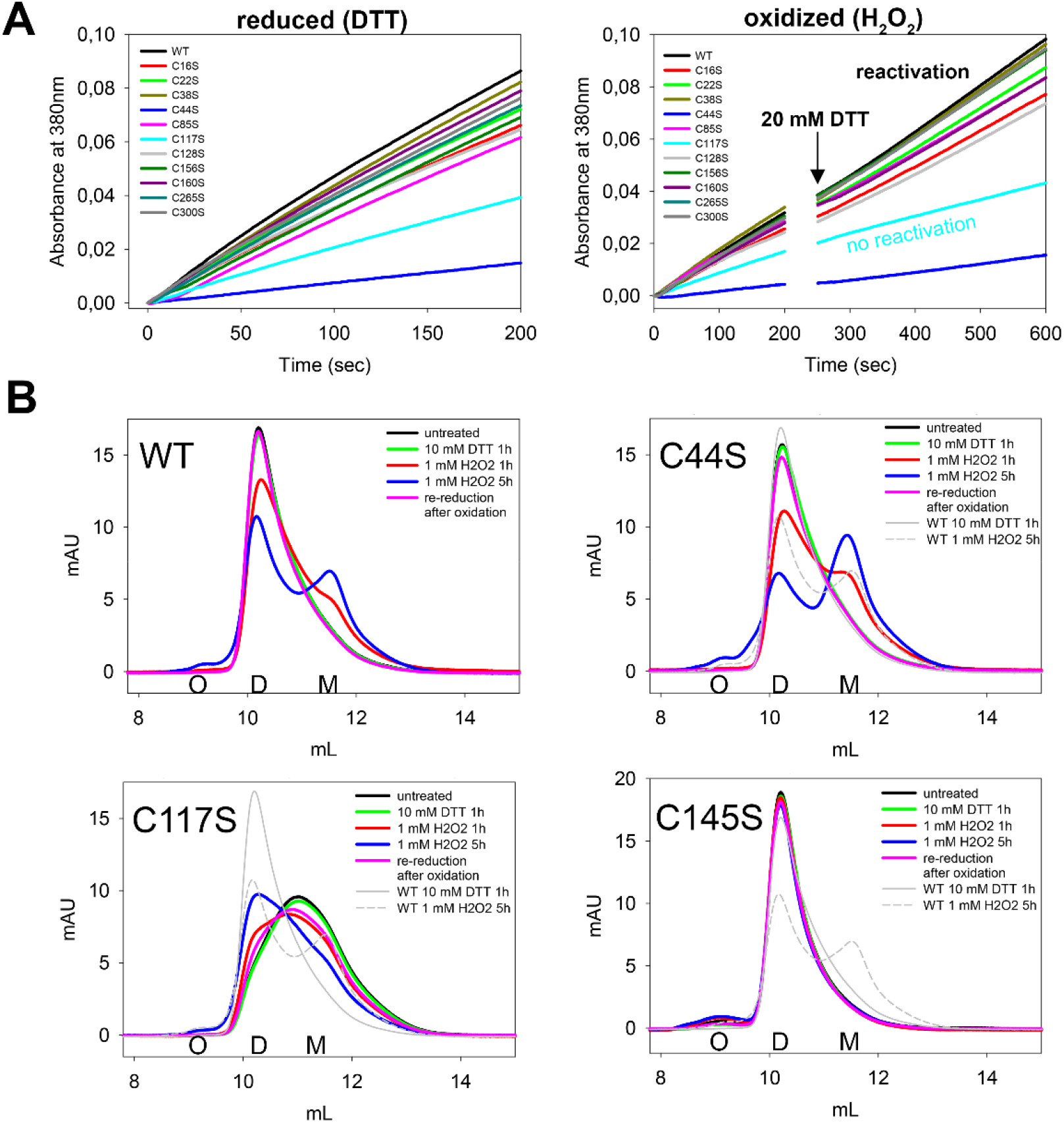
Redox-dependent enzymatic activity and oligomeric state of SARS-CoV-2 main protease (M^pro^) wild-type (WT) and cysteine variants. (**A**) Progress curves of substrate turnover for the reduced (left panel, 1 mM DTT) versus oxidized (right panel, 2 h 1 mM H_2_O_2_) enzyme. The relative activities are summarized in SI Table 1. In case of the oxidized enzyme, 20 mM DTT were added after a reaction time of 200 s to reactivate the enzyme by re-reduction. Reactivation was monitored up to a total reaction time of 10 min. Note that all enzyme variants except for C117S become reactivated. Variant C145S with a substitution of the catalytic cysteine is enzymatically inactive and not shown. All experiments were done in duplicate and with two independent biological replicates. (**B**) Gel filtration analysis of the oligomeric state of M^pro^ wild-type (WT) and selected, phenotypically outstanding cysteine variants in the reduced state and after different reaction times with H_2_O_2_. Abbreviations: O, oligomer; D, dimer; M, monomer. Note the progressive formation of the monomer with increasing oxidation times in case of the WT enzyme. For variant C44S, a larger fraction of the monomer is observed that likely reflects a kinetic rather than a thermodynamic effect (SI Fig. 12). Variant C117S is phenotypically unique in the stabilization of the monomer under reducing conditions and formation of the dimer upon oxidation. In contrast to the WT and all other cysteine variants tested, the redox switch on the quarternary level is not fully reversible for C117S. Variant C145S does not undergo monomerization in the course of oxidation under the conditions used highlighting the essential role of C145 for the redox switch.

Treatment of M^pro^ wild-type and variants with 1 mM H_2_O_2_ for 2 h on ice results in decreased enzymatic activities to a similar extent in all proteins (~3-fold reduction) pinpointing the central role of catalytic residue C145 as a major site of redox modification. Re-reduction of the protein with DTT leads to a reactivation of enzymatic activity in all cysteine variants except for C117S, where activity is irreversibly lost (**Fig. 2A**, **SI Table 1, SI Fig. 10**). For some of the SONOS variants such as C22S_C44S_K61A, a partial recovery of enzymatic activity can be observed but only after long incubation times with reductant (**SI Table 1, SI Fig. 11**).

The analysis of the monomer-dimer equilibrium by gel filtration experiments under either reducing or oxidizing conditions further substantiated the critical roles of C145, C117 and the SONOS residues for the redox switching of M^pro^ (**Fig. 2B, SI Tables 3 & 4**). Variant C145S was the only variant, for which no marked monomer formation was detectable under the conditions used. Variant C117S was unique in forming a detectable fraction of the monomer already under reducing conditions. Intriguingly, upon oxidation, the oligomeric equilibrium shifted to the dimer. Notably, the redox-dependent shift on the quaternary level is not fully reversible for variant C117S, while all other variants tested undergo a reversible switching. For SONOS variant C44S, we observed a larger fraction of the monomer that is likely to result from a faster oxidation reaction as the analytical ultracentrifugation experiments indicate similar dissociation constants for the variant under reducing and oxidizing conditions as for M^Pro^ wildtype (**SI Fig. 12**). While the analytical ultracentrifugation experiments for variant C117S are compatible with the gel filtration analysis, variant C145S undergoes dissociation upon oxidation in these experiments, albeit only at low concentrations (**SI Fig. 13**). This might indicate that the monomer-dimer equilibrium in this variant is not solely thermodynamically controlled but also kinetically (gel filtration experiments are conducted immediately after oxidative insult, ultracentrifugation analysis is preceded by an overnight incubation to allow the system to equilibrate). In general, the gel filtration experiments indicate that all variants, particularly these with substitutions of SONOS residues, tend to form higher oligomers/aggregates under oxidizing conditions suggesting a role of the SONOS bridge for structural stabilization (**SI Table 4**).

The comparative analysis of all proteins by far-UV CD spectroscopy regarding secondary structure content and thermal unfolding identified variants C145S and C117S as phenotypically conspicuous (**SI Fig.14**). In contrast to the wild-type enzyme, the increase of ß-strand elements at the expense of α-helical content upon oxidation is not observed for the oxidized proteins (**SI Table 5**). Also, the melting temperatures of the reduced and oxidized forms are almost identical (**SI Fig. 14, SI Table 6**). SONOS variants, in particular those containing an exchange of K61, are very susceptible to aggregation and exhibit an atypical early onset of thermal denaturation (30-35 °C) with almost no cooperativity of unfolding, indicating a very loosely structured protein (**SI Fig. 14**). This would imply a stabilizing function of the SONOS bridge under oxidizing conditions. The X-ray crystallographic analysis of the K61A variant in complex with the acyl intermediate formed between C145 and Q306 of a symmetry-related M^pro^ molecule reveals structural changes throughout the whole molecule with an r.m.s.d. of the Cα-carbons of 2.87 Å for chain A and 3.10 Å for chain B, respectively, compared to the wild-type structure (**SI Fig. 15**). The structural changes for K61A are clearly more pronounced than observed for variants C44S (0.89/0.91 Å) and Y54F (0.24 Å), highlighting the structural importance of K61.

We next set out to identify the redox modifications of M^pro^ by mass spectrometry-based redox proteomics and Western blot analysis (**Fig. 3**). Cysteines in M^pro^ exhibit a low sensitivity towards irreversible overoxidation (**Fig. 3A**). Only treatment with high concentrations of H_2_O_2_ leads to a substantially increased formation of sulfenylation and further overoxidation when incubated in presence of dimedone. Mass spectrometry identified cysteines 145, 156, and 300 as sulfenylated at high physiological H_2_O_2_ concentrations (1 mM) and cysteines C85, C117, C128, C145, C156, C265 and C300 as sulfonylated but only at 20 mM H_2_O_2_ (**Fig. 3B, SI Fig.16**). The existence of SONOS linked peptides could not be directly proven, but we noticed that C22 and C44 were only accessible for alkylation after reduction (following oxidation with H_2_O_2_) implicating a previous oxidation of both sites. As C22 and C44 were not found to be sulfenylated, sulfinylated, sulfonylated or in a disulfide linkage, this might be considered as indirect evidence for the existence of the SONOS bridge. Two disulfide-linked cysteine pairs, C145/C117 and C145/C156, were identified repetitively and exclusively in peptides of H_2_O_2_-oxidized MPro but not of reduced M^pro^ (**Fig. 3B**, **SI Fig. 17**). Since a substitution of C145 and C117 led to variants with impaired or abolished redox regulation (see above), but not in case of the C156S variant, we conclude that the C145/C117 disulfide is a decisive modification underlying the redox switching of M^pro^ rather than C145/C156. Inspection of the X-ray structure of M^pro^ determined in the reduced state indicates that residues C145 and C117 cannot directly form a disulfide bond as residue N28 lies in between the two side chains (**Fig. 3C**). Interestingly, N28 was previously demonstrated to be important for catalysis and dimer stability (*31*). MD simulations were carried out for M^pro^ with and without the disulfide linkage between C145 and C117. One of the main structural differences is the displacement of residue N28 (**SI Fig. 18**). Computed dimerization enthalpies show that the formation of the disulfide link reduces the latter by 4.2 kcal/mol (**SI Fig. 18**). We thus hypothesize that redox switching of M^pro^ entails a structural reorganisation upon oxidation that brings C145 and C117 into direct spatial proximity and expels N28 from its original position (switch 1 in **Fig. 3D**). The mutant data suggest that oxidation of the catalytic C145 to sulfenic acid is central to this structural change. Oxidative insult also leads to formation of the SONOS bridge formed between C22, C44 and K61. In the course of this reaction, C44 flips from the “in” to the “out” conformation (switch 2 in **Fig. 3D**). Overall, the redox switch mechanism protects M^pro^ against oxidative damage by forming a disulfide as a stable storage of the catalytic cysteine avoiding an irreversible overoxidation. The SONOS bridge structurally stabilizes the monomer that is formed upon oxidation by covalently tethering three structural elements within a monomer. Additionally, C44 seems to control the reactivity of the catalytic dyad in such a way that an oxidation of C44 and generation of SONOS leads to a reduced enzymatic activity and with that reduced propensity to become (over-)oxidized at C145. Overoxidation, as observed after treatment with 20 mM H_2_O_2_, however, overrides this mechanism and results in the occurrence of irreversibly inactivated dimers (**SI Fig. 5**). Since both disulfides and SONOS bridges are reversible modifications, enzymatic activity and the dimeric assembly of M^pro^ will be regained upon reduction ensuring the survival of the viral replication machinery under transient oxidative stress conditions. Sequence analysis indicates that the disulfide-forming residues C145/C117 and adjacent N28 are highly conserved in main proteases from different coronaviruses including SARS-CoV and MERS suggesting that these orthologs are also subject to redox regulation (**Fig. 3E, SI Fig.19**). A visualization of the general sequence conservation between viral main proteases, mapped on the structure of SARS-CoV-2 M^Pro^, is shown in **SI Fig. 20**. Based on sequence conservation, the SONOS redox modification seems to be a later evolutionary development and is found in SARS-CoV and SARS-CoV-2 but not in MERS or other coronaviruses (**Fig. 3E, SI Fig.19**).

**Figure 3.**
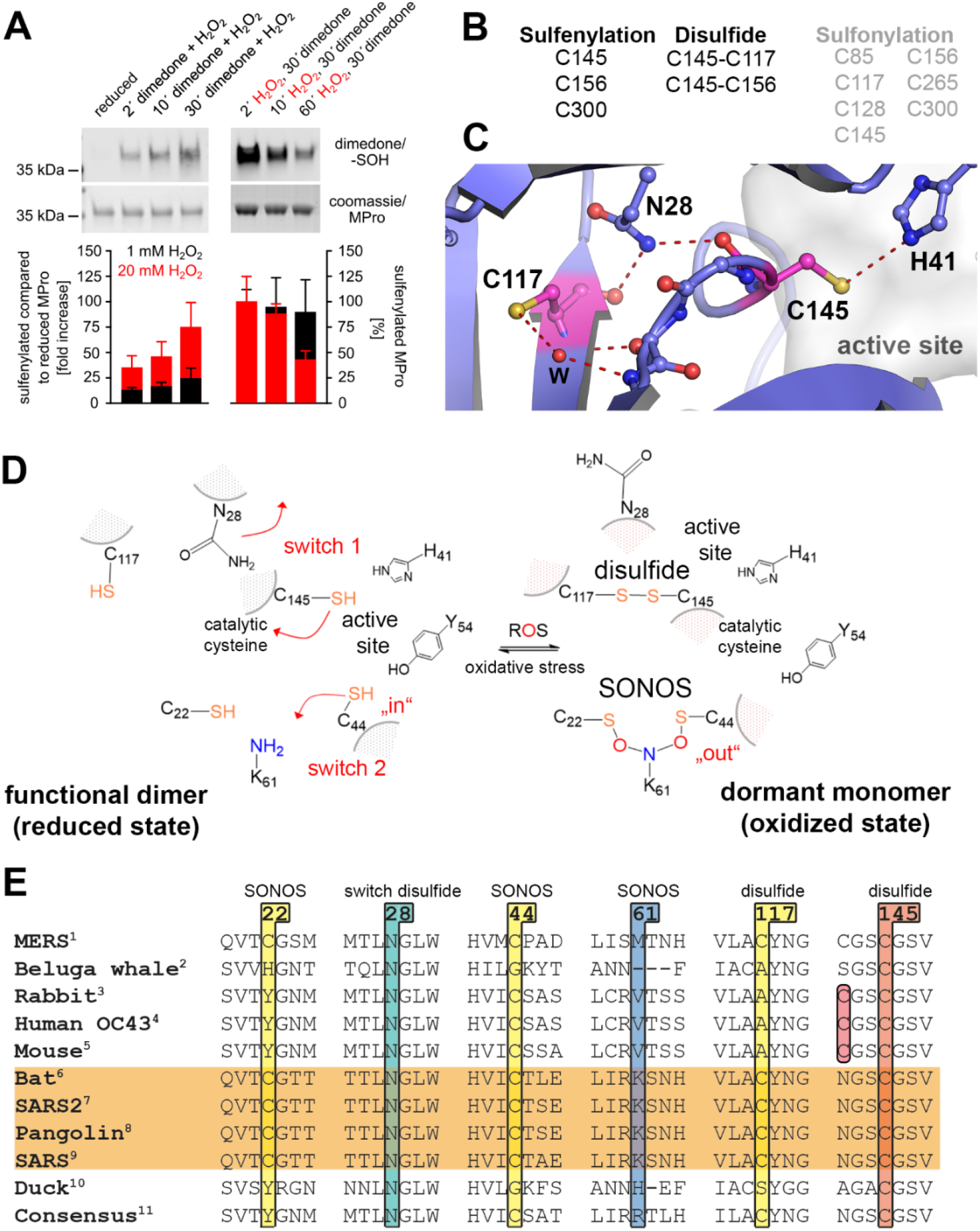
Analysis of redox modifications of SARS-CoV-2 main protease (M^pro^). (**A**) Western blot analysis of cysteine sulfenylation (formation of sulfenic acid). Reduced M^Pro^ was oxidized with either 1 mM (black bars) or 20 mM H_2_O_2_ (red bars). Sulfenylated thiols were trapped by addition of 5 mM dimedone added either simultaneously (left panel) or after preincubation with H_2_O_2_ (right panel) for/after indicated time points. Bars represent quantification of sulfenylated thiol/protein using western blots (n=2-6, SEM). (**B**) Mass specbased redox proteomics analysis of site-specific cysteine modifications in M^pro^ after oxidation with 1 mM H_2_O_2_ (sulfenylation, disulfide) or 20 mM H_2_O_2_ (sulfonylation). Representative mass spectra are shown in SI Fig. 16,17. (**C**) Structure of the M^pro^ active site and immediate vicinity highlighting the disulfide-forming C145 and proximal C117 interspaced by N28 (pdb code 7KPH). (**D**) Suggested redox switching mechanism of M^pro^. Under oxidizing conditions, catalytic C145 becomes sulfenylated inducing a structural transition (switch 1) that brings C145 and C117 together resulting in formation of the C117-C145 disulfide. This leads to a shift of the oligomeric equilibrium towards the monomeric state. Residues C22, C44 and K61 form the trivalent SONOS bridge (switch 2). (**E**) Sequence conservation of disulfide-forming residues C117 and C145 incl. the bridging residue N28 as well as of SONOS bridge-forming residues C22, C44 and K61 in coronaviruses. UniProtKB ID of polyprotein 1ab: ^1^ K9N7C7, ^2^ B2BW31, 3 H9AA60, ^4^ P0C6X6, ^5^ P0C6X9, ^6^ E0XIZ2, ^7^ P0DTD1, ^8^ A0A6G6A2G5, ^9^ P0DTD1, ^10^ A0A0F6WGL5, ^11^ From 67 sequences. Note that for proteins that do not possess an equivalent cysteine at position of C117, catalytic C145 is found to be in a C-X-X-C_145_ motif suggesting the possibility of another disulfide switch.

Finally, we tested whether the redox switching can be mimicked by redox-independent crosslinkers as a potential novel approach to design M^pro^ inhibitors. As cysteine and lysine residues constitute the genuine redox switches, we tested homobifunctional and heterobifunctional crosslinkers with warheads targeting thiol and amine functional groups. From the compounds tested, maleimidoacetic acid N-hydroxysuccinimide ester (MAH) gave the most promising results (**Fig. 4**). Addition of 1 mM MAH resulted in the almost complete dissociation of the M^pro^ dimer as monitored by both gel filtration and analytical ultracentrifugation experiments (**Fig. 4A/B**). Enzymatic activity is irreversibly lost after MAH incubation. In contrast, M^pro^-targeting drug Nirmatrelvir strongly stabilizes the dimer indicating different modes of inhibition for MAH and Nirmatrelvir. Mass spectrometric analysis identified catalytic C145 and K137 as MAH-binding residues (**Fig. 4C, SI Fig. 21**). As both residues are separated by ~20 Å in all structures determined to date, they could not simultaneously react with MAH without a structural change. Interestingly, the dimer interface, including residues E166 and N-terminal S1' from the second chain, is located between C145 and K137 rationalizing why a crosslink between the two residues destabilizes the dimer (**Fig. 4D**). We investigated the effect of MAH binding to C145 computationally. A model of the crosslinker binding at C145 was built and simulated for a total production time of 3.5 μs. The simulation shows that by covalent bonding of MAH to C145 alone the N28 residue is displaced (**SI Fig.22**), breaking the amide interaction with the backbone atoms of C117 and C145, and establishing a new hydrogen bond to the backbone of G146. This in turn facilitates C117 approaching C145 (**SI Fig. 22**). We further compared the dimerization energies. The MAH/C145 covalent bond structure has a penalty of 5.4 kcal/mol for dimer formation, showing that even before the K137-C145 crosslink formation the oligomerization energetics are already affected (**SI Fig. 18D**). A similar dimer-destabilizing effect, albeit not as quantitative as in case of MAH, is observed upon addition of the homobifunctional crosslinker bismaleimidoethane (BMOE) that preferentially crosslinks C145 and C117 thus mimicking the redox switch (**SI Fig. 23**). Using a cell-based SARS-CoV-2 infection model we can demonstrate as proof-of-concept that both MAH as well as BMOE exhibit antiviral activity (more than 50-fold reduced virus progeny) without affecting the survival of host cells (**SI Fig. 24, 25**).

**Figure 4.**
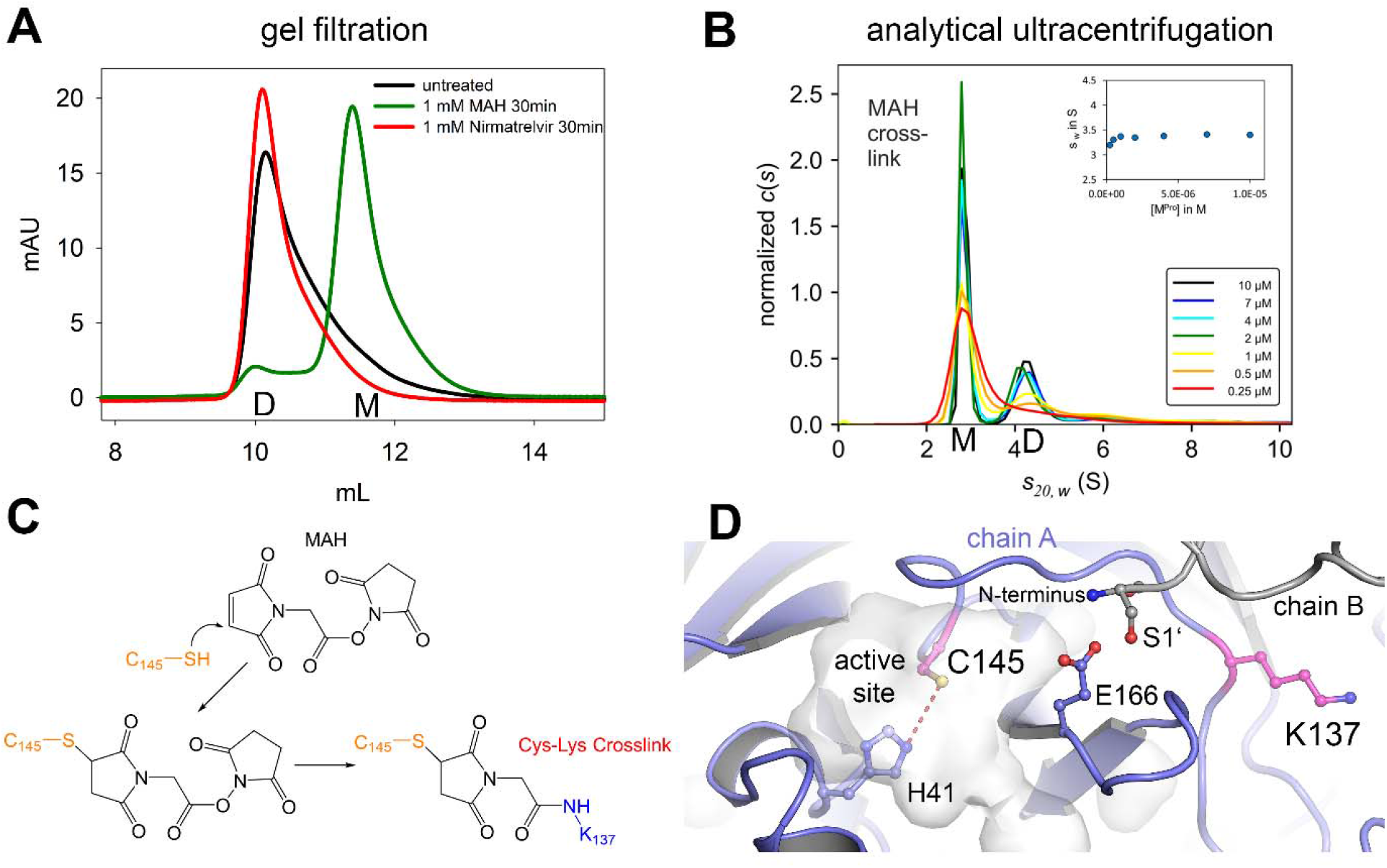
Impact of the heterobifunctional crosslinker MAH on the oligomeric equilibrium of M^pro^. (**A**) Gel filtration analysis of M^pro^ after addition of 1 mM MAH, 1 mM nirmatrelvir (Paxlovid™) and untreated as control. Abbreviations: D, dimer; M, monomer. Note that addition of MAH entails formation of the monomer, whereas addition of nirmatrelvir stabilizes the dimeric state. (**B**) Sedimentation velocity analysis of SARS-CoV-2 M^pro^ in a concentration range from 0.25 to 10 μM after incubation with 1 mM MAH. Inset shows the s_w_ binding isotherm, as calculated from the corresponding c(s) distributions. Abbreviations: M, monomer (s_20,w_ = 2.9 S); D, dimer (s_20,w_ = 4.5 S). Note the preferential destabilization of the dimer in support of the gel filtration experiments (see A). (**C**) Chemical mechanism of MAH crosslinking to M^pro^ as suggested by mass spec analysis that identified catalytic C145 (thiol) and K137 (amine) as main reaction sites. (**D**) Structure of the M^pro^ active site and dimer interface vicinity highlighting the MAH crosslink-forming C145 and K137 (pdb code 7KPH). Note that the monomermonomer interaction between E166 and S1', which is critical for dimer formation, is located between C145 and K137, providing the structural basis for destabilization of the dimeric assembly upon MAH crosslinking.

## Conclusions

In summary, we have reported on a hitherto unknown mode of redox regulation of SARS-CoV-2 M^pro^ that protects the redox-vulnerable catalytic cysteine and the structural integrity of the protein under oxidative stress conditions. Additional redox regulatory mechanisms might involve glutathionylation of cysteines as recently reported (*32*). The detected redox switches in the main protease seem to be widespread amongst coronaviruses and it is likely that other viral cysteine proteases such as the papain-like protease have evolved similar defense mechanisms. As the redox switching can be mimicked by non-redox chemistry, this offers novel opportunities in the design of inhibitors targeting viral cysteine proteases using the redox switches as common druggable sites.

## Supporting information

Appendix 1 - Gel filtration experiments

Appendix 2 - CD spectra

Appendix 3 - Unfolding data

Appendix 4 - Crosslinking experiments

## Acknowledgements

This study was supported by the Max-Planck Society and the DFG-funded Göttingen Graduate Center for Neurosciences, Biophysics, and Molecular Biosciences GGNB (to KT). We further thank the Coronavirus Forschungsnetzwerk Niedersachsen (COFONI) for funding project 13FF22 to MD. The analytical ultracentrifuge Beckman Coulter Optima AUC was funded by the Deutsche Forschungsgemeinschaft (DFG) – INST 192/534-1 FUGG. The study was also supported through DFG grants MA5063/4-1 (to RAM) and 417677437/GRK2578 (to CB). We acknowledge access to beamline P14 at DESY/EMBL Hamburg, Germany and thank G. Bourenkov and T. Schneider for local support. We thank R. Golbik for discussion of the circular dichroism data and Lidia Litz for excellent technical assistance of the analytical ultracentrifugation experiments.

