## Supplementary Data for "Redox regulation of the SARS-CoV-2 main protease provides new opportunities for drug design"

### Supplementary Information

### Experimental Procedures

#### Experimental Procedures

##### General information

The protein concentration was determined by UV/Vis spectroscopy using the absorbance signal at 280 nm and the molar extinction coefficient ($\varepsilon_{MPro}$ = 32890 $M^{-1}cm^{-1}$), which was calculated according to Gill and von Hippel (*S1*).

##### Mutagenesis

Variants of SARS-CoV-2 $M^{pro}$ were generated by site-directed mutagenesis PCR using the QuikChange site-directed mutagenesis protocol (Stratagene, La Jolla, CA, USA) and expression vector pGEX-6P1 NSP5 (https://mrcppureagents.dundee.ac.uk/reagents-view-cdna-clones/703227). Correctness of the introduced mutations was confirmed by complete sequencing of the gene.

The following primers were used:

| | |
|---|---|
| C16S | 5´-P-AGTGGAAGGTtctATGGTACAGGTGACATG-3´ upr<br>5´-P-TTGCCGGACGGAAACGCC-3´lwr |
| C22S | 5´-P-ACAGGTGACAtccGGCACCACAA-3´ upr<br>5´-P-ACCATACAACCTTCCACTTTGC-3´ lwr |
| C38S | 5´-P-CGTAGTCTATtctCCTCGTCATGTC-3´upr<br>5´-P-TCGTCTAACCACAACCCA-3´ lwr |
| C44S | 5´-P-TCATGTCATCtccACCTCTGAGG-3´ upr<br>5´-P-CGAGGGCAATAGACTACG-3´ lwr |
| C44A | 5´-P-TCATGTCATCgccACCTCTGAGGAC-3´ upr<br>5´-P-CGAGGGCAATAGACTACG-3´ lwr |
| C85S | 5´-P-CATGCAGAATtccGTCCTTAAAC-3´ upr<br>5´-P-CTATGACCAATAACGCGC-3´ lwr |
| C117S | 5´-P-AGTGTTAGCGtccTATAACGGCA-3´ upr<br>5´-P-GAAAAGGTCTGACCAGGC-3´ lwr |
| C128S | 5´-P-TGTGTATCAGtctGCTATGCGTCC-3´ upr<br>5´-P-CCAGAGGGACTGCCGTTA-3´ lwr |
| C145S | 5´-P-TAATGGCAGCtctGGTTCGGTGG-3´ upr<br>5´-P-AGGAAGCTGCCTTTGATC-3´ lwr |
| C156S | 5´-P-CGACTACGATagcGTTAGCTTCT-3´ upr<br>5´-P-ATGTTAAAGCCCACCGAAC-3´ lwr |
| C160S | 5´-P-CGTTAGCTTCtccTATATGCACC-3´ upr<br>5´-P-CAATCGTAGTCGATGTTAAAG-3´ lwr |
| C265S | 5´-P-GCTGGATATGtctGCCAGTCTGAAAG-3´ upr<br>5´-P-ACAGCAATGCCCGTCTGT-3´ lwr |
| C300S | 5´-P-GGTGCGTCAGtctAGCGGTGTCA-3´ upr<br>5´-P-ACATCGAAGGGAGTGAACTCATC-3´ lwr |
| K61A | 5´-P-CCTGATCCGCgcaTCCAACCACA-3´ upr<br>5´-P- AGATCTTCGTAATTCGGATTG-3´ lwr |
| Y54F | 5´-P- CAATCCGAATttcGAAGATCTCCTG-3´ upr<br>5´-P- AGCATGTCCTCAGAGGTG-3´ lwr |

**Expression**

For recombinant expression, vector pGEX-6P1 NSP5 containing the $M^{Pro}$ gene was transformed into BLR(DE3) chemically competent *E. coli* cells (Novagen, Merck Biosciences, affiliate of Merck KGaA, Darmstadt, Germany), containing an pREP4 plasmid, according to Inoue et al. (*S2*). The bacteria were grown in LB media (*S3*) containing 50 µg/mL kanamycin sulfate and 100 µg/µL carbenicillin (disodium salt) at 37 °C until an optical density at 600 nm ($OD_{600}$) of 0.6 was reached. The cells were then incubated at 18 °C for 30 min until an $OD_{600}$ of 0.8. Subsequently, gene expression was induced by addition of 500 µM isopropyl-ß-D-thio-galactopyranoside (IPTG) for ~20 h at 18 °C. The cells were harvested by centrifugation at 5750x*g* and either directly used or flash frozen in liquid nitrogen and stored at -80 °C until usage.

**Protein purification**

All purification steps were performed at 4 °C or on ice. Cells were resuspended in buffer A (20 mM Tris/HCl pH 7.8, 5 mM imidazole, 150 mM NaCl, 1 mM dithiothreitol), supplemented with 100 µM phenylmethanesulfonyl fluoride, 0.5 mg/mL lysozyme (AppliChem GmbH Darmstadt, Germany), 5 mM $MgCl_2$ and 5 µg/mL DNaseI (Thermo Fisher Scientific Braunschweig, Germany), and subsequently lysed by five passages through a LM10 Microfluidizer® High Shear Fluid Homogenizer (Microfluidic Corp, Newton, MA, USA).
Next, the lysate was centrifuged at 75000x*g* for 30 min and the thereby obtained supernatant was loaded onto a HisTrap™ HP 3x5 mL column (GE Healthcare Munich, Germany). The His6x-$M^{Pro}$ fusion protein was then eluted with buffer B (20 mM Tris/HCl pH 7.8, 300 mM imidazole, 150 mM NaCl, 1 mM dithiothreitol) and subsequently dialyzed against 2 L buffer A overnight. To remove the His-tag, 1 mg PreScission Protease was added per 10 mg $M^{Pro}$. The cleaved His-tags and non-cleaved protein were then separated from untagged $M^{Pro}$ via affinity chromatography as described above. The PreScission Protease was removed via an additional affinity chromatography step, employing a GSTrap™ HP column (GE Healthcare Munich, Germany). $M^{Pro}$ was then treated with 1 mM EDTA and subjected to size exclusion chromatography using a HiLoad 16/60 Superdex 75 prep grade gel filtration column (GE Healthcare, Munich) in buffer C (20 mM Tris/HCl pH 7.8, 150 mM NaCl, 1 mM dithiothreitol). The purified protein was either directly used for experiments or supplemented with 20% (*v*/*v*) glycerol, flash frozen with liquid nitrogen and stored at -80°C until usage.

**Steady-State kinetics**

For steady-state kinetic analysis of enzymatic activity of $M^{Pro}$ wild-type and variants under reducing and oxidizing conditions, the cleavage of an artificial $M^{Pro}$ peptidic substrate (Ac-Abu-Tle-Leu-Gln-AMC, Biosynth Carbosynth, Switzerland) was monitored spectrophotometrically at 380 nm in a UV–Vis spectrometer (V-750, Jasco GmbH, Germany).
Prior to the kinetic measurements, $M^{Pro}$ in assay buffer (20 mM Tris pH 7.3, 100 mM NaCl, 1 mM EDTA) was incubated with either 1 mM $H_2O_2$ or 1 mM dithiothreitol (DTT) for 2 h. For measurements of reactivation, the protein was first incubated with 1 mM $H_2O_2$ for 2 h on ice, whereupon the $H_2O_2$ was removed using a 5 mL HiTrap Desalting column (GE Healthcare Munich, Germany). Subsequently, oxidized $M^{pro}$ was (re)-reduced by incubation with 20 mM DTT for 3 or 20 h. The reaction was started by adding 1 µM $M^{Pro}$ to a preincubated reaction mix (200 µL) containing 200 µM peptide substrate and either 1 mM $H_2O_2$ or 1 mM DTT in assay buffer at 20 °C. The change in absorption was continuously monitored at 380 nm ($\varepsilon_{AMC}$ = 2400 $M^{-1}cm^{-1}$). Initial rates were estimated by linear regression of the absorbance signal over the first 10 s of the measurements or, in cases in where substrate activation was observed, using eq 1

$$A_{340}(t) = A_0 - \Delta ss \cdot t + \frac{\Delta ss - \Delta_0}{k_{obs}} \cdot \left[1 - \exp^{(-k_{obs} \cdot t)}\right]$$

Eq 1

in which $A_0$ denotes the starting absorbance at 380 nm, Δss the absorbance changes at steady-state (steady-state rate), $\Delta_0$ the absorbance changes at $t$ = 0 (initial rate), and $k_{obs}$ the first-order rate constant of activation.

**Secondary structure and thermal unfolding analysis**

To analyze secondary structure contents and thermal stability of $M^{Pro}$ wild-type and variants, far-UV circular dichroism (CD) spectra and thermal unfolding data were collected using a circular dichroism spectrometer (Chirascan, Applied Photophysics, UK). Far-UV CD spectra were collected in a range of 195-260 nm and using a concentration of 0.2 mg/ml protein treated with either 1 mM $H_2O_2$ or 1 mM DTT for 2 h on ice, in 100 mM $Na_2HPO_4$ pH 7.8, with a step size of 1 nm and at least 20 accumulations for 0.5 sec per wavelength. Secondary structure contents were calculated using the CDNN software (*S4*).

Thermal unfolding was monitored at a wavelength of 222 nm in a temperature range from 20-95 °C (real sample temperature was determined using a temperature probe) with a ramping speed of one °C/min. Each temperature data point was collected for 10 sec.

**Analytical ultracentrifugation (AUC)**

Sedimentation velocity experiments (SV) were performed in analytical ultracentrifuges ProteomeLab XL-I or Optima AUC (Beckman Coulter, USA) at 50000 rpm and 20 °C using An-50 Ti rotors. Concentration profiles were measured using the absorption scanning optics at 230 nm with 3 or 12 mm standard double sector centerpieces filled with 100 µl or 400 µl sample, respectively. Stock solutions of SARS-CoV-2 $M^{pro}$ and mutants thereof were dialyzed overnight against buffers containing 0.15 M NaCl and 20 mM Tris pH 7.3 in the absence (AUC buffer) or presence of 1 mM DTT (AUC buffer + DTT). After dilution to concentrations in the range of 0.25 to 10 µM, samples were allowed to equilibrate for 23 h at room temperature before SV analysis, since it has been found that the monomer-dimer equilibrium of $M^{pro}$ is slow on the time-scale of centrifugation (*11*).

To test whether $M^{pro}$ can be regenerated after removal of DTT, $M^{pro}$ stock solution was first dialyzed overnight against AUC buffer, afterwards diluted to 0.25 to 10 µM in AUC buffer + DTT. Samples were allowed to equilibrate for 23 h at room temperature before SV analysis. In a second experiment following dialysis against AUC buffer and dilution in AUC buffer to final concentration of 0.5 to 10 µM and another 23 h of incubation at room temperature, a 20 mM DTT stock solution (in AUC buffer) was added to a final concentration of 1 mM. $M^{Pro}$ was further incubated for 23 h at room temperature before SV analysis. Addition of DTT stock solution resulted in a slight dilution of the samples (0.47 to 9.5 µM).

For $M^{pro}$ oxidation by $H_2O_2$, DTT containing stock solutions were transferred to AUC buffer by ZEBA Spin Desalting Colums (Thermo Scientific), diluted to 25 µM and incubated with 1 mM $H_2O_2$ for 2 hours on ice. Subsequently, $H_2O_2$ was removed by ZEBA Spin Desalting Colums, samples were centrifuged for 20 min at 15,000 xg and protein concentrations were determined spectrophotometrically. Proteins were diluted to 0.25 to 10 µM and were analysed by SV about 2 h after dilution and at least 3 hours after $H_2O_2$ treatment. A similar protocol was applied for chemical crosslinking of $M^{pro}$ with heterobifunctional crosslinker maleimidoacetic acid N-hydroxysuccinimide ester (MAH), except that $H_2O_2$ treatment was replaced by incubation with MAH for 30 min on ice and 50 mM sodium dihydrogen phosphate pH 7.3 was used as a buffer. Untreated proteins were analysed in 50 mM sodium dihydrogen phosphate pH 7.3 as a control.

For data analysis, a model for diffusion-deconvoluted differential sedimentation coefficient distributions (continuous c(s) distributions) implemented in the program SEDFIT (*S5*) was used. Partial specific volume and extinction coefficient of the protein as well as buffer density and viscosity, were calculated from amino acid and buffer composition, respectively, by the program SEDNTERP (*S6*) and were used to calculate protein concentration and correct experimental s-values to $s_{20,w}$.

Signal-averaged s-values $s_w$ were obtained by integration of the c(s) distributions in the s-value range where monomers and dimers were observed using the program GUSSI (*S7*) and plotted as a function of concentration to obtain binding isotherms for the monomer-dimer equilibrium of SARS-CoV-2 $M^{pro}$.

#### Analytical size exclusion chromatography

To analyze the distribution of higher oligomers/aggregates, dimers and monomers of $M^{Pro}$ under reducing and oxidizing conditions, 25 µM $M^{Pro}$ in assay buffer were pre-incubated with either 1 mM $H_2O_2$ or 1 mM DTT for 1-5 h on ice and then loaded onto a Superdex 75 increase 10/300 GL column (GE Healthcare, Munich) via ÄKTA Pure 25M (GE Healthcare, Munich) at 6 °C. The peak heights and integrals were analysed using the UNICORN 7.1 software.

#### Crystallization and Cryoprotection

$M^{pro}$ crystals were grown at 20 °C using the hanging-drop vapor diffusion method with a reservoir solution containing 0.1 M MES pH 6.5, 8-10% PEG 3350, 1.5% DMSO and 0.1 M sodium citrate. 1 µL of reservoir solution was mixed with 1 µL protein solution containing 10 mg/mL $M^{Pro}$ in buffer C (20 mM Tris/HCl pH 7.8, 150 mM NaCl, 1 mM DTT). Cryoprotection was carried out using 20 % (*v*/*v*) glycerol in well solution, soaking the crystals for up to 90 seconds.

#### X-ray data collection, processing and model building

Diffraction data of $M^{Pro}$ single crystals (variants C44S, Y54F, K61A) were collected using synchrotron radiation at beamline P14 of the DESY/EMBL Hamburg, Germany at a wavelength of 0.827 or 0.976 Å. Data were collected at cryogenic temperature (100 K) with an EIGER 16M detector. Processing using anisotropic cut-off limits was performed using autoPROC (*S8*), which calls on the XDS package (*S9*),the CCP4 suite of programs (*S10*) and STARANISO (*S11*).

Subsequent refinement and model building was performed employing Phenix.REFINE (*S12*) and COOT (*S13*). Phasing was performed using MOLREP (*S14*) using the published $M^{Pro}$ structure (PDB ID 6LU7) as starting model. The geometry of the structural models was validated using MOLPROBITY (*S15*). Structural representations were prepared using PyMOL (*S16*). Crystallographic statistics are provided in Supplementary Table 2.
The refined structural protein models and corresponding structure-factor amplitudes have been deposited under PDB accession codes 7ZB6 (C44S), 7ZB7 (Y54F) and 7ZB8 (K61A). The Ramachandran statistics are 91.45 % in the favoured, 8.55 % in the allowed and 0 % in the outlier region for 7ZB6; 98.03 %, 1.64 % and 0.33 % for 7ZB7; and 87.99 %, 9.87 % and 2.14 % for 7ZB8.

**Redox proteomics**

$M^{Pro}$ was analyzed to study **a**) sulfenylation after western transfer (using the BioRad system) and **b**) to determine site-specific oxidative modifications via mass spectrometry. For a), $M^{Pro}$ was incubated for 30 min with 10 mM DTT on ice. DTT was removed using Zeba spin columns (Thermo Scientific). An amount of 0.5, 1.0, or 5.0 µg of reduced MPro were incubated in 20 mM Tris/HCl pH 7.8, 150 mM NaCl on ice for different periods of time (2 – 60 min) with 1 or 20 mM $H_2O_2$. Dimedone (5 mM) was either added simultaneously or after pre-incubation with $H_2O_2$. Remaining thiols were blocked by addition of 100 mM NEM (N-Ethylmaleimide). After SDS-PAGE, proteins were transferred to nitrocellulose membranes and sulfenylation was visualized by anti-dimedone antibodies (*S17*). b) Similar treated $M^{Pro}$ (see above) was directly applied for mass spectrometry or Coomassie-stained proteins were prepared for mass spectrometry. In addition, $M^{Pro}$ was treated with DTT (1 mM, 2 h on ice) and $H_2O_2$ (1 mM, 2 h on ice and 20 mM, 30 minutes on ice). The thereby obtained protein was either used for crosslinking after gel-filtration and buffer exchange to phosphate buffered saline including 1 mM EDTA, or cysteines were blocked with 100 mM NEM or, alternatively, with 55 mM iodoacetamide in 50 mM ammonium hydrogen carbonate. Crosslinking of $M^{Pro}$ was carried out by incubating 10 µg $M^{Pro}$ in a total volume of 20 µl phosphate buffered saline including 1 mM EDTA and 1 mM crosslinker (BMOE or MAH) for 30 minutes at 22 °C. The reaction was stopped by adding 1 µl 1 M Tris pH 7.5 and 4x sample buffer including DTT followed by polyacrylamide gel separation. $M^{Pro}$ was separated under reducing (+150 mM DTT) or non-reducing conditions (without DTT) in polyacrylamide gels and stained with Coomassie brilliant blue essentially as described (*S18*). Protein-containing bands were cut out of the gel and - depending on the experiment - reduced with DTT and alkylated with iodoacetamide as described (*S18*) or only alkylated with iodoacetamide. Finally, the $M^{Pro}$ samples were in-gel digested with 0.1 µg chymotrypsin in 23 µl of 100 mM Tris-HCl and 10 mM $CaCl_2$ in water (pH

7.8) overnight. Resulting peptides were extracted from the gel and resuspended in 0.1% trifluoroacetic acid as previously described (*S18*). Subsequently, peptides were separated using an Ultimate 3000 rapid separation liquid chromatography system (Thermo Fisher Scientific) as described before (*S18*). Briefly, peptides were loaded on a 2 cm length trap column for 10 minutes and subsequently separated over 54 minutes on a 20 cm C18 analytical column. Eluting peptides were directly sprayed into the mass spectrometer via a nanosource electrospray interface. An Orbitrap Fusion Lumos (Thermo Fisher Scientific) mass spectrometer, operated in positive mode, was used for the analysis of $M^{Pro}$ peptides. First, precursor spectra were recorded in the orbitrap in profile mode (resolution 60000, scan range 400-1800 m/z, maximum injection time 50 ms, AGC target 100000). Thereafter, 2-10 fold charged precursors were selected by the quadrupole (isolation window 1.6 m/z, minimum intensity 50000) fragmented via higher-energy collisional dissociation and analyzed in the orbitrap and afterwards newly selected and fragmented with collision-induced dissociation and analysis in the orbitrap. Fragment spectra were recorded in centroid mode (resolution 30000, maximum injection time 120 ms, AGC target 50000, scan range: auto). The cycle time was 2 seconds, already fragmented precursors were excluded from isolation for the next 60 seconds. Peptide and crosslink identification were carried out with MaxQuant version 2.0.3.0 (Max-Planck Institute for Biochemistry, Planegg, Germany) with standard parameters if not stated otherwise. The $M^{Pro}$ amino acid sequence was used as search template, following variable modifications were considered: acetylation (N-terminus), oxidation (methionine), carbamidomethylation (cysteine), glutathionylation (cysteine), di-oxidation (cysteine), tri-oxidation (cysteine). Depending on the analysed samples, additional modifications with NEM (cysteine), NEM + water (cysteine) and dimedone (cysteine) were considered. Crosslink searches were enabled by screening for disulfides (-2.0157), links between cysteines by BMOE (+220.0484) and BMOE + water (+238.059), links between cysteines with MAH (+137.0113) and MAH + water (+155.0219) and between cysteines and lysines with MAH (+137.0113) and MAH + water (+155.0219). The match-between-runs option was enabled, proteins and peptides were identified at a false discovery rate of 1%. Data analysis was carried out in excel based on “evidence” and “crosslinkMsms” tables. Here, identified spectra were counted for each run or intensities for all modified peptide variants were summed up per analysed sample. Disulfide crosslinks were accepted upon the following criteria: found in at least two independent experiments, a minimum of 4 identified spectra in at least one experiment, not found in the reduced control sample (1 mM DTT, 2 h on ice).

##### Cell culture

Vero E6 cells (Vero C1008) were maintained in Dulbecco's modified Eagle's medium (DMEM with GlutaMAX$^{TM}$, Gibco) supplemented with 10% fetal bovine serum (Merck), 50 µg/mL streptomycin (Gibco), 50 units/mL penicillin, 10 µg/mL ciprofloxacin (Bayer) and 2 µg/mL tetracycline (Sigma) at 37 °C in a humidified atmosphere with 5% $CO_2$.

##### MAH/BMOE treatment and SARS-CoV-2 infection

20,000 cells per well were seeded into 24-well-plates and incubated overnight at 37 °C. Cells were treated with either 20 mM MAH (Sigma-Aldrich, diluted in PBS, pH adjusted to 7.5), 20 mM BMOE (Sigma-Aldrich, diluted in PBS + 5% DMSO, pH adjusted to 7.5) or the PBS control for 1 h before infection, and then throughout the time of infection, using medium containing 2% fetal bovine serum (FBS). Cells were infected with virus stocks corresponding to $1*10^7$ RNA-copies of SARS-CoV-2 (= 30 FFU) and incubated for 48 h at 37 °C, as described (*S19*). Cell morphology was assessed by bright field microscopy.

##### Quantification of lactate dehydrogenase release to determine cytotoxicity

The release of lactate dehydrogenase (LDH) into the cell culture medium of MAH- or BMOE-treated cells was quantified by bioluminescence using the LDH-Glo$^{TM}$ Cytotoxicity Assay kit (Promega). 10% (*v*/*v*) Triton X-100 was added to untreated cells for 15 min to determine the maximum LDH release, whereas the medium background (= no-cell control) served as a negative control. Percent cytotoxicity was calculated using the following formula and reflects the proportion of LDH released to the media compared to the overall amount of LDH in the cells.

$$\text{Cytotoxicity (\%)} = 100 \times \frac{\text{(Experimental LDH Release - Medium Background)}}{\text{(Maximum LDH Release Control - Medium Background)}}$$

##### Quantitative RT-PCR for SARS-CoV-2 quantification

For RNA isolation, the SARS-CoV-2-containing cell culture supernatant was mixed with the Lysis Binding Buffer from the MagNA Pure LC Total Nucleic Acid Isolation Kit (Roche) to inactivate the virus. The viral RNA was isolated as described (*S19*) and quantitative RT-PCR was performed according to a previously established RT-PCR assay (*S20*), to quantify SARS-CoV-2 RNA yield. The amount of SARS-CoV-2 RNA determined upon infection without any treatment was defined as 100%, and the other RNA quantities were normalized accordingly. A two-sided unpaired Student's t-test was calculated using GraphPad Prism 9.

**Immunofluorescence analyses**

Vero E6 cells were treated/infected as indicated. After 48 hours of SARS-CoV-2 infection, the cells were washed once in PBS and fixed with 4% formaldehyde in PBS for 1 hour at room temperature. After permeabilization with 0.5% (*v*/*v*) Triton X-100/PBS for 30 min and blocking in 10% FBS/PBS for 10 min, primary antibodies were used to stain the SARS-CoV-2 Spike (S; GeneTex#GTX 632604, 1:2000) and Nucleoprotein (N; Sino Biological #40143-R019, 1:8000) overnight. The secondary Alexa Fluor 488 donkey anti-mouse IgG and Alexa Fluor 546 donkey anti-rabbit IgG (Invitrogen, 1:500, diluted in 10% FBS/PBS) antibodies were added together with DAPI for 1 h at room temperature. Slides with cells were mounted with DAKO and fluorescence signals were detected by microscopy (Zeiss Axio Scope.A1).

**Immunoblot analysis**

Vero E6 cells were treated/infected as indicated. After 48 hours of SARS-CoV-2 infection, the cells were washed once in PBS and then harvested in RIPA lysis buffer (20 mM TRIS-HCl pH 7.5, 150 mM NaCl, 10 mM EDTA, 1% (*v*/*v*) Triton-X 100, 1% deoxycholate salt (*w*/*v*), 0.1% (*v*/*v*) SDS, 2 M urea), supplemented with protease inhibitors. After sonication and equalizing the amounts of protein, samples were separated by SDS-PAGE. To determine the presence of viral proteins, the separated proteins were transferred to a nitrocellulose membrane, blocked in 5% (*w*/*v*) non-fat milk in TBS-T for 1 h, and incubated with primary antibodies at 4 °C overnight, followed by incubation with peroxidase-conjugated secondary antibodies (donkey anti-rabbit or donkey anti-mouse IgG, Jackson Immunoresearch). The SARS-CoV-2 Spike (S; GeneTex#GTX 632604, 1:1000) and Nucleoprotein (N; Sino Biological #40143-R019, 1:5000), and GAPDH (abcam ab8245, 1:5000) were detected using Immobilon Western Substrate (Millipore).

**Analysis of $M^{Pro}$ sequence conservation**

The dataset used for analysis was generated using the replicase polyprotein 1ab of SARS-CoV2 (Uniprot-ID P0DTD1) as query for *blastp*. A total of 67 1ab polyprotein sequences were selected from the resulting search for further analysis. These were truncated to the $M^{Pro}$-squence. Alignment and tree-generation were performed using ClustalOmega (*S21*). Tree visualization was performed using iTOL (*S22*). Alignment analysis was performed using Jalview (*S23*).

#### Quantum chemical calculations

##### Parametrization of NOS and SONOS

Given that there are no available parameters in the standard Amber forcefield sets for any cysteine-lysine covalent linkage, we carried out a parameterisation of the NOS and SONOS bonds. In a first stage, we built the parameters for the single NOS bond using a small model system ($CH_3NOSCH_3$). We minimized the structure, employing the Gaussian 16 RevA.03 software package (*S24*) at the B3LYP-D3(BJ) level of theory (*S25-27*) and the def2-SVP basis set (*S28,29*). The partial atomic charges were assigned using the RESP procedure,(*S30*) at the HF/6-31G* level. The Seminario approach (implemented in the CartHess2FC tool provided with the Amber20 program package) was then employed for the R enantiomeric form of the NOS, in order to obtain the parameters for the NT-OS-S angle (*S31*). We then performed the scans along the dihedral angles CT-NT-OS-S, H1-CT-NT-OS, H1-CT-S-OS, NT-OS-S-CT and OS-NT-OS-S, at the same DFT level previously mentioned. At this point, a genetic algorithm was employed in order to fit the dihedral angles potentials (http://www.ub.edu/cbdd/?q=content/small-molecule-dihedrals-parametrization) to the DFT values, considering the non-bonded interactions. All the other parameters were provided by the antechamber tool for Amber type atoms.

For the SONOS, we made use of the crystal structure coordinates of the C22, C44 and K61 residues present in the X-ray structure (PDB: 7JR4). We capped the backbones at the C and N atoms saturating them with hydrogens. We then optimized the system at the aforementioned DFT level, constraining all the non-hydrogen backbone atoms to their crystallographic positions. At this stage, we obtained the partial atomic charges at HF/6-31G* level, subtracting the values of the added cap H atoms. The antechamber tool was used to generate the Amber parameters to the oxidized lysine residue and the disulfide bridge atom types were used for the cysteines bound to the lysine.

##### Starting structure for the disordered loop of the SONOS containing $M^{pro}$ dimer

The missing residues (#46, #47 and #48) were modelled by setting up a system formed by the mentioned residues and the neighbouring #45 and #49, which were capped at the terminal C and N, saturating them with hydrogen atoms. The system was then minimized using the semiempirical PM6 Hamiltonian (*S32*) constraining the non-hydrogen atoms of residues #45 and #49. The obtained cartesian coordinates of the minimized non-hydrogen atoms were then manually added to the X-ray crystal structure (PDB: 7JR4).

**Modelling of the MAH containing amino acid**

In order to model the MAH-C145 bonded system, a model system was built capping at the C145 backbone C and N atoms, saturated with hydrogens. The H atoms were relaxed at the B3LYP-D3(BJ)/ def2-SVP level of theory. The partial atomic charges were assigned using the RESP procedure, subtracting the values of the added cap H atoms, at HF/6-31G* level. The forcefield parameters were assigned with the antechamber tool using Amber atom types.

**Replica-exchange constant pH simulations**

Molecular dynamic simulations were performed setting HIS41, GLU47, ASP48, LYS61, HIS64, HIS163 and HIS164 as titratable. Every other GLU, ASP and LYS are set as charged groups and the protonation states for the other histidines were set as: HID80, HIE172 and HIE246. We have performed the simulations for the reduced, disulfide and SONOS containing dimeric and monomeric systems.

The RE-cpH simulations were performed with the AMBER20 software package,(*S33,34*) using sander and pmemd, employing the ff10 force field (*S35,36*). The protein was set in a cuboid periodic box leaving an 8 Å distance between the protein atoms and the periodic box wall. TIP3P water molecules neutralized with Na+ and Cl- counter ions were included (*S37*). The cut off for non-bonded interactions was set to 8 Å, employing particle-mesh Ewald summation with a fourth-order B-spline interpolation and a tolerance of $10^{-5}$. The non-bonded list was updated every 50 fs, and the MD time step was set to 2 fs, employing the SHAKE algorithm to constrain bonds involving hydrogen atoms (*S38*).

The H atoms and residues #44, #46, #47 and #48 of the system were first minimized for 2,000 cycles (1,000 with steepest descendent and 1,000 with conjugate gradient) by restraining the rest of the atoms with a 1,000 kcal/mol/Å$^2$ force constant. The system was then minimized for another 3,000 cycles (1,000 with steepest descendent and 2,000 with conjugate gradient) restraining the non-hydrogen backbone atoms of the protein with a 10 kcal/mol/Å$^2$ force constant. Finally, the system was minimized for 10,000 cycles (2,000 with steepest descendent and 8,000 with conjugate gradient), allowing all the atoms to relax.

The system was heated from 0 to 300 K in the first 800 ps of an overall 1 ns run, using a NVT ensemble, employing Langevin dynamics with a collision frequency of 5 ps$^{-1}$. The system was then equilibrated for 1 ns in the NPT ensemble at 300 K and with isotropic position scaling and a relaxation time of 5 ps. The production phase is done using 16 replicas employing the same ensemble and parameters as in the equilibration phase. The production is carried for 128 ns, attempting to change the protonation state every 200 fs and attempting replica exchanges every 4 ps. The heating and production phases were performed using graphics processing unit (GPUs) (*S39,40*).

**Molecular dynamic simulations**

The molecular dynamic simulations were performed setting all the GLU, ASP and LYS residues charged and the histidines in the following protonation states: HID41, HID64, HID80, HIE163, HIE172, HIE246 and HIP164.

The simulations were performed with the AMBER20 software package, using sander and pmemd, employing the ff99SB force field (*S41*). The protein was set in a cuboid periodic box of 8 Å, between the protein and the periodic box wall, of TIP3P water molecules neutralized with Na+ and Cl- counter ions. The cut off for non-bonded interactions was set to 8 Å, employing particle-mesh Ewald summation with a fourth-order B-spline interpolation and a tolerance of $10^{-5}$. The non-bonded list was updated every 50 fs, and the MD time step was set to 2 fs, employing the SHAKE algorithm to constrain bonds involving hydrogen atoms.

The H atoms and residues #44, #46, #47 and #48 of the system were first minimized for 2,000 cycles (1,000 with steepest descendent and 1,000 with conjugate gradient) by restraining the rest of the atoms with a 1,000 kcal/mol/Å$^2$ force constant. The system is then minimized for another 3,000 cycles (1,000 with steepest descendent and 2,000 with conjugate gradient) restraining the non-hydrogen backbone atoms of the protein with a 10 kcal/mol/Å$^2$ force constant. Finally, the system was minimized for 10,000 cycles (2,000 with steepest descendent and 8,000 with conjugate gradient), allowing all the atoms to relax.

The system was then heated from 0 to 300 K in the first 800 ps of an overall 1 ns run, using a NVT ensemble, employing Langevin dynamics with a collision frequency of 5 ps$^{-1}$. The system was then equilibrated for 1 ns in NPT ensemble at 300 K with isotropic position scaling and a relaxation time of 5 ps. The production phase was carried out using the same ensemble and parameters as in the equilibration phase. The production was performed for 150 ns for the SONOS and 600 ns for the reduced and disulfide containing dimers. The MAH-covalent bound system was simulated for a total time of 3.5 µs. This was performed in an attempt to identify how this chemical modification could change the interactions around the catalytic cysteine. The heating and production phases were performed using graphics processing units (GPUs).

**Analysis of the molecular dynamics**

All the structural analysis of the molecular dynamic simulations were performed using the CPPTRAJ (V4.25.6) tool from AmberTools (V20.15) (*S42*). The dimerisation energies were analysed using the MMPBSA.py implementation (*S43*) making use of 500 frames extracted from the first 150ns of each system MD.

### Supplementary Information

### Figures

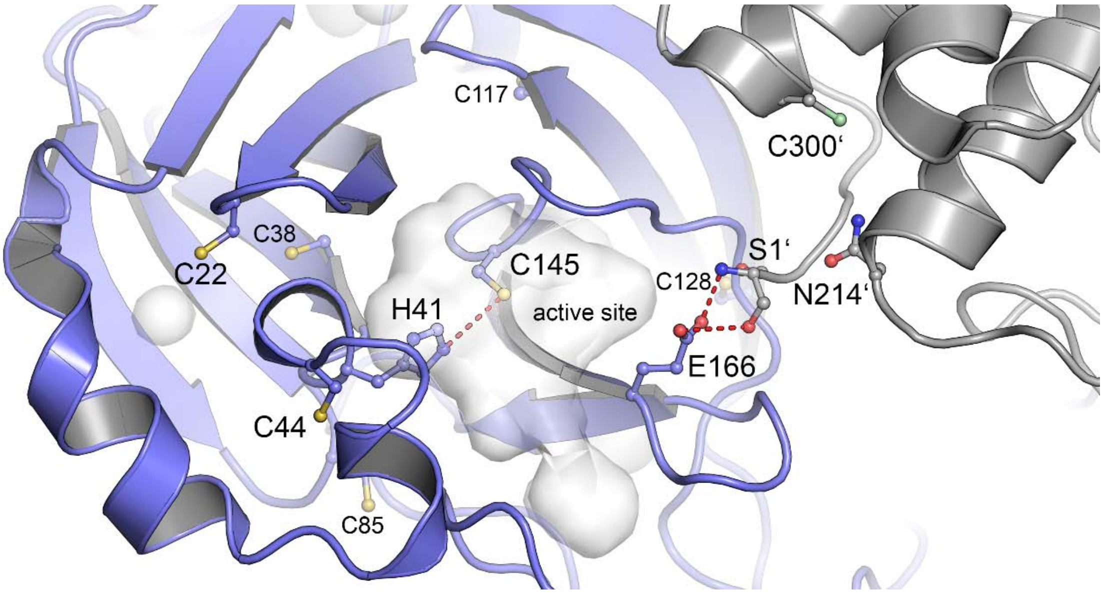


**SI Figure 1**. Structure of $M^{pro}$ showing the positions of cysteine residues in a monomer and at the dimer interface (pdb code 7KPH). The two monomers of the functional dimer and corresponding cysteines are colored individually. Residues contributed by the second monomer are marked with an apostrophe. The transparent volumes indicate cavities including the active site pocket. Important hydrogen-bond interactions are highlighted including that between catalytic residues C145 and H41 as well as that between E166 and N-terminal residue S1’ at the dimer interface. The structure of the dimer is shown in Figure 1A of the main manuscript.

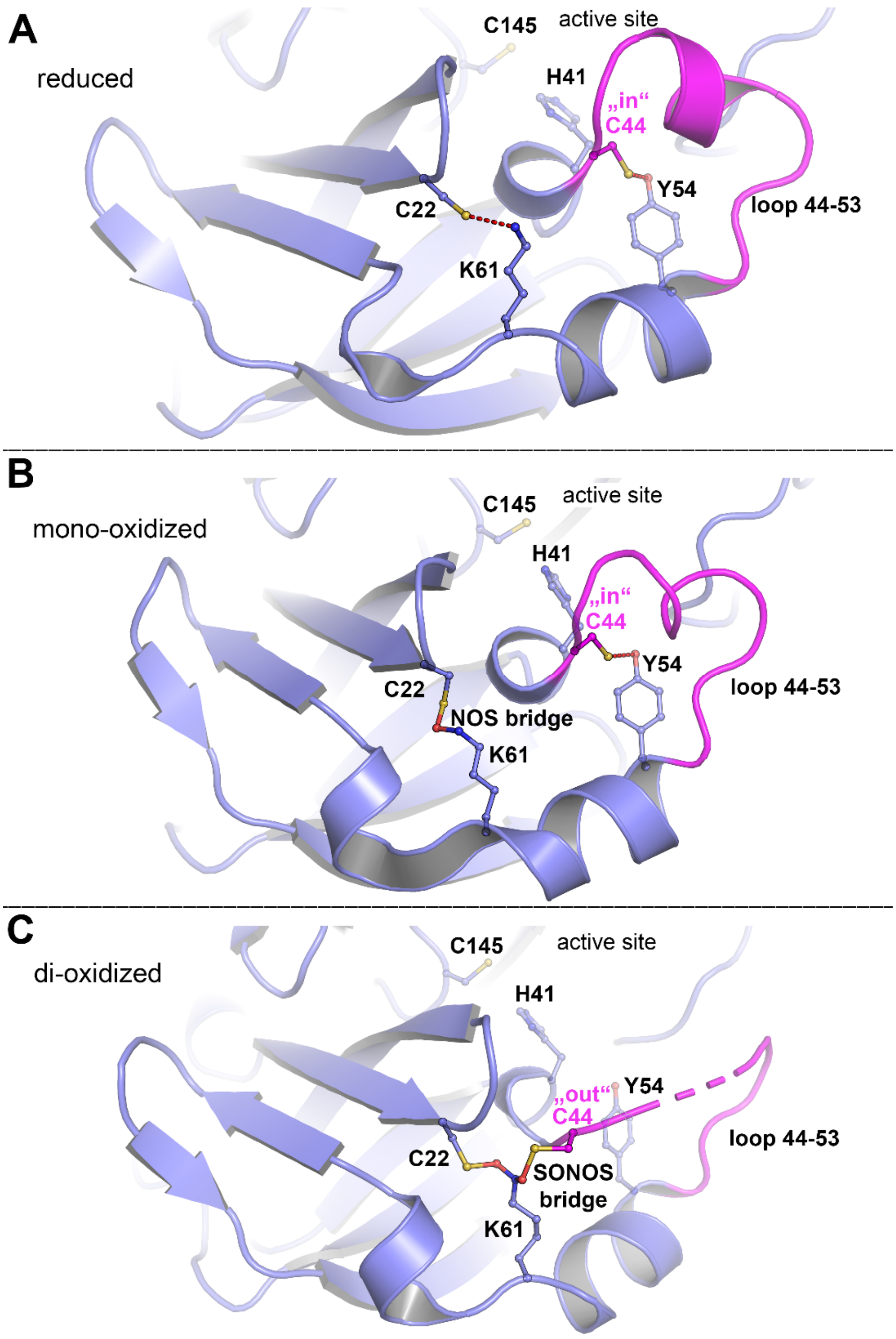


**SI Figure 2.** NOS and SONOS bridges in the main protease M$^{pro}$ from SARS-CoV-2. (**A**) Structure of M$^{pro}$ in the reduced state (pdb code 7JR3) showing the redox switch at the protein surface formed by residues C22, C44 and K61 as well as the active site with residues C145 (catalytic nucleophile), H41 and Y54. A mobile loop bearing C44 is indicated in magenta. Note that C44 is in the "in" conformation and interacts with Y54. (**B**) Structure of M$^{pro}$ in a mono-oxidized state with an NOS bridge formed between C22 and K61 (pdb code 6XMK). C44 is found in the "in" conformation. (**C**) Structure of M$^{pro}$ in a di-oxidized state with a SONOS bridge formed between C22, K61 and C44 (pdb code 7JR4). C44 is found in the "out" conformation.

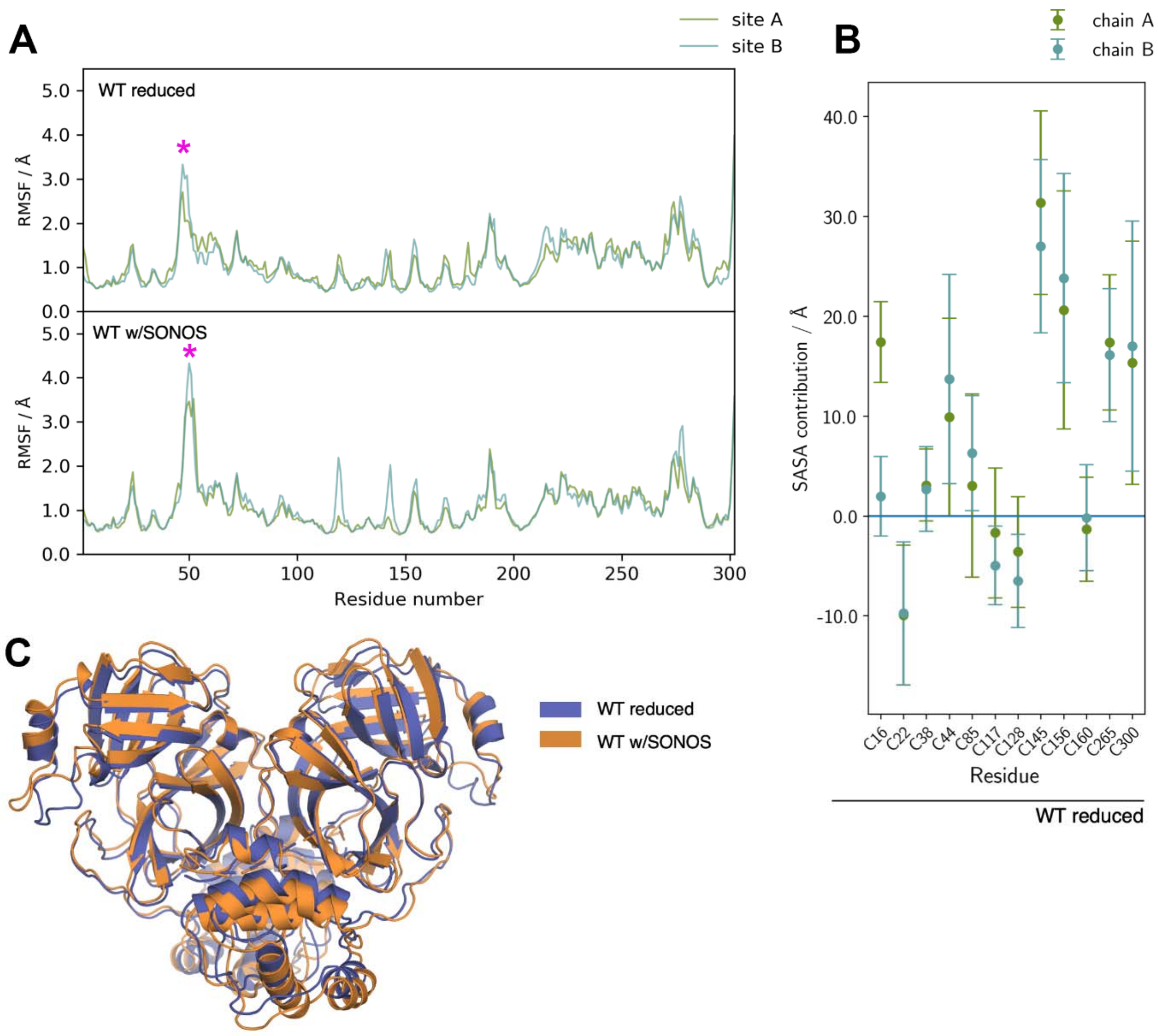


**SI Figure 3.** (**A**) Root-mean-square fluctuations extracted from the molecular dynamics trajectories for the WT M$^{pro}$ in its reduced state (top) and with a SONOS bridge (bottom). The flexible loop 44-53 (highlighted by an asterix in magenta) is visible and both structures show very similar fluctuation patterns. (**B**) Solvent accessible surface area (SASA) contributions of individual cysteines to the full SASA of the system, following the linear combinations of pairwise overlaps (LCPO) method of Weiser et al. (*33*). Positive contributions show cysteines which are exposed to solvent and do not hinder neighbouring residues. Negative values indicate cysteines which are blocked or effectively block from solvent contact other residues, with a net result of reducing solvent exposure. The values are based on the molecular dynamics simulations of the reduced WT. (**C**) Root-mean-square difference overlap using the alpha-carbon positions for the averaged MD structures of the reduced WT and with a SONOS bridge (RMSD of 1.44 Å). Both structures are very close to identical with the exception of the 44-53 loop.

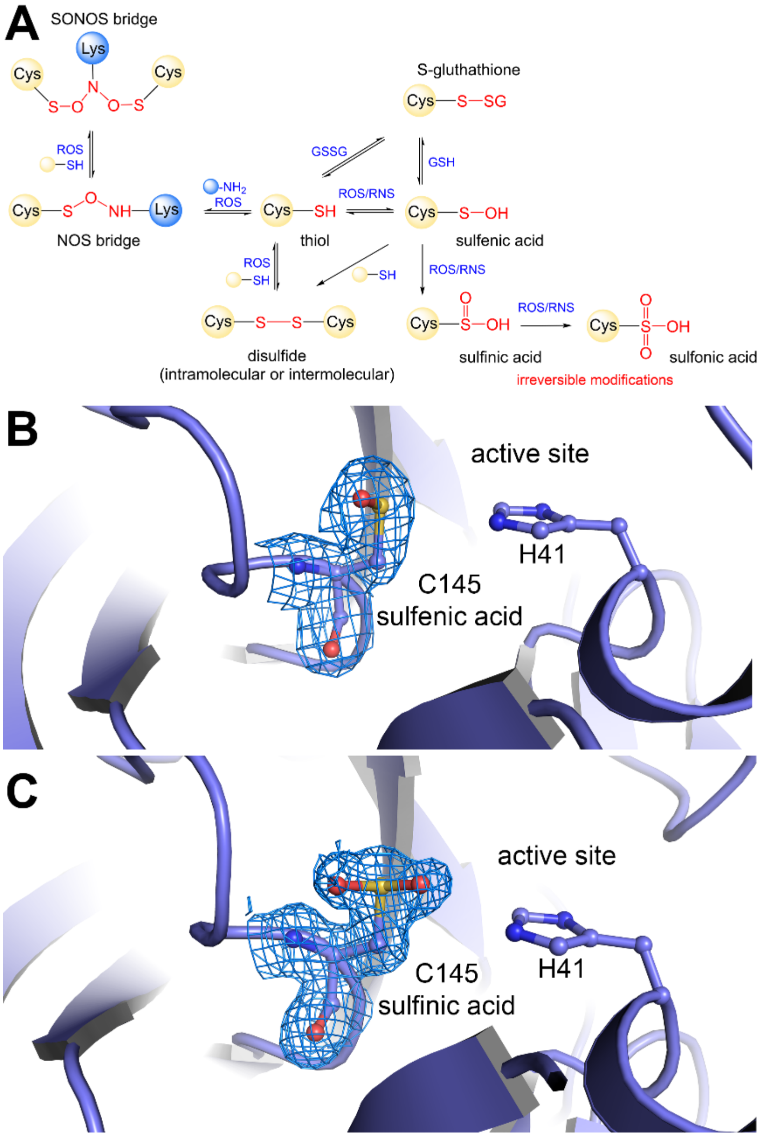


**SI Figure 4.** Redox modifications of the catalytic cysteine C145 in $M^{pro}$ from SARS-CoV-2. (**A**) Redox modifications of cysteines in proteins showing key species involved in oxidative and reductive transformations. Abbreviations: Cys, cysteine; Lys, lysine; ROS, reactive oxygen species; RNS, reactive nitrogen species; GSH, glutathione; GSSG, glutathione disulfide. (**B**) Structure of $M^{pro}$ with catalytic C145 in the mono-oxidized sulfenic acid state (pdb code 6XKF). The structural model of C145 is superposed with the 2mFo-DFc electron density map at a contour level of 1σ. (**C**) Structure of $M^{pro}$ with catalytic C145 in the di-oxidized sulfinic acid state (pdb code 6XKH). The structural model of C145 is superposed with the 2mFo-DFc electron density map at a contour level of 1σ.

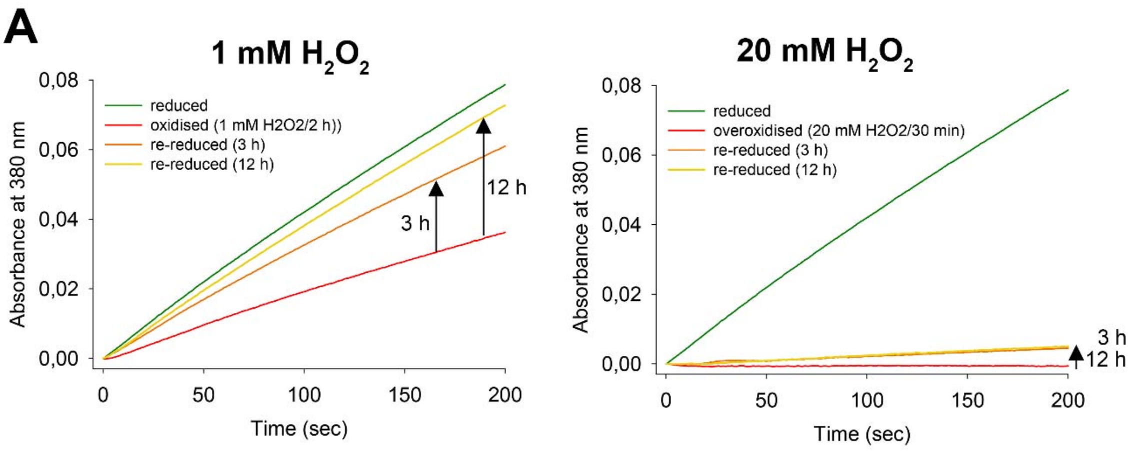


| | reduced | (over)oxidized | re-reduced (3h) | re-reduced (12h) |
|---|---|---|---|---|
| WT (oxidation) | 100.00% | 33.45% | 69.79% | 83.92% |
| WT (overoxidation) | 100.00% | 0.13% | 4.43% | 4.93% |

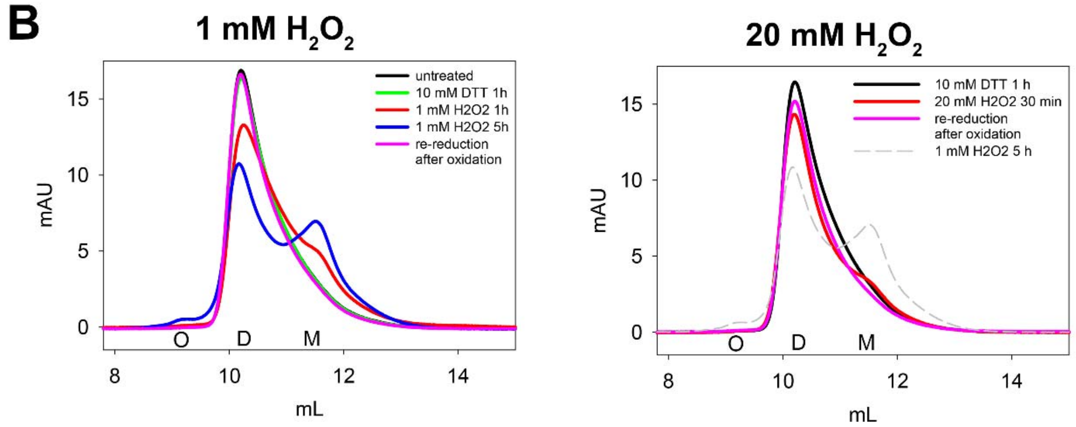


**SI Figure 5.** Redox-dependent enzymatic activity and oligomeric equilibrium of $M^{pro}$. (**A**) Progress curves of $M^{pro}$-catalyzed substrate turnover in the reduced state, after oxidation with either 1 mM $H_2O_2$ (left panel) or 20 mM $H_2O_2$ (right panel) and after re-reduction with reductant DTT. Experimental details are provided in the Supplementary Methods. The estimated relative enzymatic activities are summarized in the accompanying table. An activity of 100% refers to that of the reduced enzyme. Note that an oxidation with 20 mM $H_2O_2$ leads to an irreversible inactivation of $M^{pro}$, while treatment with 1 mM $H_2O_2$ entails a reversible loss of activity. (**B**) Gel filtration analysis of the oligomeric state of $M^{pro}$ wild-type in the reduced state, after different reaction times with either 1 mM $H_2O_2$ (left panel) or 20 mM $H_2O_2$ (right panel) and after re-reduction. Abbreviations: O, oligomer; D, dimer; M, monomer. Note the progressive formation of the monomer with increasing oxidation times after reaction with 1 mM $H_2O_2$. Re-reduction restores the dimer.

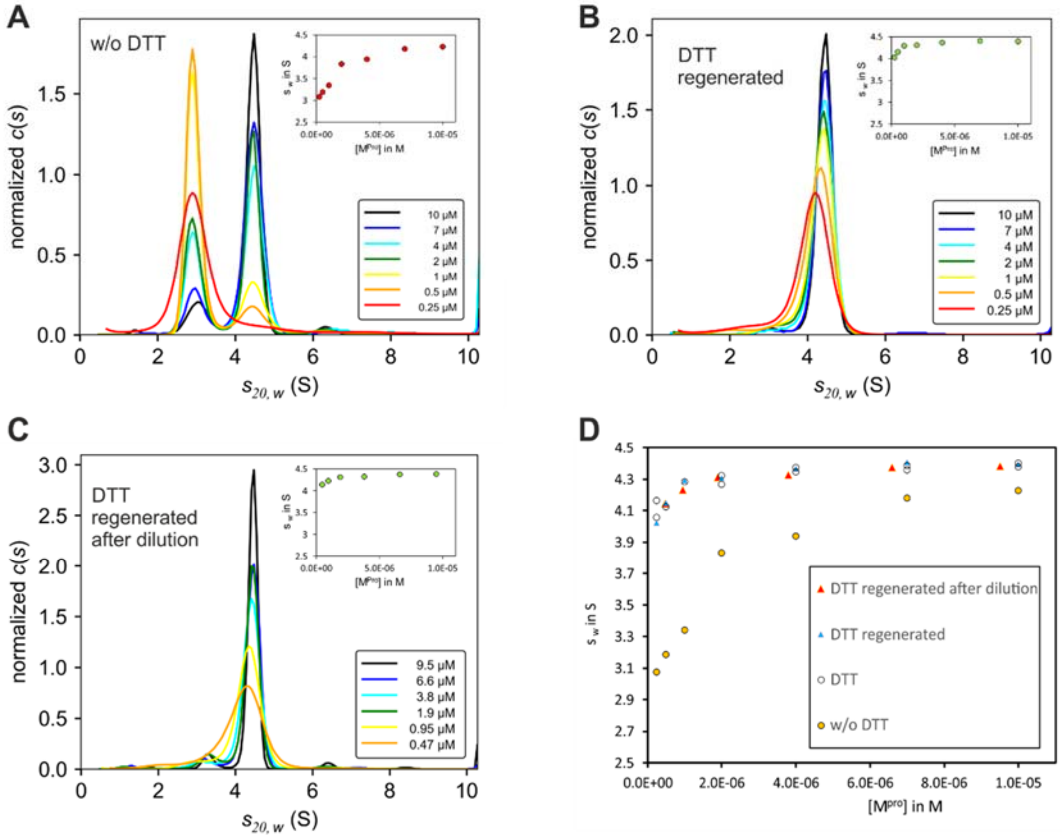


**SI Figure 6.** Oxidation and regeneration of SARS-CoV-2 $M^{pro}$ after removal of DTT. The corresponding c(s) distributions were obtained from sedimentation velocity analyses. Insets show $s_w$ binding isotherms, as calculated from the corresponding c(s) distributions. (**A**) $M^{pro}$ in non-reducing buffer (devoid of reductant DTT). Samples were diluted to 0.25 - 10 µM and analyzed by analytical ultracentrifugation (AUC) 23 h after dilution. (**B**) After overnight dialysis of $M^{pro}$ against non-reducing buffer, protein stock solution was diluted to 0.25 -10 µM with buffer + DTT. After 23 h of further incubation at room temperature, samples were analyzed by AUC. (**C**) After overnight dialysis against non-reducing buffer, $M^{Pro}$ was diluted with non-reducing buffer to 0.5 - 10 µM and incubated for 23 h at room temperature. Afterwards, 1 mM DTT was added and $M^{Pro}$ was allowed to regenerate for 23 h at room temperature before AUC analysis. (**D**) Comparison of $s_w$ isotherms obtained after $M^{pro}$ regeneration with DTT from experiments in (B) and (C) with those obtained for $M^{Pro}$ in buffer + DTT without previous oxidation (see Fig. 1B in the main manuscript) and in non-reducing buffer (A). $M^{Pro}$ oxidation in the absence of DTT is completely reversible, even if the protein was already dissociated before DTT addition. For better representation, all c(s) distributions in (A)-(C) were normalized to the same area.

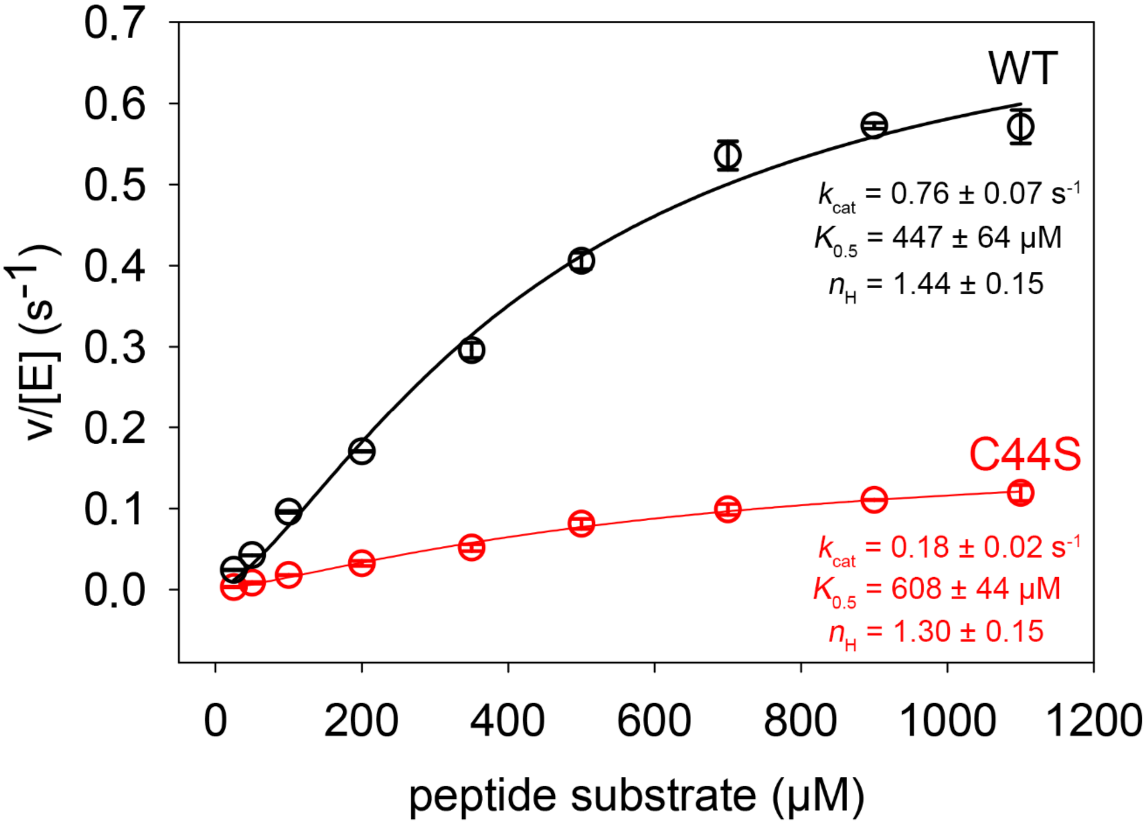


**SI Figure 7.** Steady-state kinetic analysis of $M^{pro}$ wild-type and variant C44S. The corresponding v_S plots for proteolytic cleavage of a peptidic model substrate are shown for the wild-type protein (WT) in black and for variant C44 in red. Experimental details are summarized in the Supplementary Methods. All measurements were carried out in duplicate and are shown as mean ± s.d. Data were fitted with the Hill equation to obtain estimates for the catalytic constant $k_{cat,}$ the substrate binding constant $K_{0.5}$ and the Hill coefficient $n_H$. The fits are shown as solid lines and the estimated kinetic constants are depicted for both proteins. Note that the C44S variant exhibits a ~4-fold decreased catalytic constant relative to the WT enzyme, while the substrate affinity is only slightly changed. Both the WT as well as variant C44S exhibit positive cooperativity as signified by Hill coefficients $n_H$ >1.

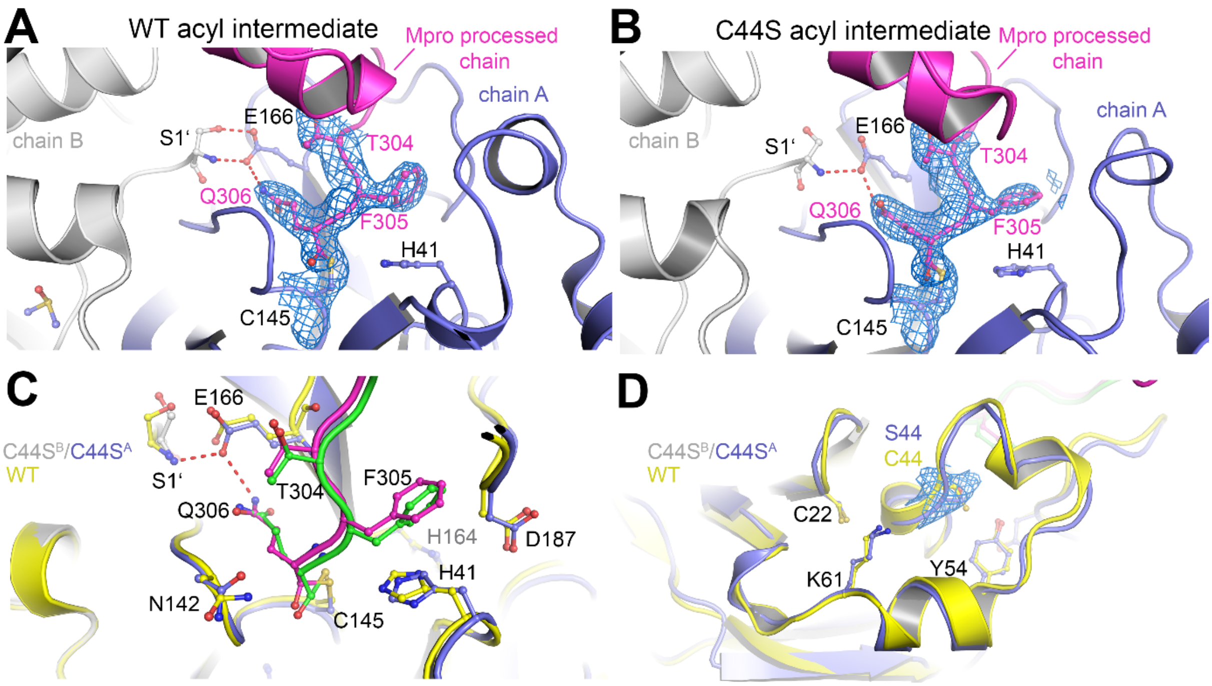


**SI Figure 8.** X-ray crystallographic structures of the covalent acyl intermediates in M^pro wild-type (WT) and variant C44S. (**A**) Snapshot of the acyl intermediate in M^pro WT (pdb code 7KHP) formed between a functional dimer in the asymmetric unit and a symmetry-related M^pro molecule. The two chains of the dimer are colored individually (chain A, blue; chain B, grey). Residues contributed by chain B are marked with an apostrophe. The C-terminus of the symmetry-related molecule, which visits the active site and forms a covalent linkage with catalytic C145, is highlighted in magenta. The structural model of C145 and the last three C-terminal residues of the symmetry-related molecule are superposed with the corresponding 2mFo-DFc electron density map contoured at 1σ. (**B**) Snapshot of the acyl intermediate in variant C44S (this study) formed between a functional dimer in the asymmetric unit and a symmetry-related M^pro molecule. The two chains of the dimer are colored individually (chain A, blue; chain B, grey). Residues contributed by chain B are marked with an apostrophe. The C-terminus of the symmetry-related molecule, which visits the active site and forms a covalent linkage with catalytic C145, is highlighted in magenta. The structural model of C145 and the last three C-terminal residues of the symmetry-related molecule are superposed with the corresponding 2mFo-DFc electron density map contoured at 1σ. (**C**) Superposition of the WT structure (yellow, symmetry-related molecule green) with that of variant C44 (blue/grey, symmetry-related molecule magenta) showing the active sites. Note the structural differences at the active site, in particular of H41, C145 and the visiting C-terminus. (**D**) Superposition of the WT structure (yellow) with that of variant C44 (blue), showing the allosteric SONOS redox switch sites. The structural model of mutation site S44 in the variant is superposed with the corresponding 2mFo-DFc electron density map.

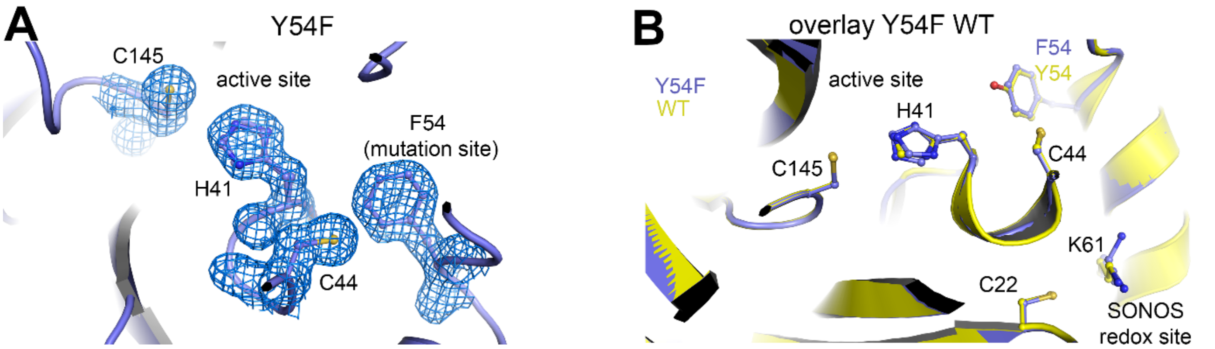


**SI Figure 9.** X-ray crystallographic structure of M$^{pro}$ variant Y54F. (**A**) Structure of the active site showing catalytic residues C145 and H41 as well as mutation site F54 and SONOS residue C44. The structural model is superposed with the corresponding 2mFo-DFc electron density map at a contour level of 1$\sigma$. (**B**) Superposition of the wild-type (WT) structure (yellow, pdb code 7KPH) with that of variant Y54F (blue, this study). Note the slight structural changes of catalytic residue H41.

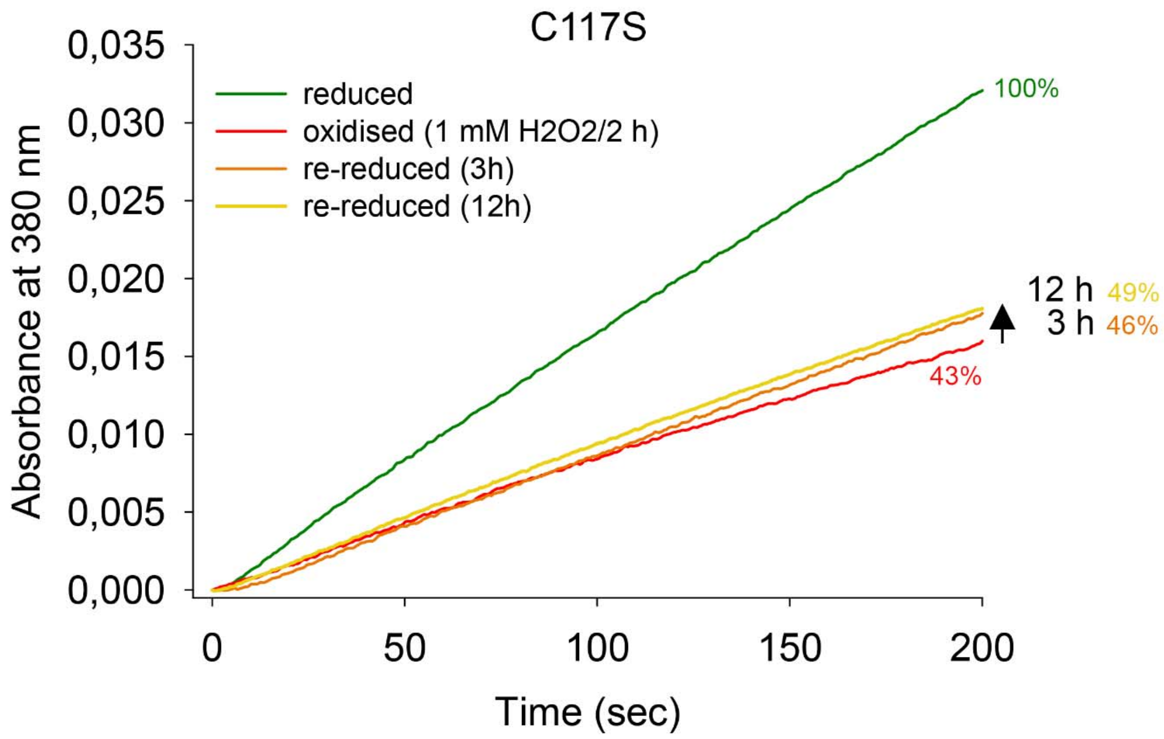


**SI Figure 10.** Redox-dependent activity of M$^{pro}$ variant C117S. Progress curves of M$^{pro}$-catalyzed substrate turnover in the reduced state, after oxidation with 1 mM $H_2O_2$ and after re-reduction with reductant DTT (3 h and 12 h). Experimental details are provided in the Supplementary Methods. An activity of 100% refers to that of the reduced enzyme. Please note that - in contrast to the wild-type protein - the loss of enzymatic activity after an oxidative insult with 1 mM $H_2O_2$ cannot be recovered.

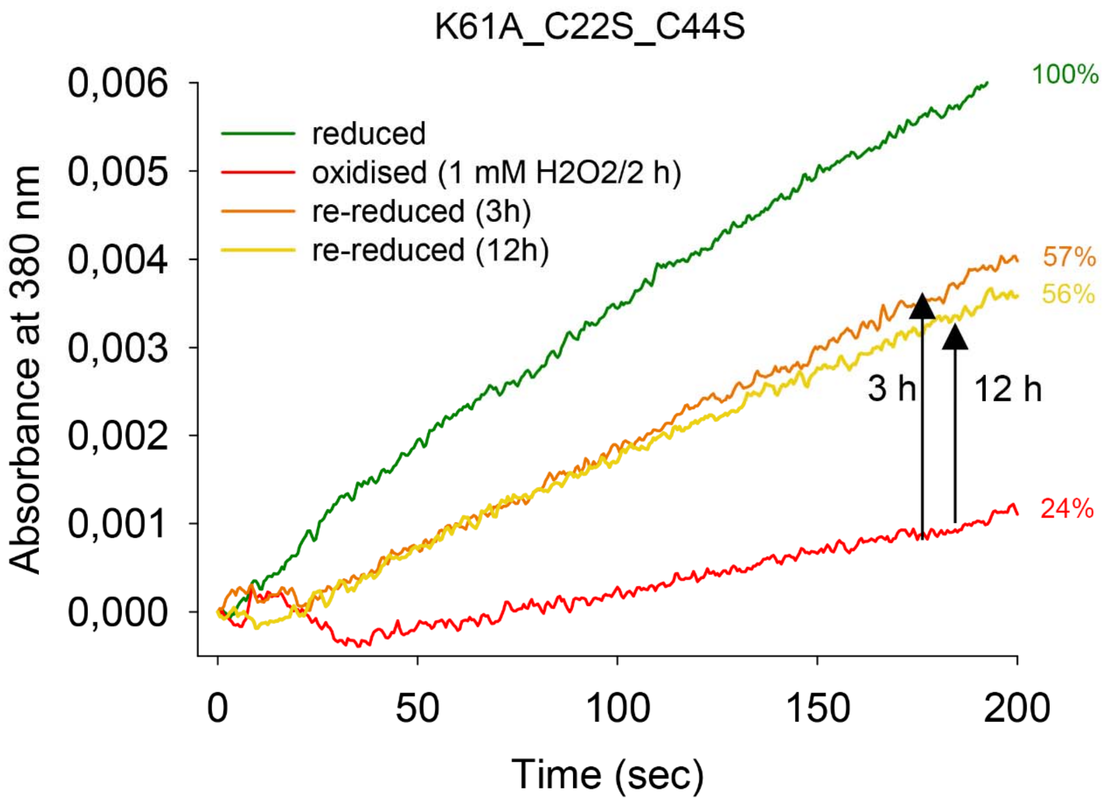


**SI Figure 11.** Redox-dependent activity of M$^{pro}$ triple variant K61A_C22S_C44S. Progress curves of M$^{pro}$-catalyzed substrate turnover in the reduced state, after oxidation with 1 mM $H_2O_2$ and after re-reduction with reductant DTT (3 h and 12 h). Experimental details are provided in the Supplementary Methods. An activity of 100% refers to that of the reduced enzyme. Please note that - in contrast to the wild-type protein - the loss of enzymatic activity after an oxidative insult with 1 mM $H_2O_2$ cannot be fully recovered.

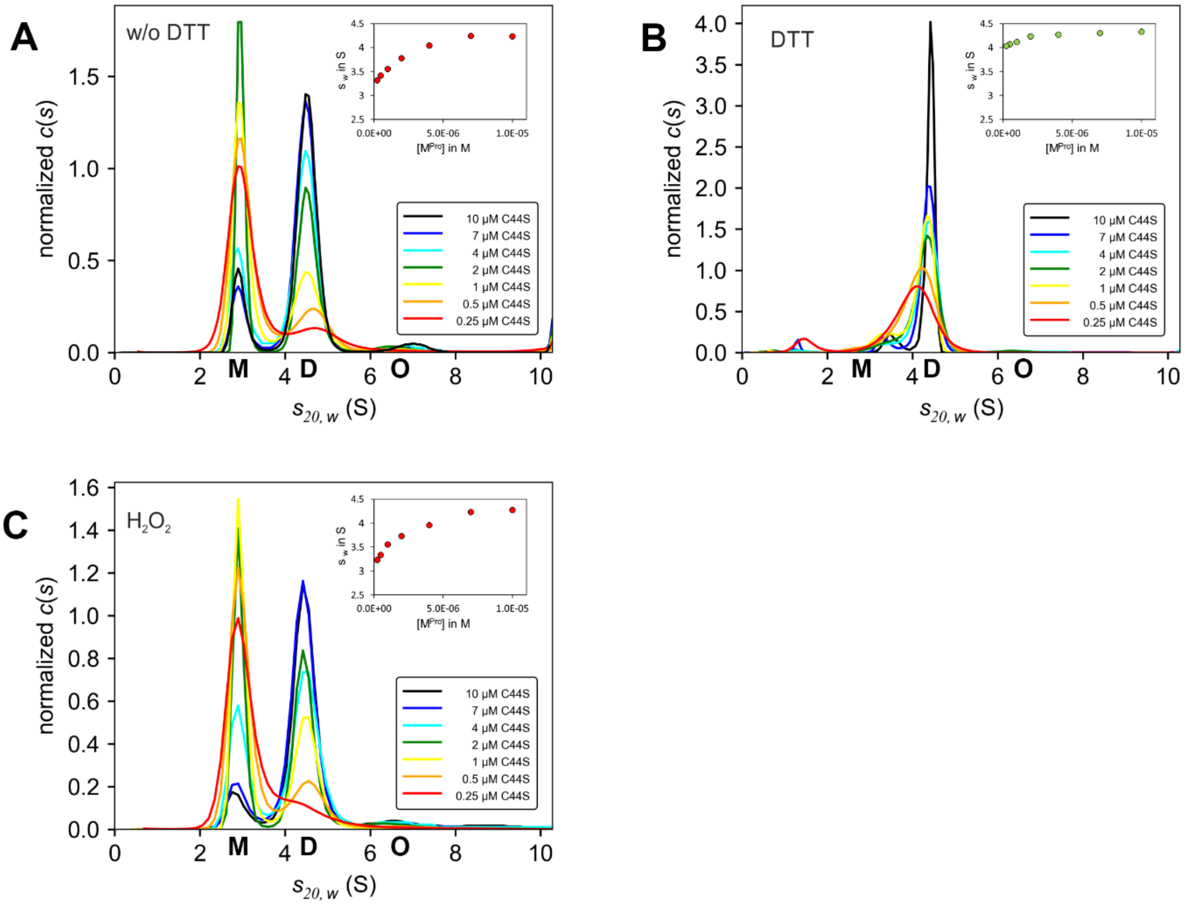


**SI Figure 12.** Redox-dependent sedimentation velocity analysis of SARS-CoV-2 $M^{pro}$ variant C44S. The oligomeric equilibrium was analyzed in a concentration range from 0.25 to 10 µM under either non-reducing (**A,** buffer devoid of DTT), reducing (**B**, buffer supplemented with 1 mM DTT) or oxidizing (**C**, 1 mM $H_2O_2$) conditions. The data indicate a redox-dependent monomer ⇔ dimer equilibrium with apparent equilibrium constants of $K_D^{app}$ <0.25 µM for the reduced enzyme and about 2.5 µM for the oxidized enzyme. Insets show $s_w$ binding isotherms, as calculated from the corresponding c(s) distributions. Abbreviations: M, monomer ($s_{20,w}$ = 2.9 S); D, dimer ($s_{20,w}$ = 4.5 S); O, oligomers ($s_{20,w} \approx$ 6.3 S).

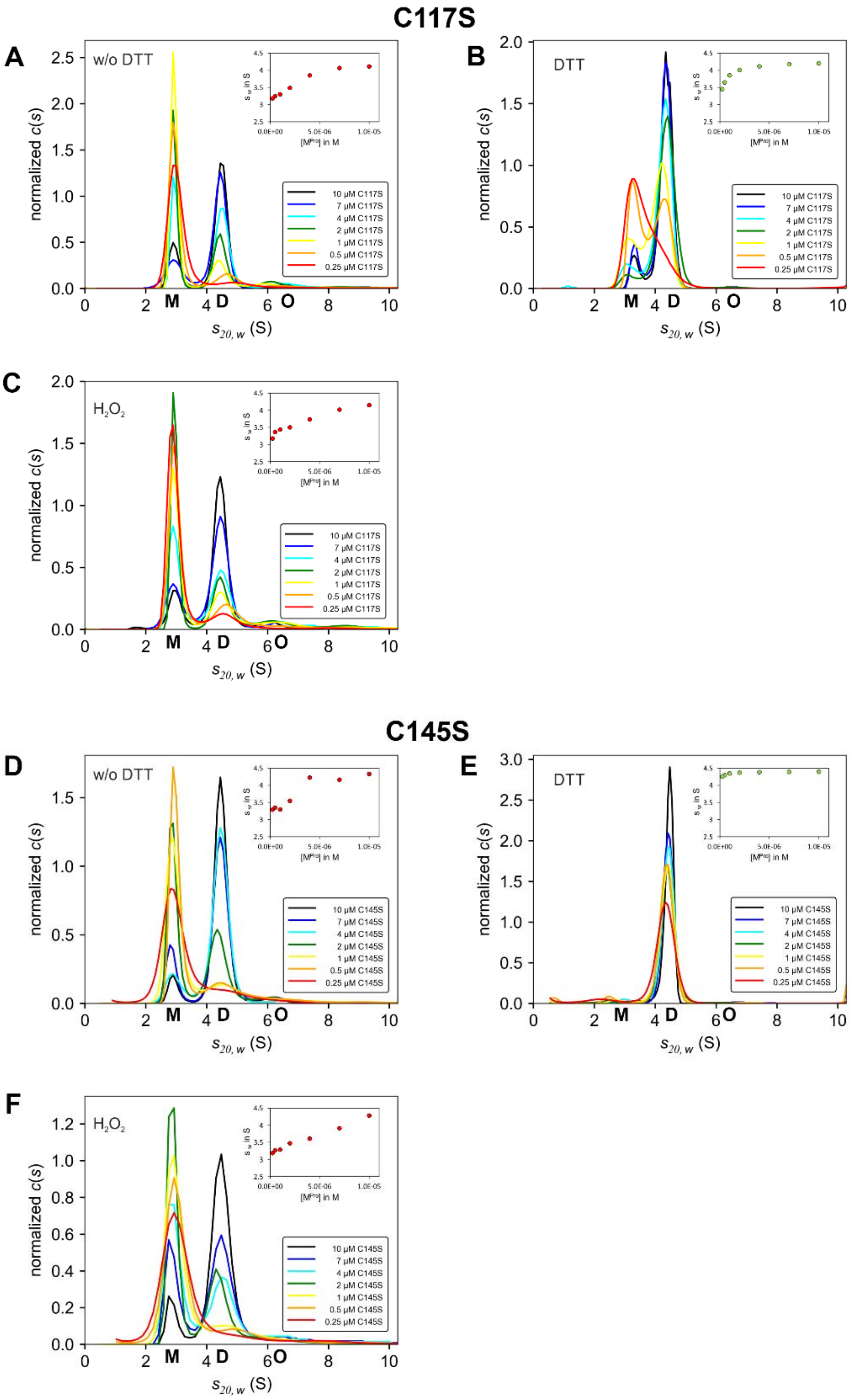


**SI Figure 13.** Redox-dependent sedimentation velocity analysis of SARS-CoV-2 $M^{pro}$ variants C117S (A-C) and C145S (D-F). The oligomeric equilibrium was analyzed in a concentration range from 0.25 to 10 µM under either non-reducing (**A/C,** buffer devoid of DTT), reducing (**B/D**, buffer supplemented with 1 mM DTT) or oxidizing (**C/F**, 1 mM $H_2O_2$) conditions. Insets show $s_w$ binding isotherms, as calculated from the corresponding c(s) distributions. As opposed to the wild-type enzyme and all other variants tested, variant C117S undergoes monomerization also under reducing conditions with an equilibrium constant $K_D^{app}$ in the range of 1 µM. For C145S, we observe a redox-dependent monomer ⇔ dimer equilibrium with an apparent equilibrium constant of $K_D^{app}$ <0.25 µM for the reduced enzyme. Under oxidizing conditions $K_D^{app}$ is in the lower micromolar range for both mutants. Abbreviations: M, monomer ($s_{20,w}$ = 2.9 S); D, dimer ($s_{20,w}$ = 4.5 S); O, oligomers ($s_{20,w}$ ≈ 6.3 S).

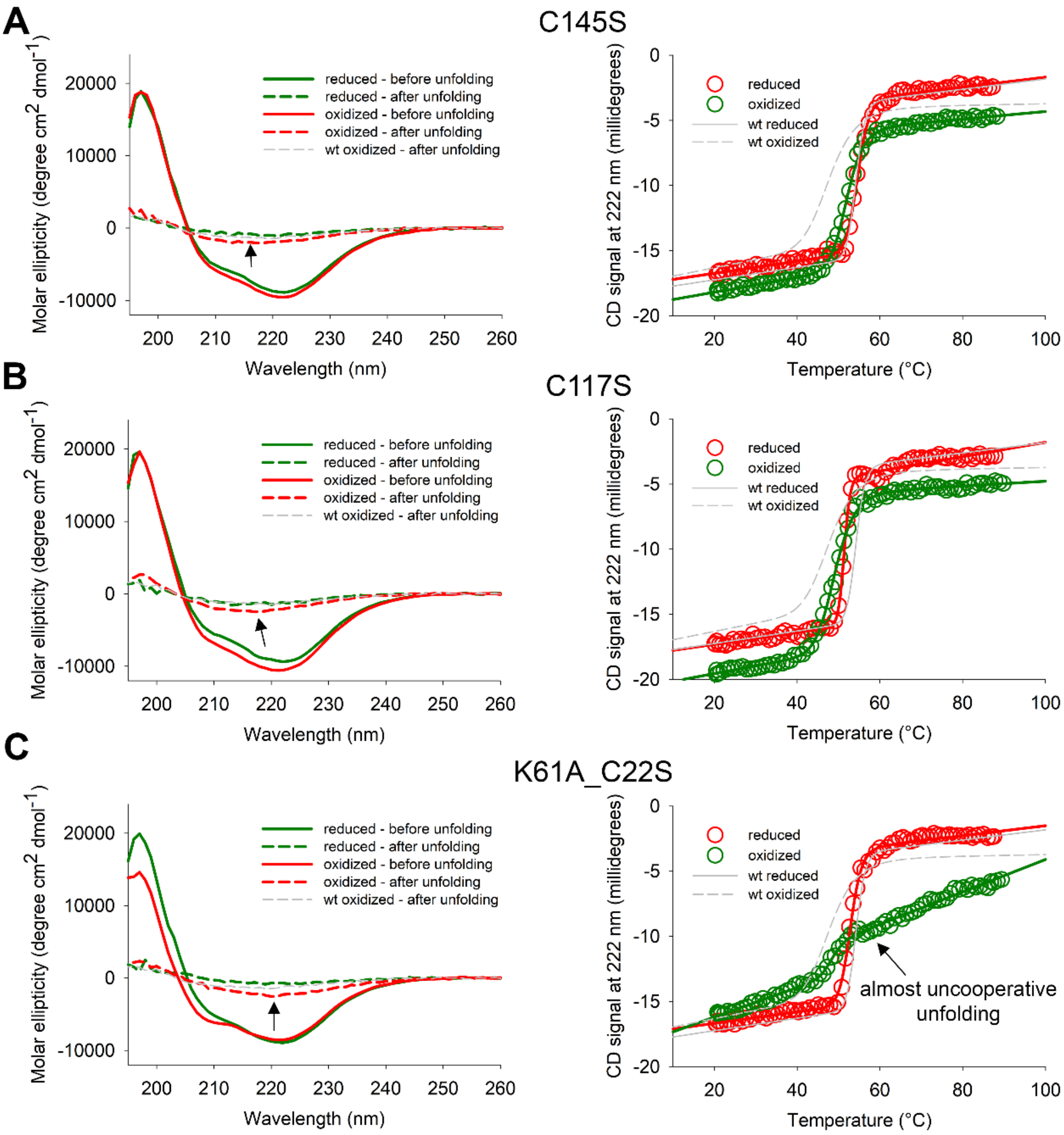


**SI Figure 14.** Secondary structure and thermal unfolding analysis of M$^{pro}$ variants C145S (**A**), C117S (**B**) and K61A_C22S (**C**) by far-UV CD spectroscopy under reducing and oxidizing conditions. The structural changes of both C145S and C117S upon oxidation are different compared to the wild-type protein (see Figure 1) and do not include an increase of ß-strand structural elements at the expense of $\alpha$-helices (Supplementary Table 5). Also, the melting temperatures (Supplementary Table 6) and cooperativity of unfolding are almost identical for both proteins under reducing and oxidizing conditions. Variant K61A_C22S shows the typical structural transition following an oxidation. Note, however, the atypical early onset of unfolding (below 40 °C) and decreased cooperativity (decreased steepness of transition) of unfolding of the oxidized enzyme in case of the K61A_C22S double variant that is implying a loosely associated structure and a high tendency to undergo aggregation. This is further supported by the detection of increased residual ß-sheet-like structures after thermal unfolding indicated by the arrows.

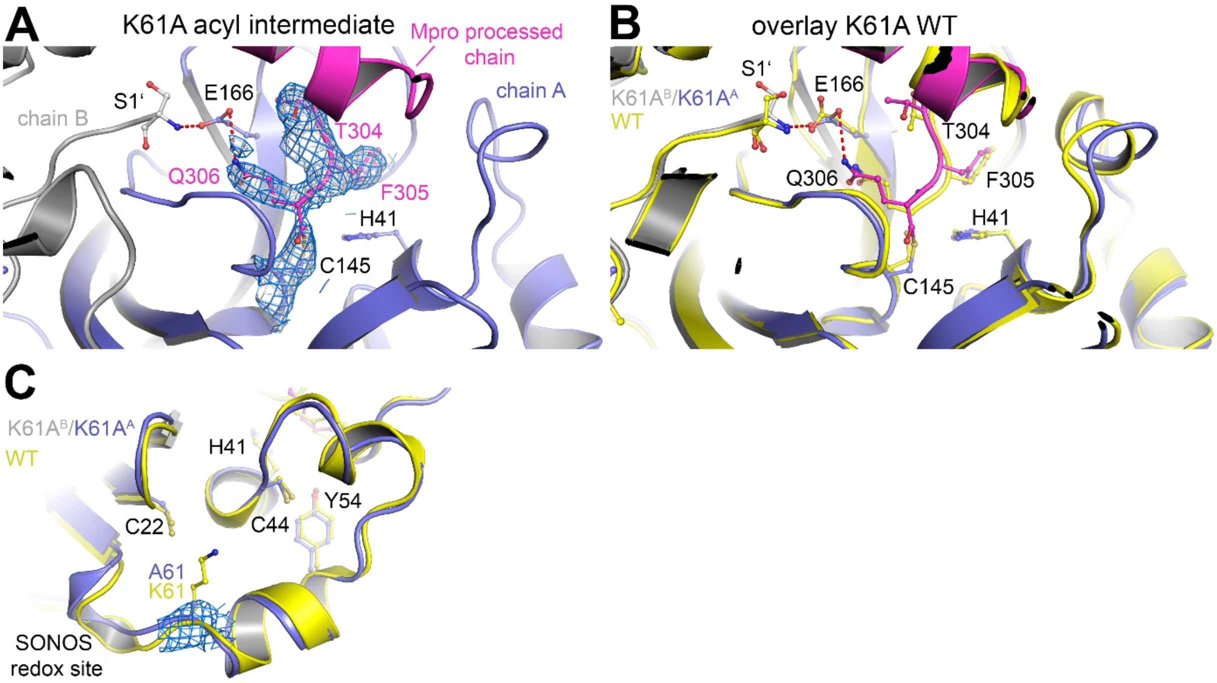


**SI Figure 15.** X-ray crystallographic structure of the covalent acyl intermediate in M$^{pro}$ variant K61A. (**A**) Snapshot of the acyl intermediate in M$^{pro}$ K61A (this study) formed between a functional dimer in the asymmetric unit and a symmetry-related M$^{pro}$ molecule. The two chains of the dimer are colored individually (chain A, blue; chain B, grey). Residues contributed by chain B are marked with an apostrophe. The C-terminus of the symmetry-related molecule, which visits the active site and forms a covalent linkage with catalytic C145, is highlighted in magenta. The structural model of C145 and the last three C-terminal residues of the symmetry-related molecule are superposed with the corresponding 2mFo-DFc electron density map contoured at 1$\sigma$. (**B**) Superposition of the WT structure (yellow) with that of variant K61A (blue/grey, symmetry-related molecule magenta) showing the active sites. Note the structural differences at the active site and at the dimer interface, in particular of C145 and E166. (**C**) Superposition of the wild-type (WT) structure (yellow) with that of variant K61A (blue), showing the allosteric SONOS redox switch sites. The structural model of mutation site A61 in the variant is superposed with the corresponding 2mFo-DFc electron density map.

Dimedone at C145

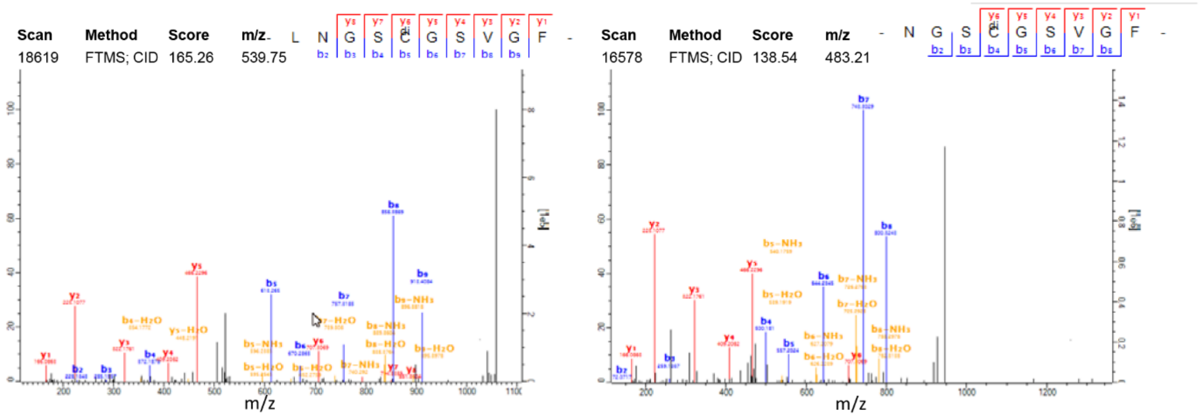


Dimedone at C156

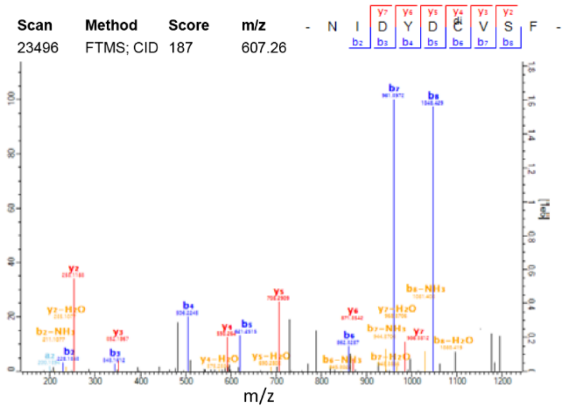


Dimedone at C300

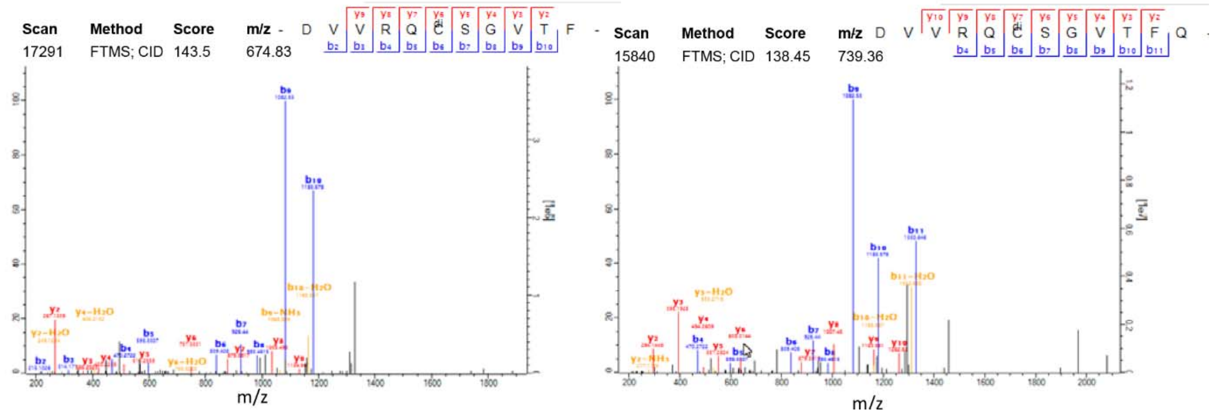


**SI Figure 16.** Representative mass spectra of dimedone containing peptides of SARS-CoV-2 $M^{pro}$. Dimedone (di) was detected as a modification of several peptides using tandem mass spectrometry upon treatment of $M^{pro}$ with 1 mM $H_2O_2$ for 10 minutes followed by incubation with 5 mM dimedone for 30 minutes. Recorded spectra suggest the existence of sulfenic acids at residues C145, C156 and C300.

#### Disulfide C117-C145

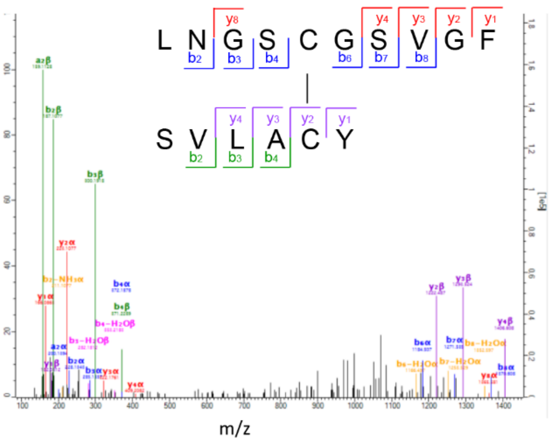


| Scan | Method | Score | PEP | Charge | m/z | Mass error [ppm] |
|---|---|---|---|---|---|---|
| 13084 | FTMS; HCD | 147.4 | 0.0225 | 2 | 796.86 | -0.024 |

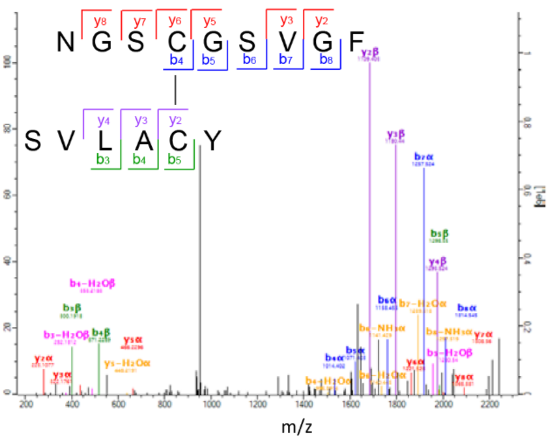


| Scan | Method | Score | PEP | Charge | m/z | Mass error [ppm] |
|---|---|---|---|---|---|---|
| 12129 | FTMS; CID | 121.93 | 0.0002 | 2 | 740.32 | -0.66 |

#### Disulfide C145-C156

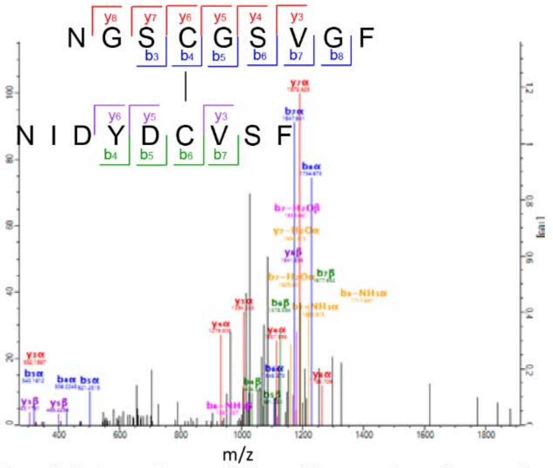


| Scan | Method | Score | PEP | Charge | m/z | Mass error [ppm] |
|---|---|---|---|---|---|---|
| 17022 | FTMS; CID | 89.6 | 0.0012 | 2 | 950.38 | 0.28 |

**SI Figure 17.** Representative mass spectra of disulfide containing peptides of SARS-CoV-2 M$^{pro}$. Disulfide linked peptides (C117-C145 and C145-C157) were found repetitively in M$^{pro}$ samples treated with 1 mM $H_2O_2$ and 20 mM $H_2O_2$ but not in M$^{pro}$ preincubated with 1 mM DTT using tandem mass spectrometry.

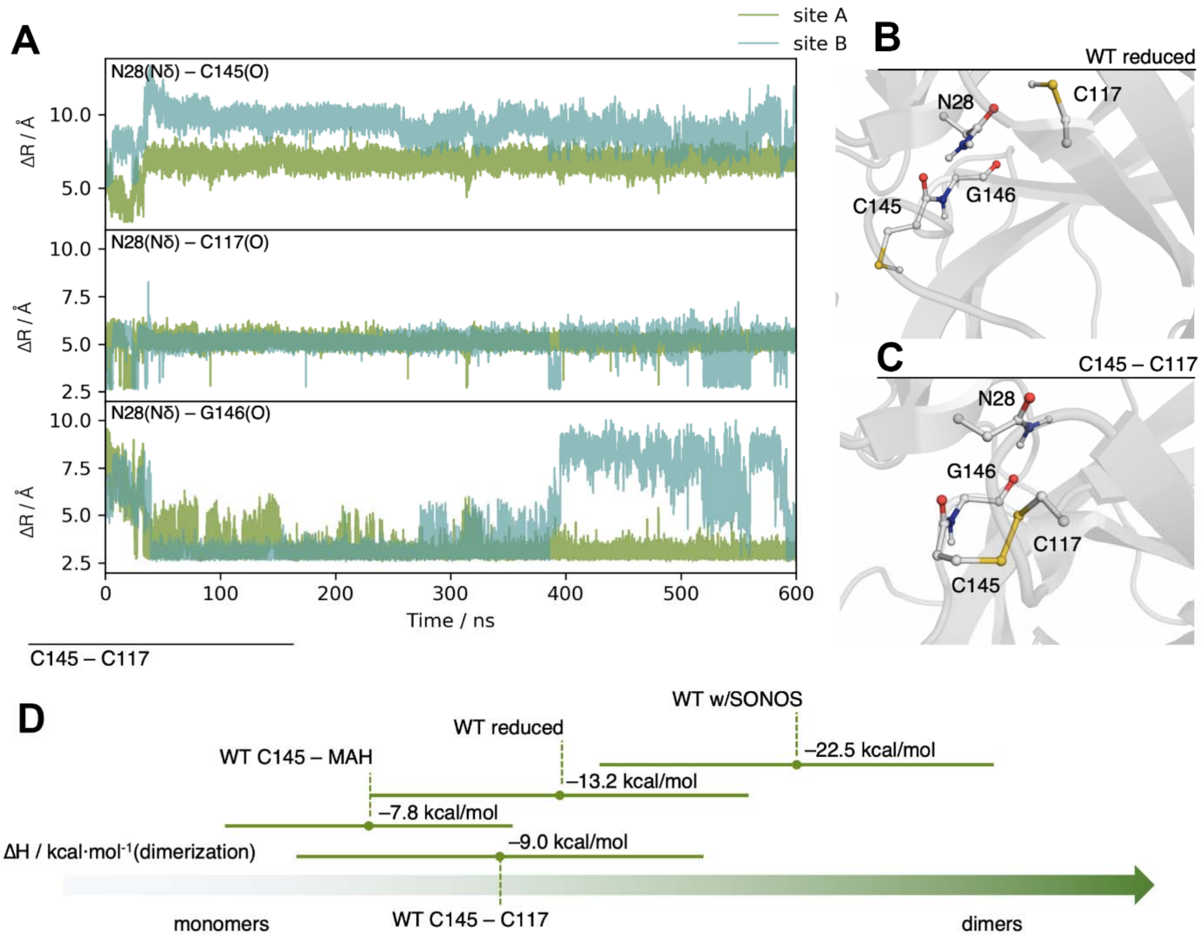


**SI Figure 18.** (**A**) Selected distances along the 600 ns MD trajectory of M$^{pro}$ with the disulfide bridge C145 - C117 formed on both monomers. After just a few nanoseconds, N28 is displaced from its interaction with the backbone carbonyls of C145 and C117 through the amide moiety. It builds an hydrogen bond to the carbonyl of G146, with occasional interactions to the C117. (**B,C**) Snapshots taken from MD trajectories illustrating the interactions of N28. In the reduced protein, N28 has stable interactions with the backbone atoms of C145 and C117 (**B**). Upon disulfide bridge formation (**C**), N28 is flipped and moves to interact with G146. (**D**) Summary of MMPBSA dimerization enthalpies for the different simulated variants of M$^{pro}$. Note the reduced dimerization enthalpies for M$^{pro}$ containg the C145-C117 disulfide and after covalent binding of MAH to C145.

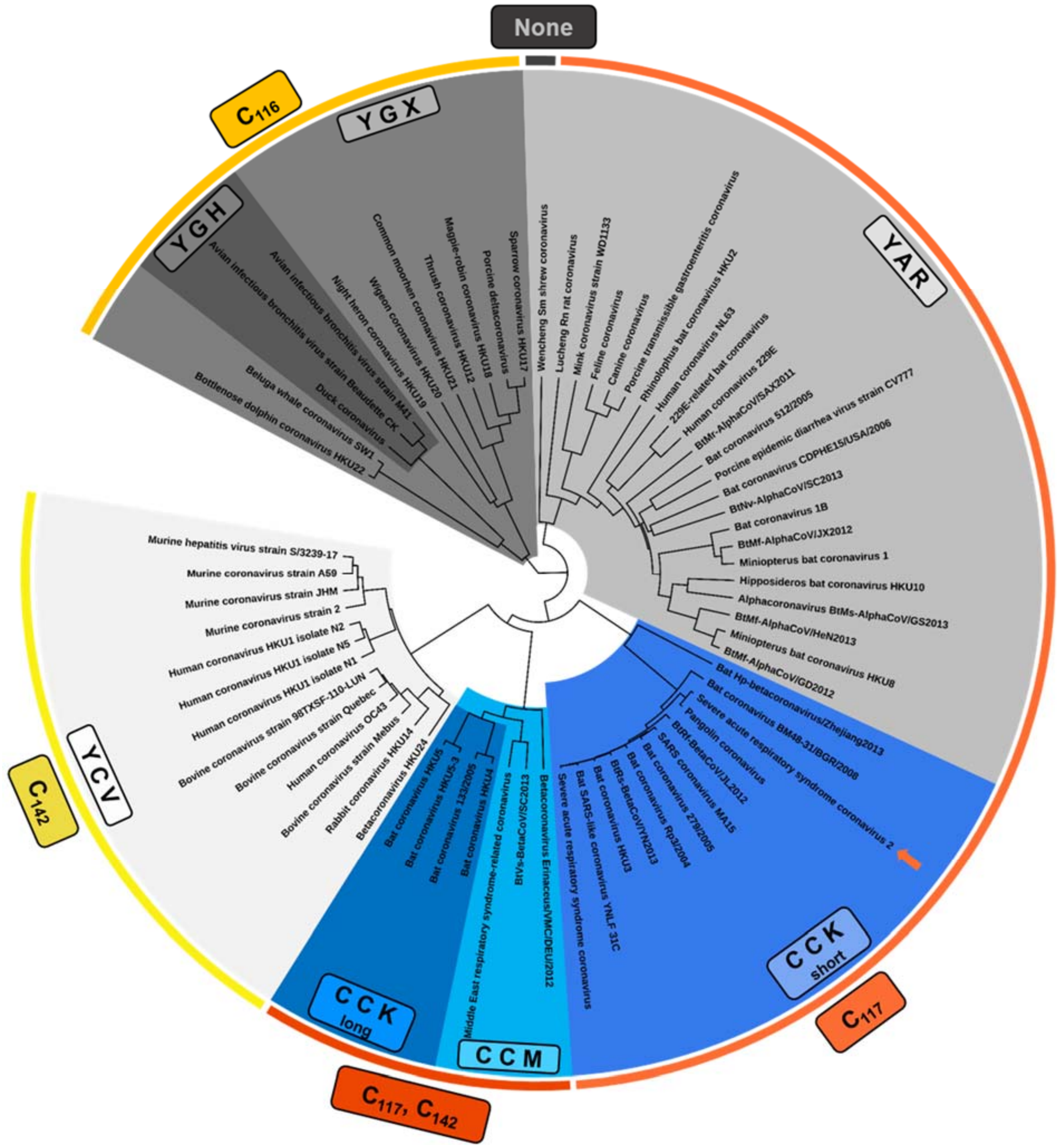

**SI Figure 19.** Tree-like representation of the dataset used for sequence conservation analyses in this work. Groups with similar SONOS triads are indicated by grey and blue hues, the respective amino acids at position 22, 44 and 61 are indicated (YCV, CCK, CCM, YAR, YGX, YGH). Groups with similar putative disulfide-forming partners for cysteine 145 are indicated by yellow and orange hues (C142, C117 or C142, C117, C116). The red arrow indicates the main protease from SARS-CoV-2.

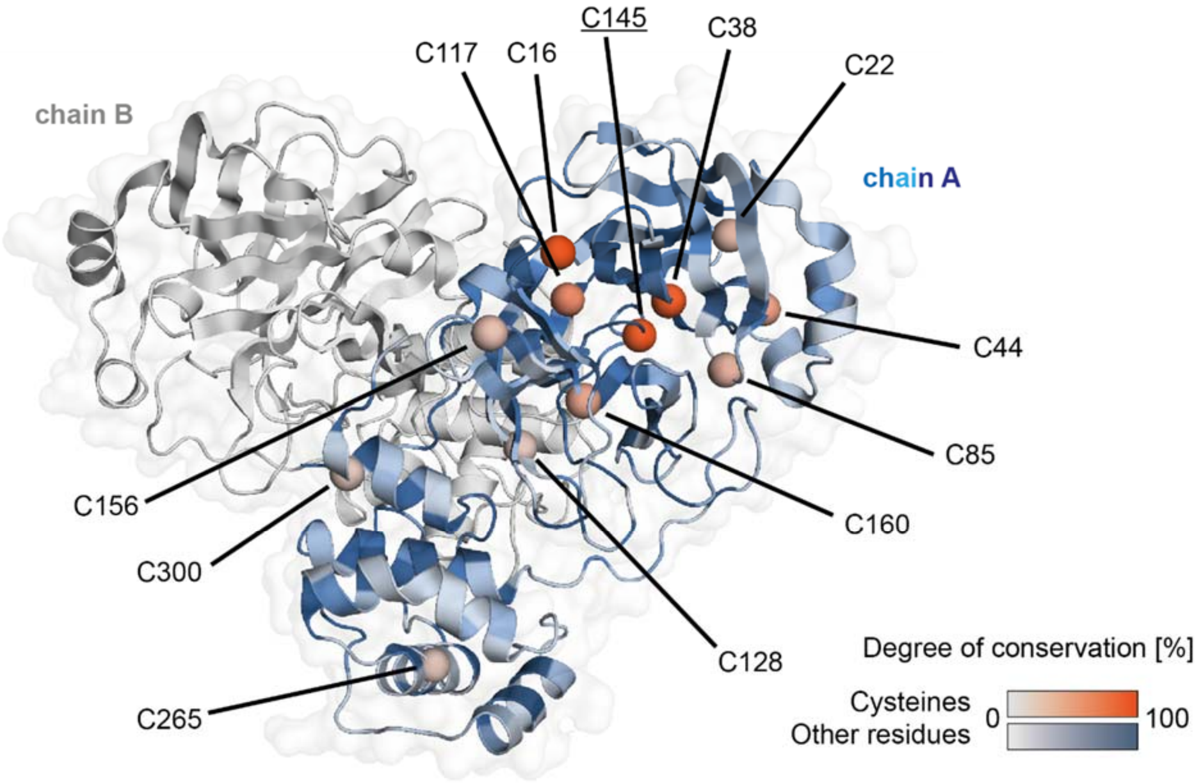


**SI Figure 20.** Mapping of sequence conservation among coronavirus main proteases onto the structure of the SARS-CoV-2 $M^{Pro}$. Overall conservation is shown in hues of blue, the conservation of the twelve cysteines in hues of red. The catalytic cysteine 145 is underscored.

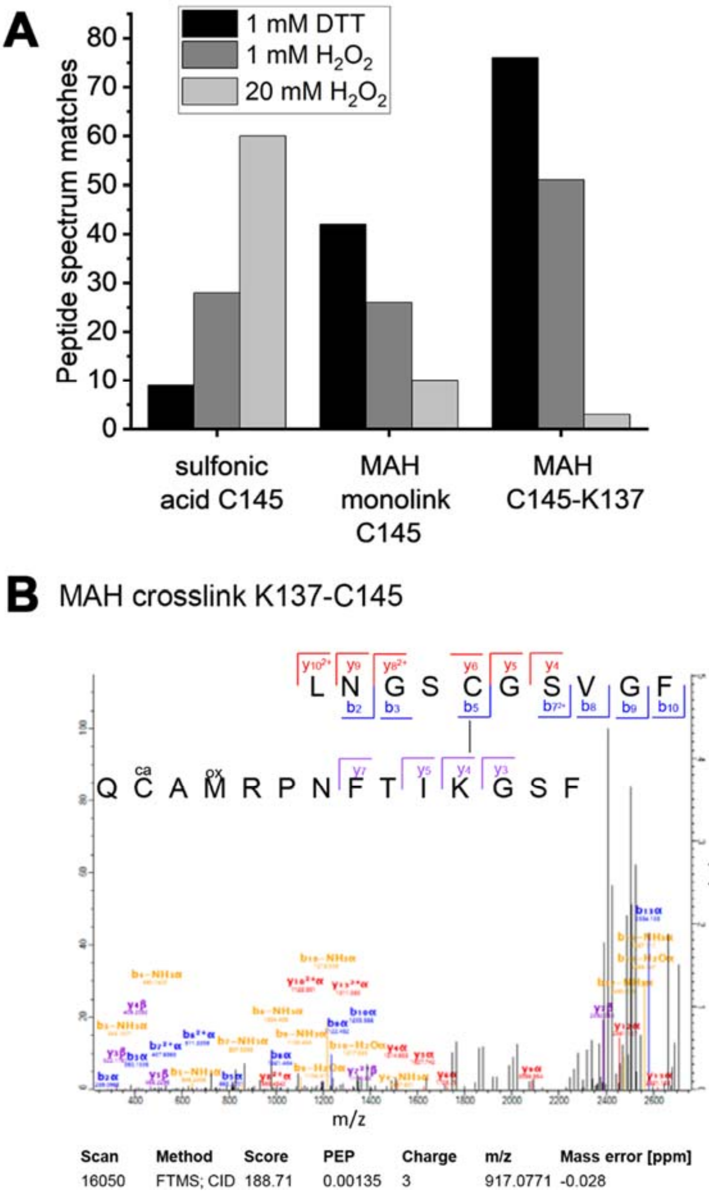


**SI Figure 21.** Mass spectrometric analysis of covalent modifications of $M^{pro}$ after reaction with heterobifunctional crosslinker maleimidoacetic acid N-hydroxysuccinimide ester (MAH) under different redox conditions. A reaction scheme for MAH crosslinking is shown in Figure 4C of the main manuscript. (**A)** Residues C145 (thiol) and K137 (amine) were identified as main reaction sites (exemplary spectrum in (**B**), ca: carbamidomethylation, ox: oxidation). With increasing $H_2O_2$ concentrations, the fraction of sulfonylated C145 becomes dominant based on the counts of peptide spectrum matches.

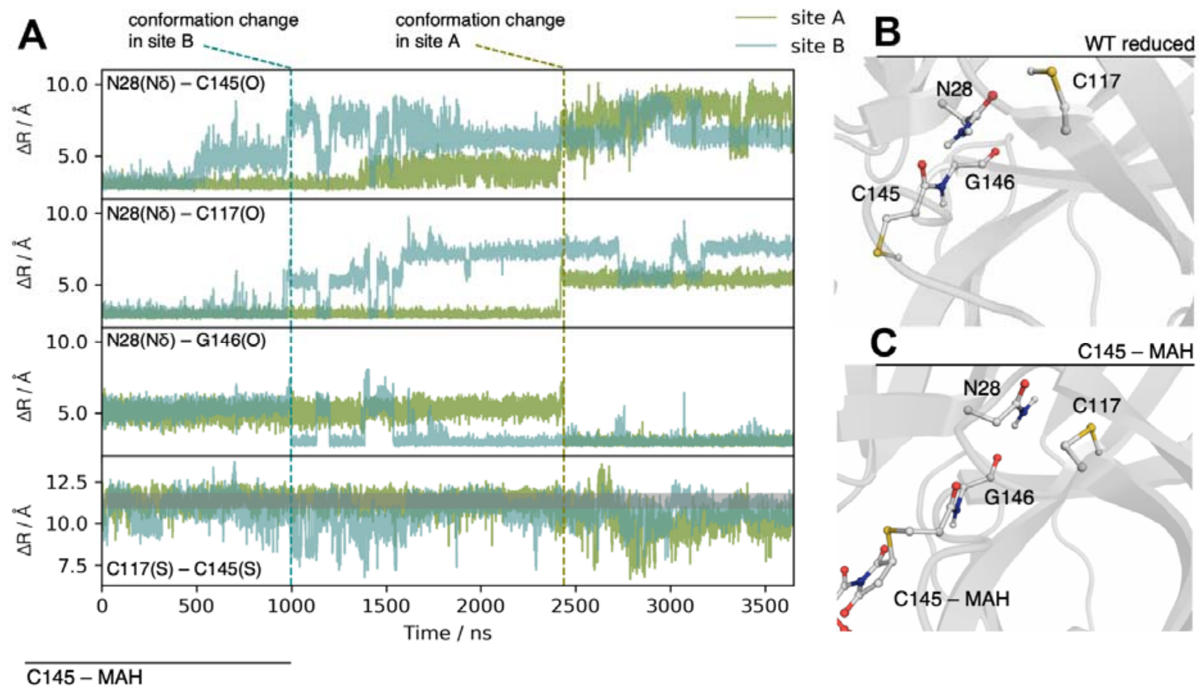


**SI Figure 22.** (**A**) Selected distances along the 3.65 µs MD trajectory of $M^{pro}$ with MAH covalently bound to C145. Long simulation times are required but a similar effect to the one observed in the disulfide C145 - C117 system takes place. After about 1 µs for the active site of chain B and 2.5 µs in chain A (approximately marked by the vertical dotted lines), the productive interactions of N28 with both cysteines are disrupted and N28 flips interacting with the backbone carbonyl of G146. After the conformation changes, C117 and C145 come in closer proximity in several events. The horizontal grey line depicts the average C117(S)-C145(S) distance in the reduced WT. (**B,C**) Snapshots taken from MD trajectories illustrating the interactions of N28. The N28 interaction with C117 and C145 is again illustrated for the reduced WT (**B**). In the case of the covalent bound C145 - MAH structure, a similar flip of N28 is observed as in the disulfide case (**C**).

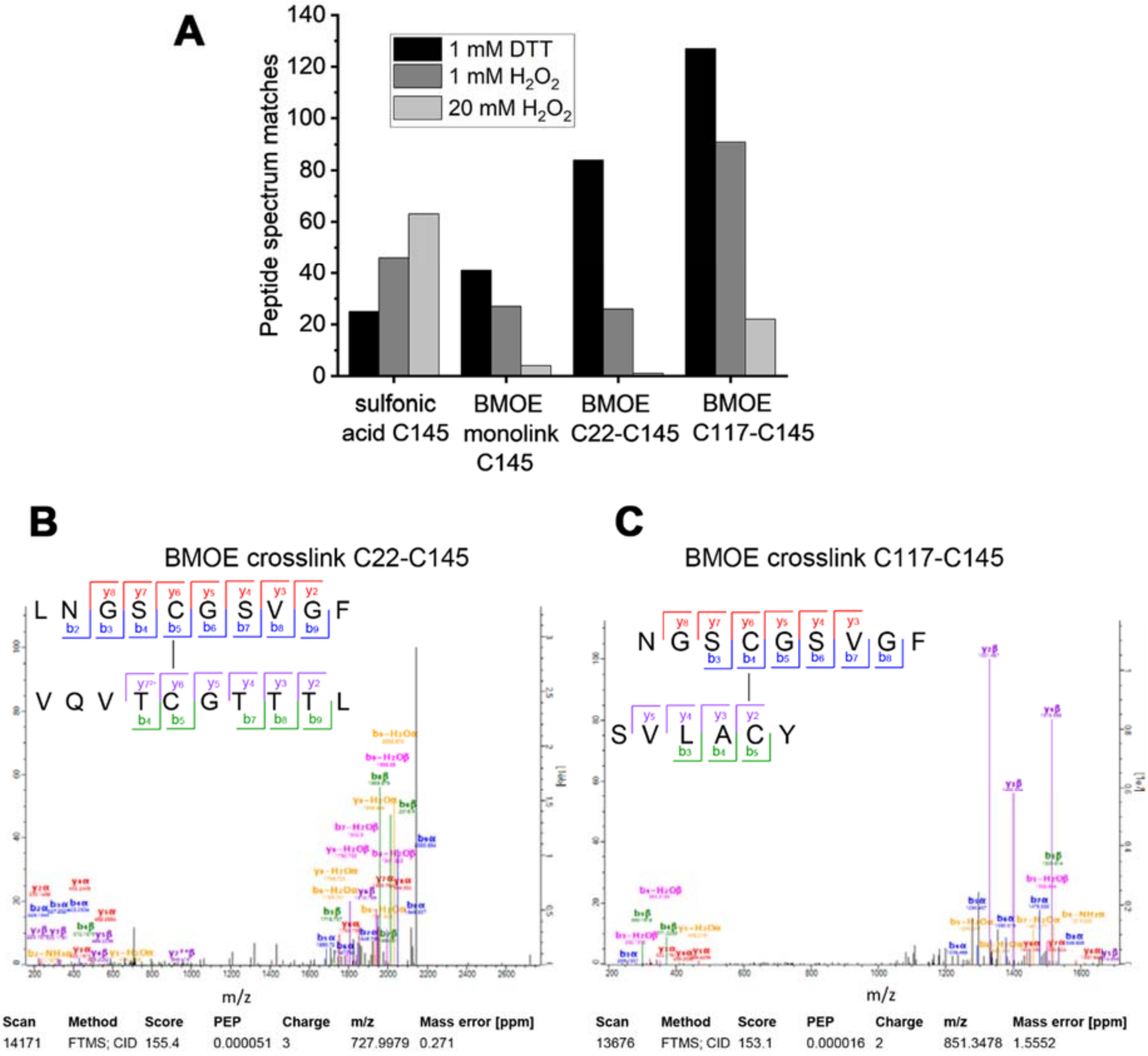


**SI Figure 23.** Mass spectrometric analysis of covalent modifications of M$^{pro}$ after reaction with homobifunctional crosslinker bismaleimidoethane (BMOE) under different redox conditions. (**A**) Residues C145 and C117 were identified as main crosslink sites (exemplary spectrum in **C**) as well as – to a lesser extent – C145 and C22 (exemplary spectrum in (**B**)). With increasing $H_2O_2$ concentrations, the fraction of sulfonylated C145 becomes dominant based on the counts of peptide spectrum matches.

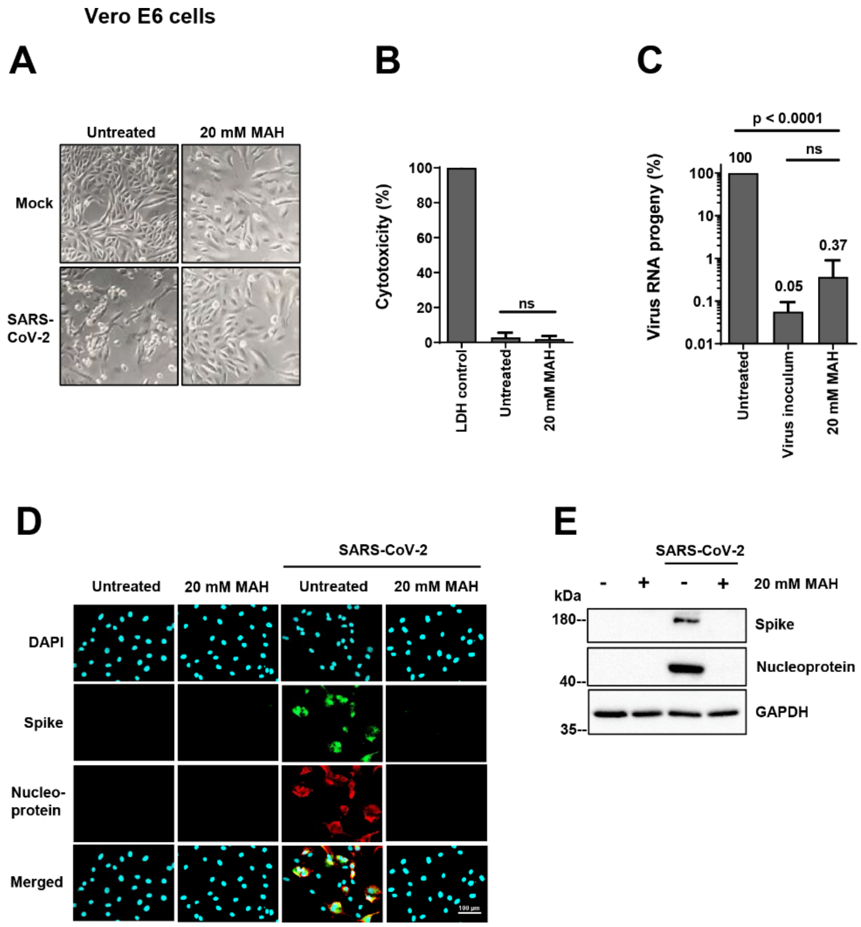


**SI Figure 24.** MAH inhibits SARS-CoV-2 propagation and the synthesis of viral proteins without detectable cytotoxicity. (**A**) Reduced cytopathic effect (CPE) upon treatment with MAH. Vero E6 cells were treated with 20 mM MAH or the PBS control for 1 h before infection, and then throughout the time of infection (48 h). Cell morphology was assessed by bright field microscopy. Note that the CPE was clearly visible in virus-infected cells but to a far lesser extent upon treatment with MAH. (**B**) Lack of measurable cytotoxicity by MAH. Vero E6 cells were treated with 20 mM MAH for 48 h. The release of lactate dehydrogenase (LDH) to the supernatant was quantified by bioluminescence as a read-out for cytotoxicity. The percentages reflect the proportion of LDH released to the media, compared to the overall amount of LDH in the cells (LDH control) (mean with SD, n=3). (**C**) Diminished virus RNA progeny by MAH. Vero E6 cells were treated and infected as described in A. RNA was isolated from the cell culture supernatant, and SARS-CoV-2 RNA was quantified by qRT-PCR. The amount of RNA found upon infection without drug treatment was defined as 100%, and the other RNA quantities were normalized accordingly. RNA was also isolated from the virus inoculum used to infect the cells. Note that MAH reduced SARS-CoV-2 RNA progeny more than 200-fold when compared to the untreated control (mean with SD, n = 3). **(D)** Representative images showing the reduction of viral protein synthesis by MAH. Vero E6 cells were treated and infected with SARS-CoV-2 as in A. Cell nuclei were stained with DAPI, and the SARS-CoV-2 Spike and Nucleoprotein were detected by immunofluorescence microscopy. (**E**) Reduced viral protein synthesis in the presence of MAH. Upon drug treatment and/or infection of Vero E6 cells, the viral Spike and Nucleoprotein as well as GAPDH (loading control) were detected by immunoblot analysis.

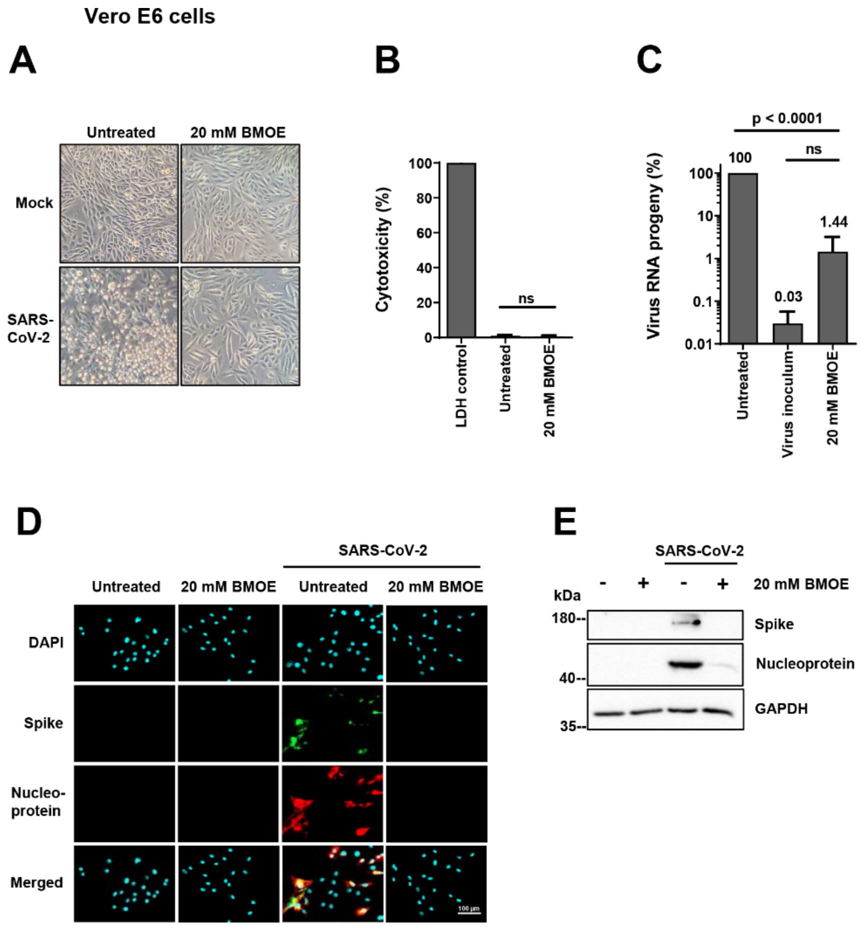


**SI Figure 25.** BMOE inhibits SARS-CoV-2 propagation and the synthesis of viral proteins without detectable cytotoxicity. (**A**) Reduced cytopathic effect (CPE) upon treatment with BMOE. Vero E6 cells were treated with 20 mM BMOE or the PBS control for 1 h before infection, and then throughout the time of infection (48 h). Cell morphology was assessed by bright field microscopy. Note that the CPE was clearly visible in virus-infected cells but to a far lesser extent upon treatment with BMOE. (**B**) Lack of measurable cytotoxicity by MAH and BMOE. Vero E6 cells were treated with 20 mM BMOE for 48 h. The release of lactate dehydrogenase (LDH) to the supernatant was quantified by bioluminescence as a read-out for cytotoxicity. The percentages reflect the proportion of LDH released to the media, compared to the overall amount of LDH in the cells (LDH control) (mean with SD, n=3). (**C**) Diminished virus RNA progeny by BMOE. Vero E6 cells were treated and infected as described in A. RNA was isolated from the cell culture supernatant, and SARS-CoV-2 RNA was quantified by qRT-PCR. The amount of RNA found upon infection without drug treatment was defined as 100%, and the other RNA quantities were normalized accordingly. RNA was also isolated from the virus inoculum used to infect the cells. Note that BMOE reduced SARS-CoV-2 RNA progeny more than 60-fold when compared to the untreated control (mean with SD, n = 3). **(D)** Representative images showing the reduction of viral protein synthesis by BMOE. Vero E6 cells were treated and infected with SARS-CoV-2 as in A. Cell nuclei were stained with DAPI, and the SARS-CoV-2 Spike and Nucleoprotein were detected by immunofluorescence microscopy. (**E**) Reduced viral protein synthesis in the presence of MAH. Upon drug treatment and/or infection of Vero E6 cells, the viral Spike and Nucleoprotein as well as GAPDH (loading control) were detected by immunoblot analysis.

### Supplementary Information

### Tables

**Supplementary Table 1.** Redox-dependent enzymatic activities of $M^{pro}$ wild-type and variants.

| $M^{pro}$ Cysteine Variants | Reduced [1] Relative activity (%) | | | Oxidized [2] Relative activity (%) | | | Reactivation (200 s) [3] Relative activity (%) | | | change (%) |
|---|---|---|---|---|---|---|---|---|---|---|
| WT | 100.0 [4] | ± | 2.6 | 30.5 | ± | 1.0 | 34.0 | ± | 1.0 | +3.5% |
| C16S | 75.3 | ± | 1.1 | 25.4 | ± | 0.1 | 27.1 | ± | 0.2 | +1.7% |
| C22S | 84.4 | ± | 1.2 | 26.3 | ± | 0.2 | 31.2 | ± | 0.8 | +4.9% |
| C38S | 92.1 | ± | 0.5 | 31.8 | ± | 0.3 | 36.4 | ± | 0.4 | +4.6% |
| C44S | 16.4 | ± | 0.1 | 7.3 | ± | 0.3 | 7.6 | ± | 0.4 | +0.3% |
| C85S | 71.2 | ± | 1.7 | 26.4 | ± | 0.8 | 29.5 | ± | 0.1 | +3.1% |
| C117S | 41.9 | ± | 0.6 | 16.7 | ± | 0.1 | 14.2 | ± | 1.1 | -2.5% |
| C128S | 74.3 | ± | 0.6 | 22.5 | ± | 0.6 | 28.4 | ± | 1.1 | +5.9% |
| C145S | n. a.[5] | | | n. a. | | | n. a. | | | |
| C156S | 77.5 | ± | 1.8 | 28.7 | ± | 0.7 | 35.0 | ± | 0.9 | +6.3% |
| C160S | 92.7 | ± | 3.0 | 28.2 | ± | 0.5 | 30.1 | ± | 0.1 | +1.9% |
| C265S | 85.1 | ± | 2.3 | 30.7 | ± | 0.4 | 34.4 | ± | 1.9 | +3.7% |
| C300S | 92.7 | ± | 0.3 | 32.1 | ± | 0.4 | 35.9 | ± | 2.0 | +3.8% |

| $M^{pro}$ SONOS Variants | Reduced Relative activity (%) | | | Oxidized Relative activity (%) | | | Reactivation (200 s) Relative activity (%) | | | change (%) |
|---|---|---|---|---|---|---|---|---|---|---|
| WT | 100.0 | ± | 2.6 | 30.5 | ± | 1.0 | 34.0 | ± | 1.0 | +3.5 |
| C22S | 84.4 | ± | 1.2 | 26.3 | ± | 0.2 | 31.2 | ± | 0.8 | +4.9 |
| C44S | 16.4 | ± | 0.1 | 7.3 | ± | 0.3 | 7.6 | ± | 0.4 | +0.3 |
| C44A | 29.4 | ± | 0.4 | 12.4 | ± | 0.5 | 11.1 | ± | 0.4 | -1.3 |
| C22S_C44S | 10.4 | ± | 0.4 | 4.0 | ± | 0.0 | 2.3 | ± | 0.2 | -1.7 |
| K61A | 55.2 | ± | 2.0 | 20.2 | ± | 2.4 | 18.7 | ± | 2.1 | -1.5 |
| K61A_C22S | 42.4 | ± | 1.9 | 16.6 | ± | 2.2 | 17.2 | ± | 0.7 | +0.6 |
| K61A_C44S | 10.2 | ± | 0.3 | 3.2 | ± | 0.0 | 3.5 | ± | 0.1 | +0.3 |
| K61A_C22S_C44S | 7.0 | ± | 1.4 | 3.0 | ± | 0.1 | 2.0 | ± | 0.3 | -1.0 |
| Y54F | 19.4 | ± | 0.0 | 7.1 | ± | 1.8 | 8.2 | ± | 0.5 | +1.1 |

[1] in presence of 1 mM DTT

[2] after oxidation with 1 mM $H_2O_2$ for 2 h on ice

[3] Reactivation was initiated by addition of 20 mM DTT to the oxidized enzyme and kinetically analyzed 200 s after reduction

[4] 100% activity refers to the activity of wild-type $M^{pro}$ under reducing conditions. For clarity, relative activities are visualized in color-coded fashion (green: high activities, yellow: medium activities, red: low activities)

[5] no measurable activity

Experimental details are provided in the Supplementary Methods.

**Supplementary Table 2.** X-ray data collection and refinement statistics.

| | C44S acyl intermediate | K61A acyl intermediate | Y54F Resting state |
|---|---|---|---|
| **Data collection** | | | |
| Wavelength [Å] | 0.827 | 0.827 | 0.976 |
| Space group | C2 | C2 | C2 |
| Asymmetric unit | functional dimer | functional dimer | monomer |
| *Cell dimensions* | | | |
| a,b,c [Å] | 122.89, 81.84, 63.56 | 122.48, 81.29, 63.70 | 113.66, 53.49, 44.59 |
| α, β, γ [deg] | 90.00, 90.04, 90.00 | 90.00, 90.22, 90.00 | 90.00, 102.03, 90.00 |
| Resolution range [Å] | 67.837 – 2.120 (2.352 – 2.120)* | 46.451 – 2.488 (2.691 – 2.477) | 55.629 – 1.634 (1.762 – 1.634) |
| $R_{meas}$ | 0.128 (1.675) | 0.171 (1.734) | 0.139 (2.695) |
| $R_{pim}$ | 0.052 (0.66) | 0.064 (0.646) | 0.034 (0.665) |
| <I/σ(I)> | 7.6 (1.4) | 6.7 (1.2) | 12.8 (1.4) |
| Ellipsoidal completeness [%] | 91.9 (61.7) | 84.1 (32.4) | 92.2 (51.7) |
| Redundancy | 7.1 (7.4) | 7.2 (6.7) | 16.8 (16.2) |
| **Refinement** | | | |
| Resolution | 46.39 – 2.12 | 46.451 – 2.488 | 55.63 – 1.63 |
| Number of reflections | 22608 | 16753 | 23931 |
| $R_{work}/R_{free}$ [%] | 26.01/29.64 | 25.8/33.6 | 17.4/21.4 |
| *No. atoms* | | | |
| Protein | 4753 | 4726 | 2531 |
| Solvent | 73 | 36 | 237 |
| *B-factors* | | | |
| Protein | 73.35 | 100.09 | 23.31 |
| Solvent | 54.22 | 54.55 | 32.91 |
| *RMSD* | | | |
| Bond lengths [Å] | 0.003 | 0.001 | 0.004 |
| Bond angles | 0.532 | 0.394 | 0.753 |
| *Ramachandran statistics* | | | |
| Favored | 91.45 | 87.99 | 98.03 |
| Allowed | 8.55 | 9.87 | 1.64 |
| Outliers | 0.00 | 2.14 | 0.33 |
| *Diffraction limits (Å) and corresponding principal axes of the ellipsoid fitted to the diffraction cut-off surface as direction cosines in the orthogonal basis (standard PDB convention), and in terms of reciprocal unit-cell vectors:* | | | |
| #1 | 2.845 (0.925, 0.000, -0.389) 0.622 a* - 0.208 c* | 2.763 (0.974, 0.000, -0.225) 0.993 a* - 0.121 c* | 2.058 (0.932, 0.000, 0.270) a* + 0.026 c* |
| #2 | 2.345 (0.000, 1.000, 0.000) b* | 2.558 (0.000, 1.000, 0.000) b* | 1.67 (0.000, 1.000, 0.000) b* |
| #3 | 2.065 (0.380, 0.000, 0.925) 0.622 a* + 0.783 c* | 2.390 (0.225, 0.000, 0.974) 0.406 a* + 0.914 c* | 1.586 (-0.270, 0.000, 0.963) -0.567 a* + 0.823 c* |

* The values in parentheses refer to the highest resolution shell.

**Supplementary Table 3.** Redox-dependent distribution of $M^{pro}$ oligomeric species for cysteine variants as analyzed by gel filtration experiments in color-coded representation [1].

| Quaternary structure | $M^{pro}$ Cysteine Variants | Untreated | Reduced | Oxidized 1h | Oxidized 5h | Re-Reduced |
|---|---|---|---|---|---|---|
| **Oligomer** | WT | 0.4% | 0.4% | 0.7% | 0.9% | 0.3% |
| | C16S | 0.4% | 0.4% | 0.9% | 9.0% | 0.4% |
| | C22S | 0.5% | 0.5% | 0.9% | 6.1% | 0.5% |
| | C38S | 0.4% | 0.4% | 0.8% | 5.5% | 0.4% |
| | C44S | 0.3% | 0.4% | 0.5% | 5.3% | 0.2% |
| | C85S | 0.4% | 0.4% | 0.5% | 4.7% | 0.3% |
| | C117S | 0.1% | 0.2% | 0.1% | 1.8% | 0.5% |
| | C128S | 0.4% | 0.2% | 1.1% | 6.9% | 0.6% |
| | C145S | 0.9% | 0.8% | 3.4% | 4.1% | 2.0% |
| | C156S | 1.7% | 0.9% | 1.8% | 4.8% | 1.2% |
| | C160S | 0.6% | 0.5% | 1.5% | 11.2% | 0.4% |
| | C265S | 0.2% | 0.1% | 0.5% | 3.4% | 0.0% |
| | C300S | 0.5% | 0.6% | 0.5% | 3.2% | 0.4% |
| **Dimer** | WT | 83.6% | 83.5% | 71.7% | 60.0% | 84.5% |
| | C16S | 63.4% | 63.3% | 49.6% | 33.4% | 61.8% |
| | C22S | 83.0% | 82.3% | 73.3% | 54.1% | 82.6% |
| | C38S | 76.7% | 76.4% | 66.7% | 45.1% | 76.1% |
| | C44S | 79.5% | 79.0% | 62.0% | 39.6% | 78.9% |
| | C85S | 76.3% | 76.1% | 62.1% | 44.0% | 75.1% |
| | C117S | 36.5% | 36.5% | 46.5% | 54.1% | 42.3% |
| | C128S | 72.1% | 72.6% | 52.0% | 35.4% | 71.3% |
| | C145S | 93.5% | 91.4% | 89.6% | 88.0% | 89.5% |
| | C156S | 81.6% | 82.3% | 72.3% | 54.5% | 81.7% |
| | C160S | 79.4% | 79.5% | 68.7% | 52.8% | 78.6% |
| | C265S | 78.2% | 77.7% | 64.6% | 46.1% | 77.7% |
| | C300S | 70.6% | 70.2% | 54.0% | 27.3% | 69.1% |
| **Monomer** | WT | 15.9% | 16.1% | 27.6% | 39.1% | 15.2% |
| | C16S | 36.3% | 36.3% | 49.5% | 57.7% | 37.8% |
| | C22S | 16.5% | 17.2% | 25.8% | 39.8% | 16.9% |
| | C38S | 22.9% | 23.2% | 32.5% | 49.4% | 23.5% |
| | C44S | 20.3% | 20.6% | 37.6% | 55.1% | 20.9% |
| | C85S | 23.3% | 23.5% | 37.4% | 51.4% | 24.6% |
| | C117S | 63.3% | 63.3% | 53.4% | 44.1% | 57.3% |
| | C128S | 27.5% | 27.2% | 46.9% | 57.7% | 28.1% |
| | C145S | 5.6% | 7.8% | 7.0% | 7.9% | 8.5% |
| | C156S | 16.7% | 16.7% | 26.0% | 40.7% | 17.1% |
| | C160S | 20.0% | 20.0% | 29.7% | 36.0% | 21.0% |
| | C265S | 21.6% | 22.1% | 34.9% | 50.5% | 22.3% |
| | C300S | 28.9% | 29.2% | 45.4% | 69.5% | 30.5% |

[1] For clarity, relative fractions of each oligomeric state are visualized in color-coded fashion (green: high amount, yellow: medium amount, red: low amount).

**Supplementary Table 4.** Redox-dependent distribution of $M^{pro}$ oligomeric species for SONOS variants as analyzed by gel filtration experiments in color-coded representation [1].

| Quaternary structure | $M^{pro}$ SONOS Variants | Untreated | Reduced | Oxidized 1h | Oxidized 5h | Re-Reduced |
|---|---|---|---|---|---|---|
| **Oligomer** | WT | 0.44% | 0.36% | 0.70% | 0.90% | 0.30% |
| | C22S | 0.48% | 0.48% | 0.91% | 6.08% | 0.51% |
| | C44S | 0.25% | 0.36% | 0.45% | 5.34% | 0.21% |
| | C44A | 0.39% | 0.51% | 0.84% | 3.52% | 0.64% |
| | C22S_C44S | 0.15% | 0.25% | 2.02% | 8.58% | 0.32% |
| | K61A | 0.58% | 0.52% | 1.04% | 4.95% | 0.72% |
| | K61A_C22S | 0.41% | 0.41% | 0.60% | 6.23% | 0.37% |
| | K61A_C44S | 0.21% | 0.16% | 0.38% | 1.01% | 0.22% |
| | K61A_C22S_C44S | 0.10% | 0.16% | 0.27% | 8.26% | 0.28% |
| | Y54F | 0.26% | 0.31% | 0.78% | 7.62% | 0.33% |
| **Dimer** | WT | 83.61% | 83.51% | 71.74% | 60.00% | 84.51% |
| | C22S | 83.01% | 82.33% | 73.30% | 54.12% | 82.56% |
| | C44S | 79.49% | 79.04% | 61.95% | 39.59% | 78.88% |
| | C44A | 83.05% | 82.54% | 71.46% | 56.22% | 82.72% |
| | C22S_C44S | 77.02% | 77.13% | 57.51% | 36.93% | 76.16% |
| | K61A | 79.47% | 79.55% | 67.14% | 53.10% | 79.28% |
| | K61A_C22S | 78.37% | 78.19% | 65.95% | 45.01% | 77.57% |
| | K61A_C44S | 77.47% | 77.41% | 60.70% | 38.49% | 76.80% |
| | K61A_C22S_C44S | 79.80% | 79.04% | 57.86% | 34.64% | 77.57% |
| | Y54F | 78.56% | 78.69% | 63.30% | 45.73% | 78.17% |
| **Monomer** | WT | 15.94% | 16.13% | 27.56% | 39.10% | 15.18% |
| | C22S | 16.51% | 17.19% | 25.79% | 39.80% | 16.93% |
| | C44S | 20.25% | 20.60% | 37.60% | 55.07% | 20.91% |
| | C44A | 16.57% | 16.95% | 27.70% | 40.27% | 16.65% |
| | C22S_C44S | 22.83% | 22.62% | 40.47% | 54.49% | 23.53% |
| | K61A | 19.95% | 19.93% | 31.83% | 41.95% | 20.00% |
| | K61A_C22S | 21.23% | 21.41% | 33.44% | 48.77% | 22.06% |
| | K61A_C44S | 22.32% | 22.43% | 38.91% | 60.50% | 22.98% |
| | K61A_C22S_C44S | 20.10% | 20.80% | 41.86% | 57.09% | 22.15% |
| | Y54F | 21.18% | 21.00% | 35.92% | 46.64% | 21.50% |

[1] For clarity, relative fractions of each oligomeric state are visualized in color-coded fashion (green: high amount, yellow: medium amount, red: low amount).

**Supplementary Table 5.** Redox-dependent secondary structure contents for $M^{pro}$ wild-type and variants as analyzed by CD spectroscopy in color-coded representation [1].

| CD spectra 195-260 nm | α-Helix | | | | |
|---|---|---|---|---|---|
| | Reduced | Oxidized | Δ Oxidized vs. Reduced | Δ Reduced rel. to WT | Δ Oxidized rel. to WT |
| WT | 28.80% | 27.70% | -1.10% | / | / |
| C22S | 30.40% | 28.20% | -2.20% | 1.60% | 0.50% |
| C44S | 28.00% | 27.20% | -0.80% | -0.80% | -0.50% |
| C117S | 29.30% | 30.90% | 1.60% | 0.50% | 3.20% |
| C128S | 28.80% | 27.80% | -1.00% | 0.00% | 0.10% |
| C145S | 24.60% | 26.60% | 2.00% | -4.20% | -1.10% |
| C156S | 27.90% | 27.90% | 0.00% | -0.90% | 0.20% |
| C300S | 27.70% | 27.90% | 0.20% | -1.10% | 0.20% |
| K61A | 28.50% | 27.30% | -1.20% | -0.30% | -0.40% |
| K61A_C22S | 28.80% | 27.60% | -1.20% | 0.00% | -0.10% |
| K61A_C44S | 29.10% | 27.40% | -1.70% | 0.30% | -0.30% |
| K61A_C22S_C44S | 27.20% | 29.30% | 2.10% | -1.60% | 1.60% |
| C22S_C44S | 28.30% | 26.80% | -1.50% | -0.50% | -0.90% |
| Y54F | 28.50% | 27.90% | -0.60% | -0.30% | 0.20% |

| CD spectra 195-260 nm | Antiparallel ß-strands | | | | |
|---|---|---|---|---|---|
| | Reduced | Oxidized | Δ Oxidized to vs. Reduced | Δ Reduced rel. to WT | Δ Oxidized rel. to WT |
| WT | 9.40% | 10.90% | 1.50% | | |
| C22S | 8.60% | 10.80% | 2.20% | -0.80% | -0.10% |
| C44S | 10.10% | 11.60% | 1.50% | 0.70% | 0.70% |
| C117S | 9.40% | 8.70% | -0.70% | 0.00% | -2.20% |
| C128S | 9.70% | 10.70% | 1.00% | 0.30% | -0.20% |
| C145S | 14.40% | 11.60% | -2.80% | 5.00% | 0.70% |
| C156S | 10.30% | 10.80% | 0.50% | 0.90% | -0.10% |
| C300S | 10.40% | 10.60% | 0.20% | 1.00% | -0.30% |
| K61A | 9.60% | 11.50% | 1.90% | 0.20% | 0.60% |
| K61A_C22S | 9.60% | 11.40% | 1.80% | 0.20% | 0.50% |
| K61A_C44S | 9.50% | 11.60% | 2.10% | 0.10% | 0.70% |
| K61A_C22S_C44S | 10.80% | 10.10% | -0.70% | 1.40% | -0.80% |
| C22S_C44S | 10.10% | 12.30% | 2.20% | 0.70% | 1.40% |
| Y54F | 9.80% | 11.10% | 1.30% | 0.40% | 0.20% |

[1] For clarity, relative fractions of α-helices (upper table) or antiparallel ß-strands (lower table) are visualized in color-coded fashion (green: high amount, yellow: medium amount, red: low amount).

**Supplementary Table 6.** Redox-dependent melting temperatures of M$^{pro}$ wild-type and variants as analyzed by CD spectroscopy in color-coded representation [1].

| **Thermal Unfolding** | Reduced $T_m$ | | | Oxidized $T_m$ | | | $\Delta T_m$ Oxidized vs. Reduced | | | $\Delta T_m$ Reduced Relative to WT | | | $\Delta T_m$ Oxidized Relative to WT | | |
|---|---|---|---|---|---|---|---|---|---|---|---|---|---|---|---|
| WT | 54.22 | ± | 0.06 | 48.05 | ± | 0.17 | -6.17 | ± | 0.23 | / | | | / | | |
| **SONOS variants** | | | | | | | | | | | | | | | |
| C22S | 52.17 | ± | 0.06 | 45.80 | ± | 0.13 | -6.36 | ± | 0.19 | -2.05 | ± | 0.12 | -2.25 | ± | 0.30 |
| C44S | 52.55 | ± | 0.06 | 46.78 | ± | 0.15 | -5.77 | ± | 0.21 | -1.67 | ± | 0.12 | -1.27 | ± | 0.32 |
| C22S_C44S | 51.08 | ± | 0.07 | 45.43 | ± | 0.19 | -5.65 | ± | 0.26 | -3.14 | ± | 0.13 | -2.62 | ± | 0.36 |
| Y54F | 53.23 | ± | 0.07 | 46.49 | ± | 0.12 | -6.74 | ± | 0.19 | -0.98 | ± | 0.13 | -1.56 | ± | 0.29 |
| K61A | 53.70 | ± | 0.10 | 48.86 | ± | 0.16 | -4.84 | ± | 0.26 | -0.52 | ± | 0.16 | 0.81 | ± | 0.33 |
| K61A_C22S | 52.87 | ± | 0.08 | 48.34 | ± | 0.32 | -4.53 | ± | 0.41 | -1.35 | ± | 0.15 | 0.29 | ± | 0.49 |
| K61A_C44S | 53.15 | ± | 0.09 | 49.79 | ± | 0.26 | -3.36 | ± | 0.35 | -1.06 | ± | 0.16 | 1.74 | ± | 0.42 |
| K61A_C22S_C44S | 51.70 | ± | 0.08 | 50.06 | ± | 0.18 | -1.64 | ± | 0.26 | -2.52 | ± | 0.15 | 2.01 | ± | 0.34 |
| **Cysteine variants** | | | | | | | | | | | | | | | |
| C117S | 51.39 | ± | 0.06 | 49.20 | ± | 0.12 | -2.19 | ± | 0.18 | -2.83 | ± | 0.13 | 1.15 | ± | 0.29 |
| C128S | 50.65 | ± | 0.12 | 48.16 | ± | 0.10 | -2.49 | ± | 0.22 | -3.57 | ± | 0.18 | 0.10 | ± | 0.27 |
| C145S | 54.64 | ± | 0.08 | 52.00 | ± | 0.08 | -2.64 | ± | 0.15 | 0.42 | ± | 0.14 | 3.95 | ± | 0.24 |
| C156S | 53.99 | ± | 0.07 | 48.84 | ± | 0.17 | -5.14 | ± | 0.24 | -0.23 | ± | 0.13 | 0.79 | ± | 0.34 |
| C300S | 53.24 | ± | 0.07 | 49.79 | ± | 0.23 | -3.45 | ± | 0.29 | -0.98 | ± | 0.13 | 1.74 | ± | 0.39 |

[1] For clarity, melting temperatures for the reduced and oxidized proteins are visualized in color-coded fashion (green: high temperature, yellow: medium temperature, red: low temperature). An independent scaling was used for the differences between oxidized and reduced proteins.
