## Appendix 1 - Gel filtration experiments for "Redox regulation of the SARS-CoV-2 main protease provides new opportunities for drug design"

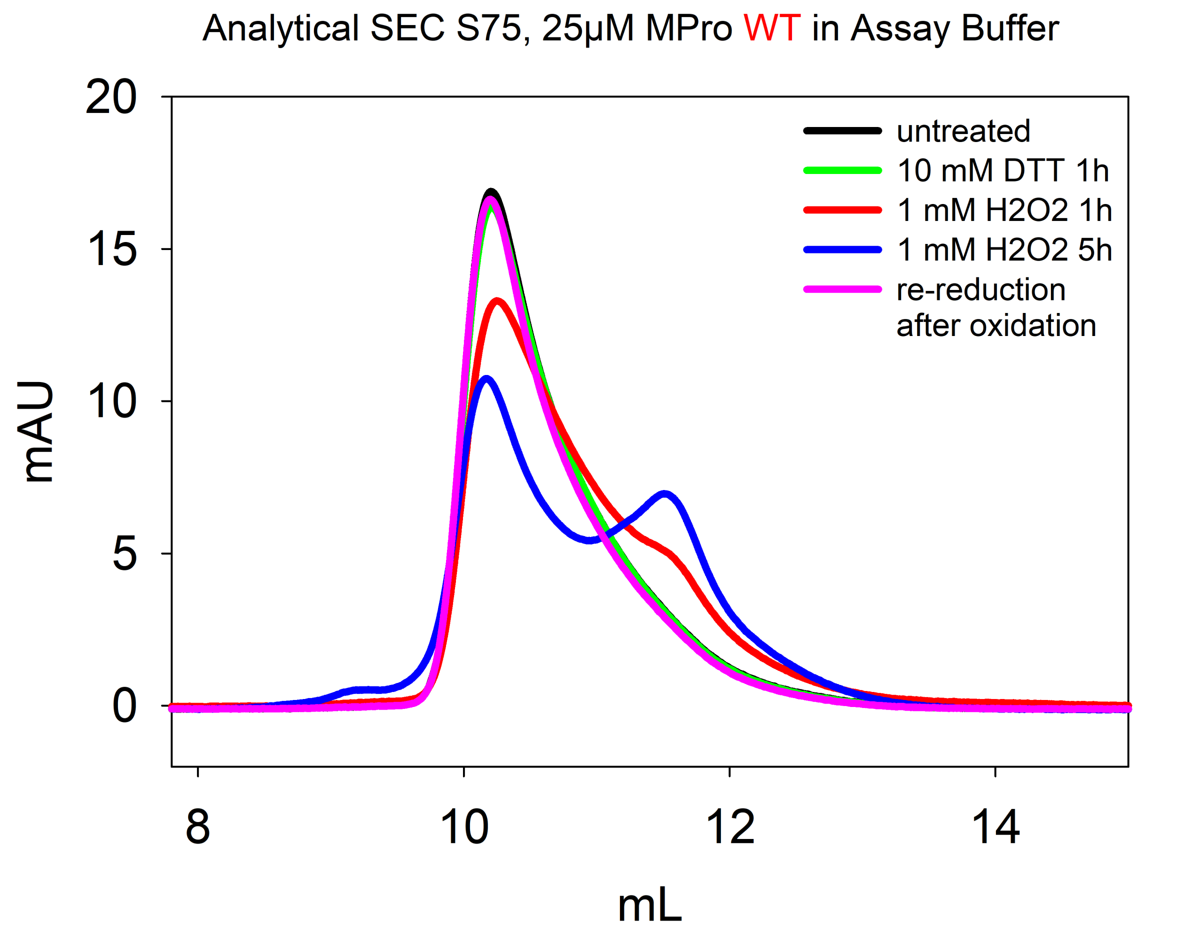
Analytical SEC S75, 25µM MPro WT in Assay Buffer
20
15
10
5
0
mAU
8
10
12
14
mL
untreated
10 mM DTT 1h
1 mM H2O2 1h
1 mM H2O2 5h
re-reduction
after oxidation

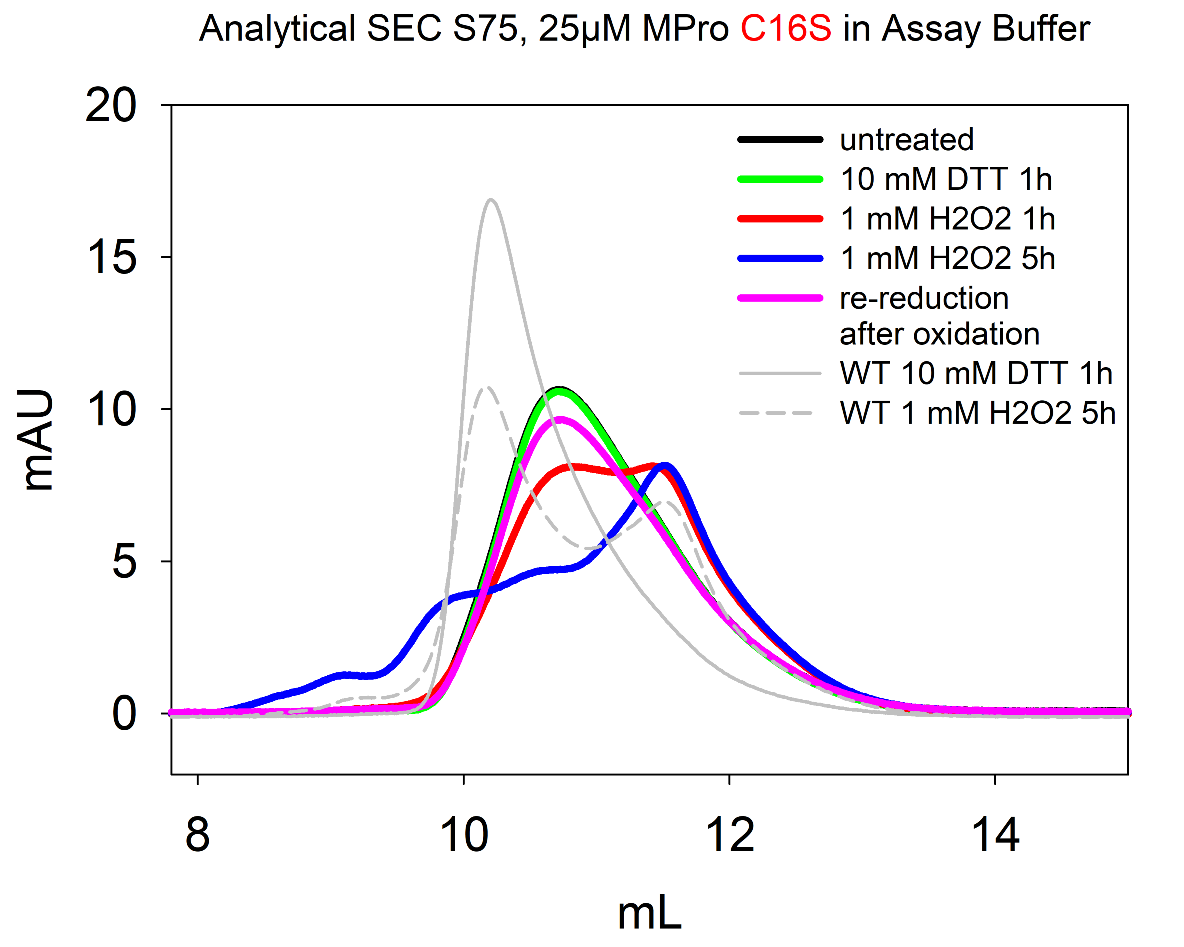

Analytical SEC S75, 25µM MPro C16S in Assay Buffer
20
15
10
5
0
mAU
8
10
12
14
mL
untreated
10 mM DTT 1h
1 mM H2O2 1h
1 mM H2O2 5h
re-reduction
after oxidation
WT 10 mM DTT 1h
WT 1 mM H2O2 5h

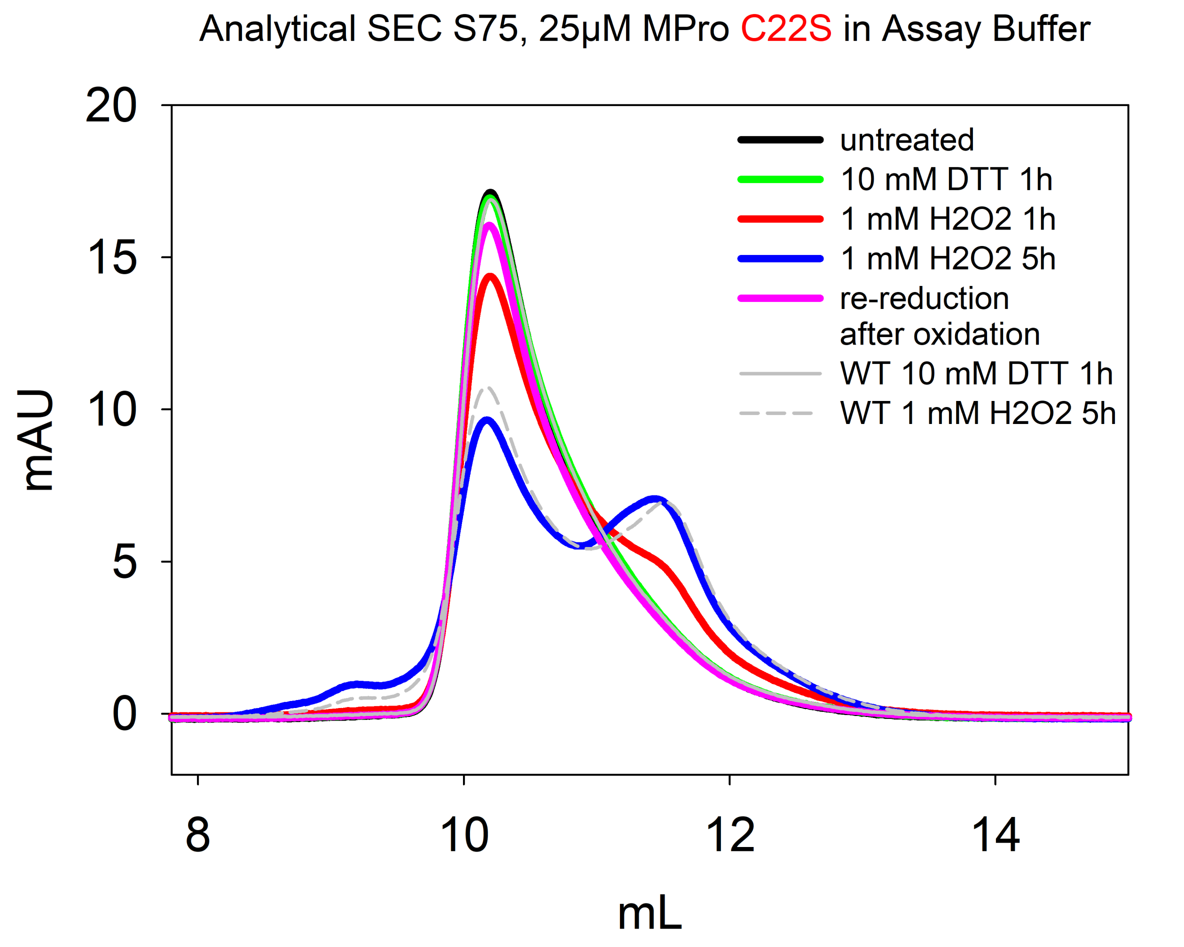

Analytical SEC S75, 25µM MPro C22S in Assay Buffer
20
15
10
5
0
mAU
8
10
12
14
mL
untreated
10 mM DTT 1h
1 mM H2O2 1h
1 mM H2O2 5h
re-reduction
after oxidation
WT 10 mM DTT 1h
WT 1 mM H2O2 5h

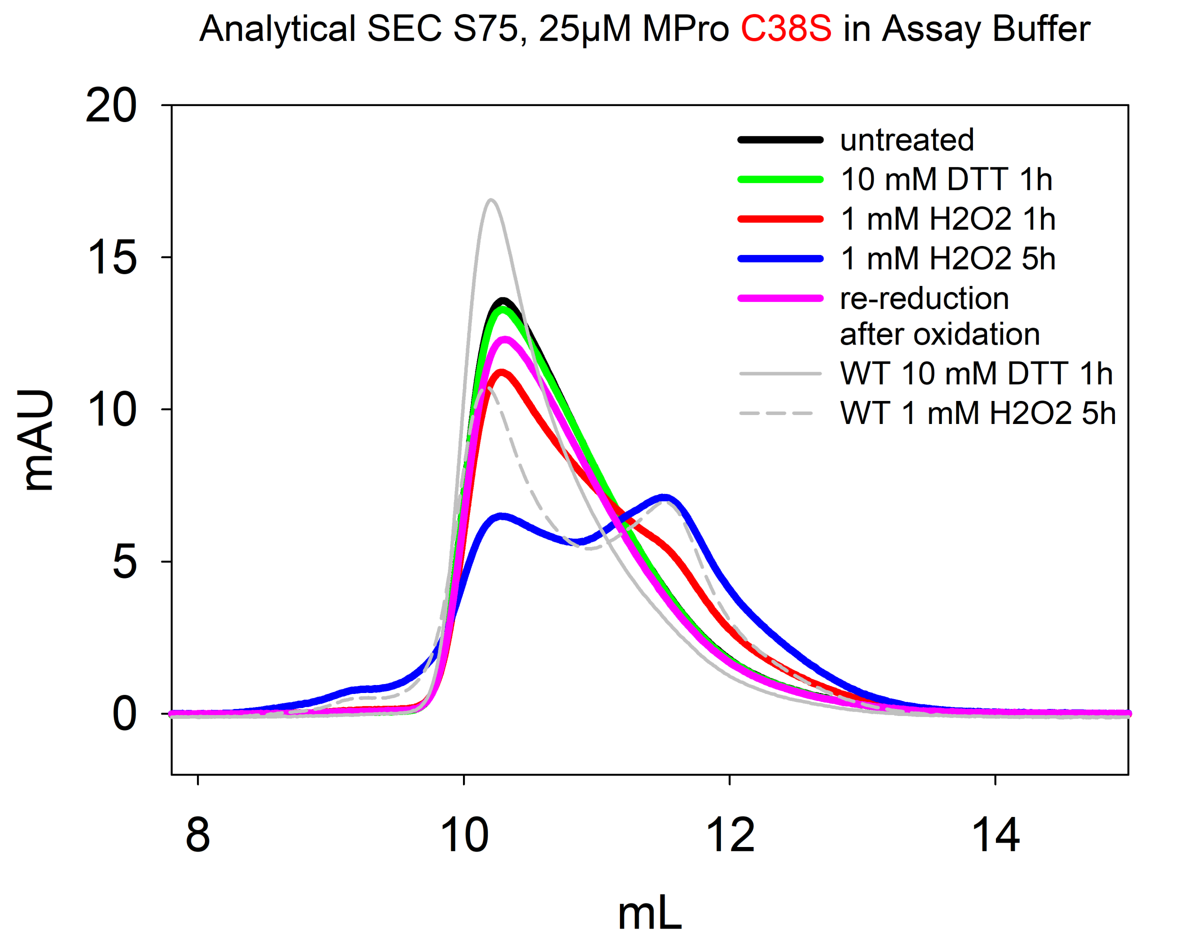

Analytical SEC S75, 25µM MPro C38S in Assay Buffer
20
15
10
5
0
mAU
8
10
12
14
mL
untreated
10 mM DTT 1h
1 mM H2O2 1h
1 mM H2O2 5h
re-reduction after oxidation
WT 10 mM DTT 1h
WT 1 mM H2O2 5h

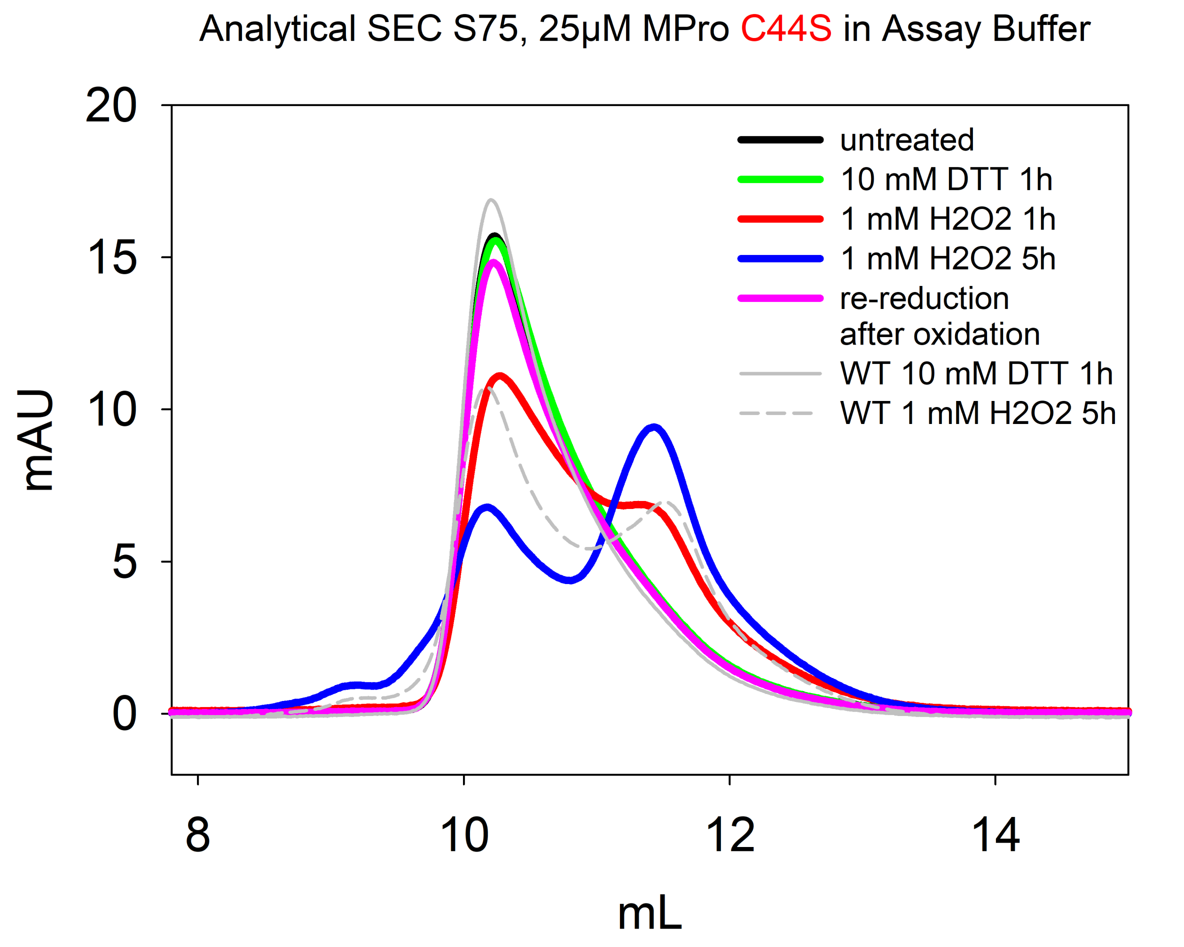

Analytical SEC S75, 25µM MPro C44S in Assay Buffer
20
15
10
5
0
mAU
8
10
12
14
mL
untreated
10 mM DTT 1h
1 mM H2O2 1h
1 mM H2O2 5h
re-reduction
after oxidation
WT 10 mM DTT 1h
WT 1 mM H2O2 5h

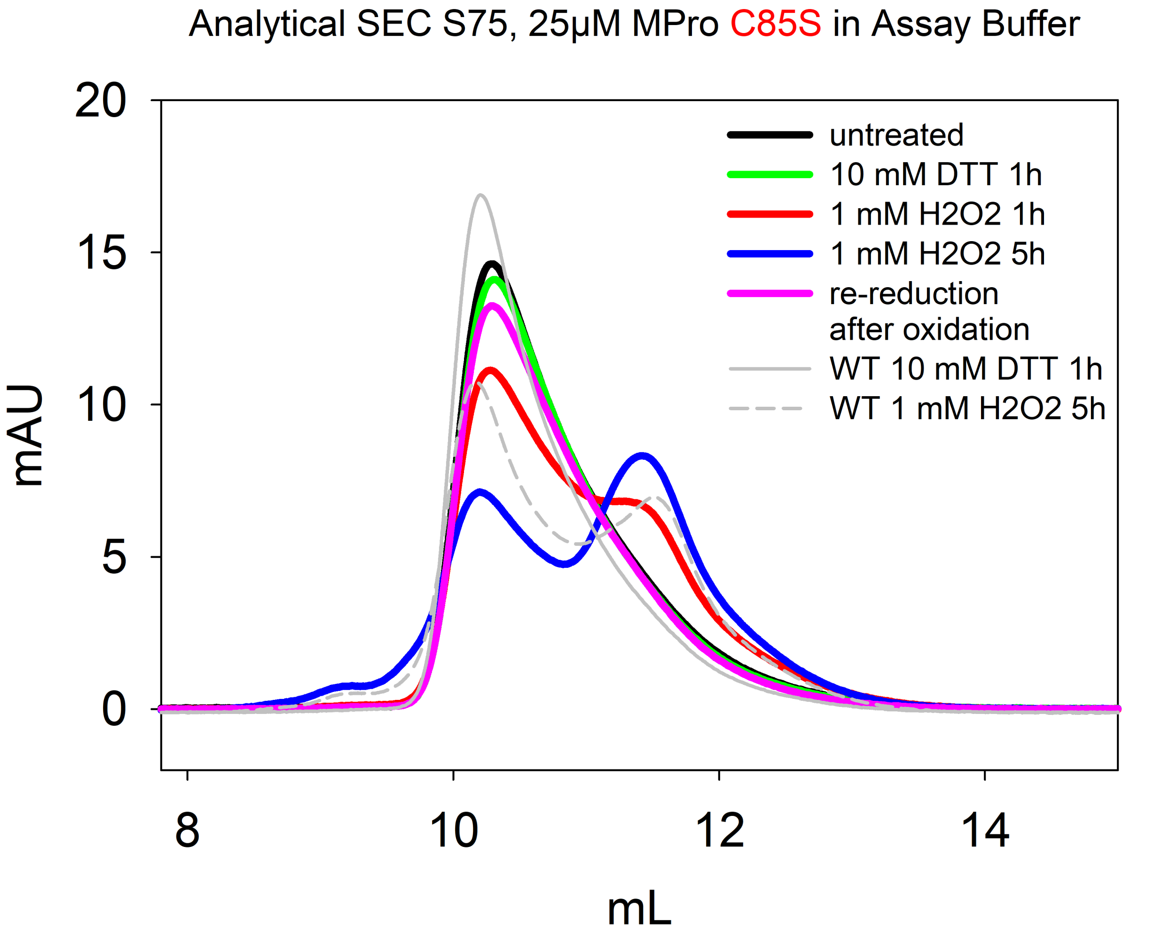
Analytical SEC S75, 25µM MPro C85S in Assay Buffer
20
15
10
5
0
mAU
8
10
12
14
mL
untreated
10 mM DTT 1h
1 mM H2O2 1h
1 mM H2O2 5h
re-reduction
after oxidation
WT 10 mM DTT 1h
WT 1 mM H2O2 5h

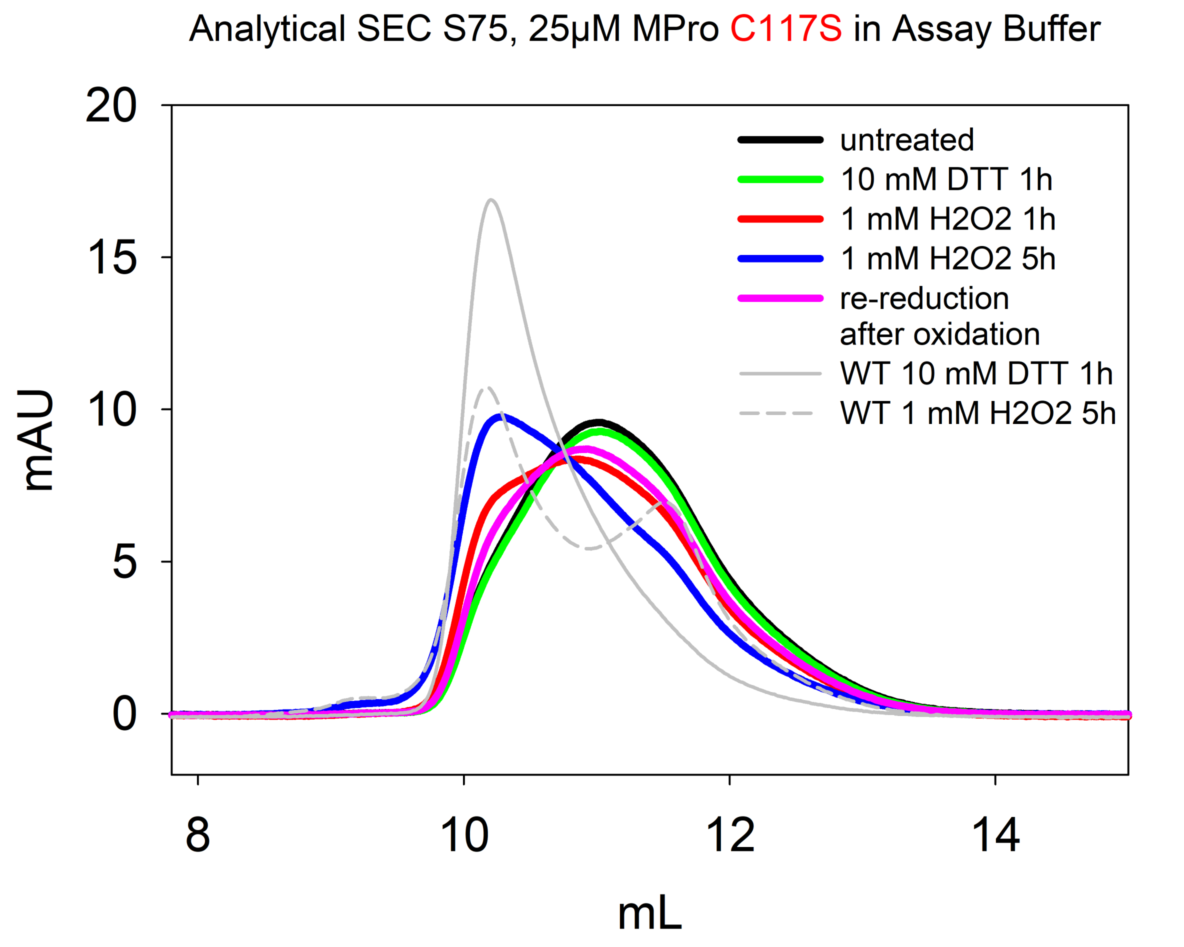
Analytical SEC S75, 25µM MPro C117S in Assay Buffer
untreated
10 mM DTT 1h
1 mM H2O2 1h
1 mM H2O2 5h
re-reduction after oxidation
WT 10 mM DTT 1h
WT 1 mM H2O2 5h
mAU
mL
0
5
10
15
20
8
10
12
14

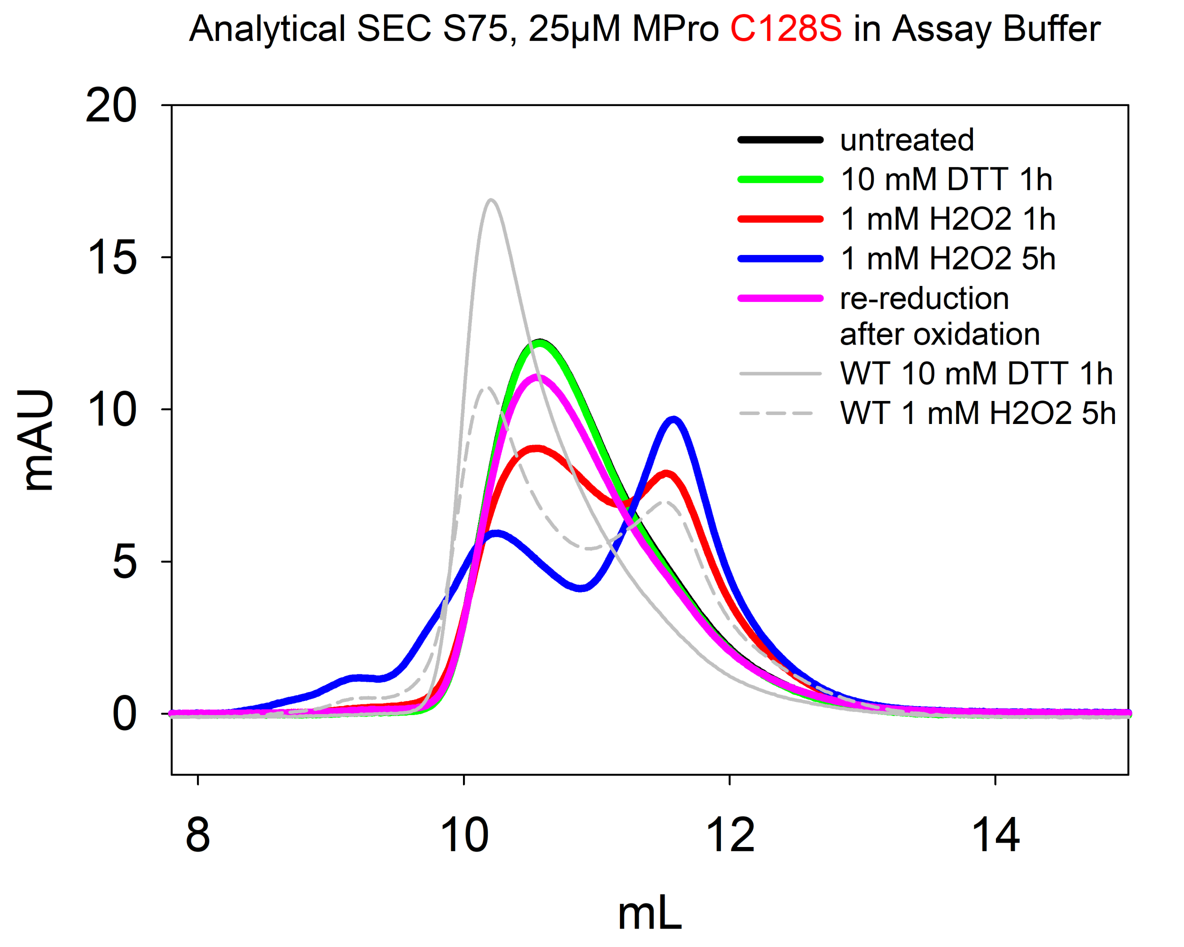
Analytical SEC S75, 25µM MPro C128S in Assay Buffer
20
15
10
5
0
mAU
8
10
12
14
mL
untreated
10 mM DTT 1h
1 mM H2O2 1h
1 mM H2O2 5h
re-reduction after oxidation
WT 10 mM DTT 1h
WT 1 mM H2O2 5h

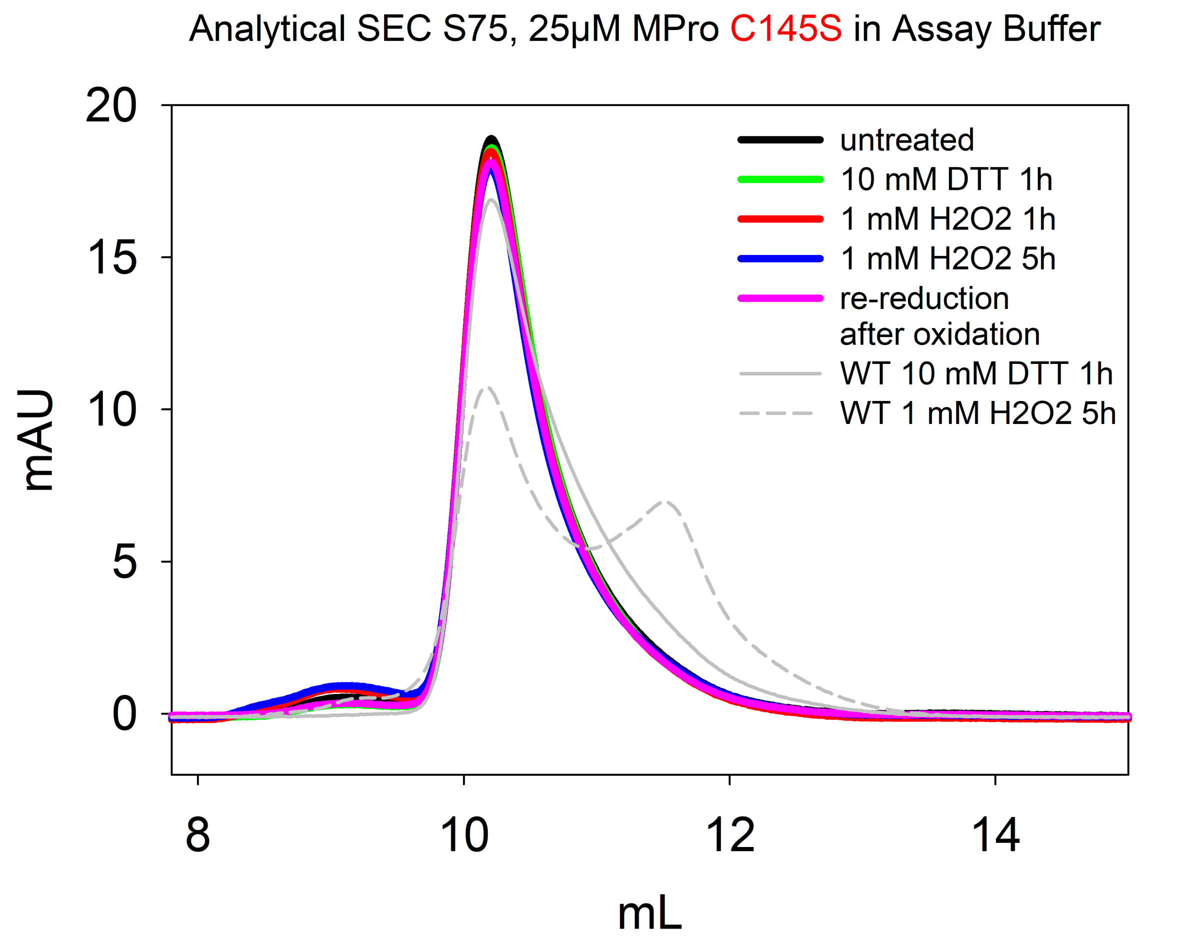

Analytical SEC S75, 25µM MPro C145S in Assay Buffer
20
15
10
5
0
mAU
8
10
12
14
mL
untreated
10 mM DTT 1h
1 mM H2O2 1h
1 mM H2O2 5h
re-reduction after oxidation
WT 10 mM DTT 1h
WT 1 mM H2O2 5h

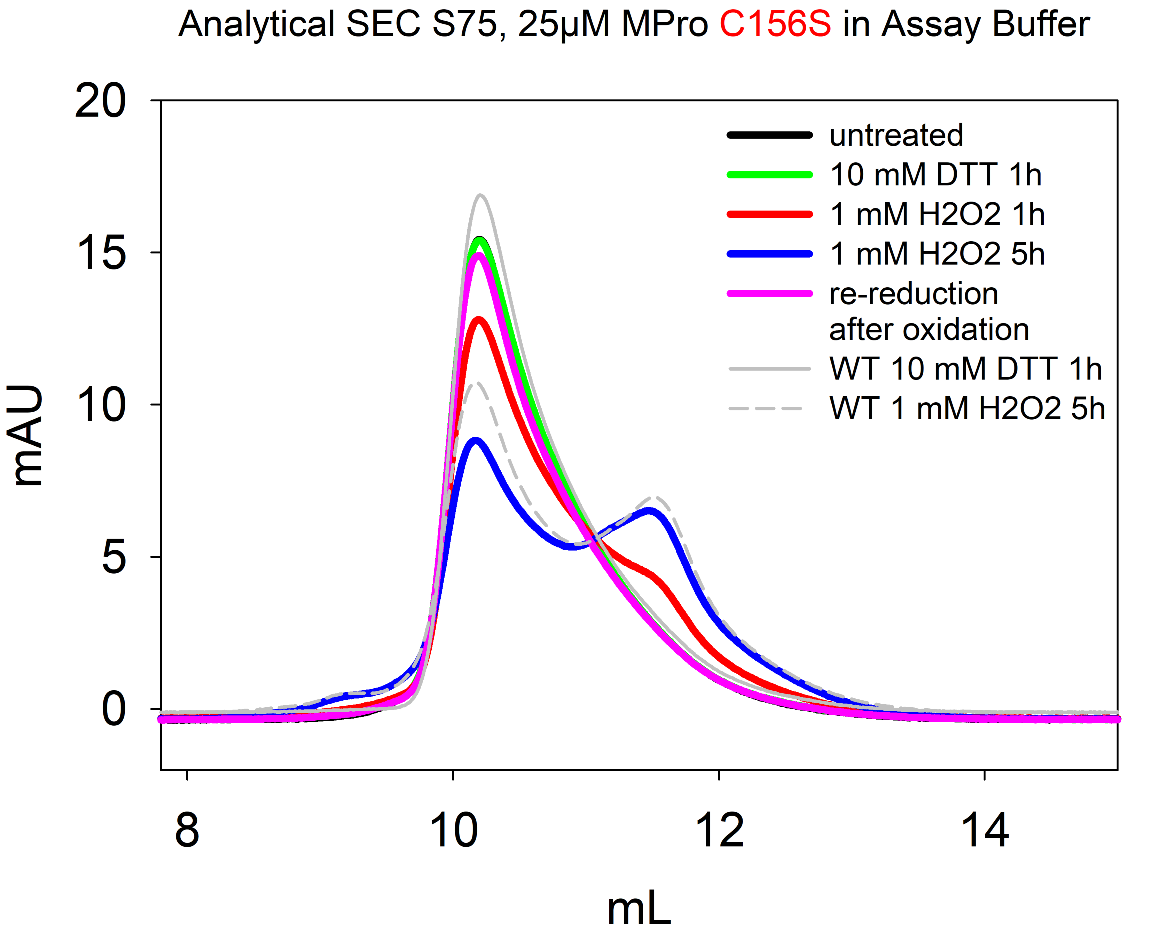
Analytical SEC S75, 25µM MPro C156S in Assay Buffer
20
15
10
5
0
mAU
8
10
12
14
mL
untreated
10 mM DTT 1h
1 mM H2O2 1h
1 mM H2O2 5h
re-reduction
after oxidation
WT 10 mM DTT 1h
WT 1 mM H2O2 5h

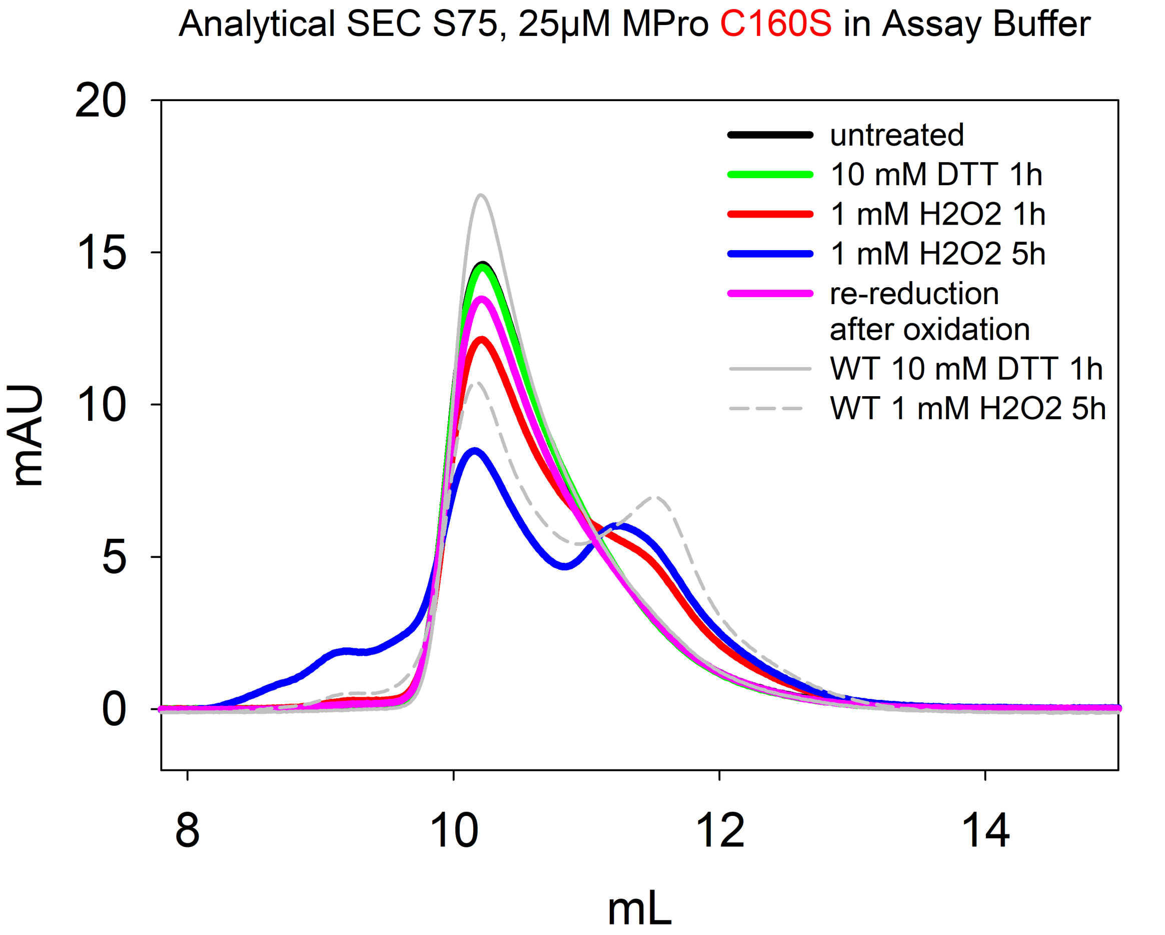

Analytical SEC S75, 25µM MPro C160S in Assay Buffer
untreated
10 mM DTT 1h
1 mM H2O2 1h
1 mM H2O2 5h
re-reduction after oxidation
WT 10 mM DTT 1h
WT 1 mM H2O2 5h
mAU
mL
0
5
10
15
20
8
10
12
14

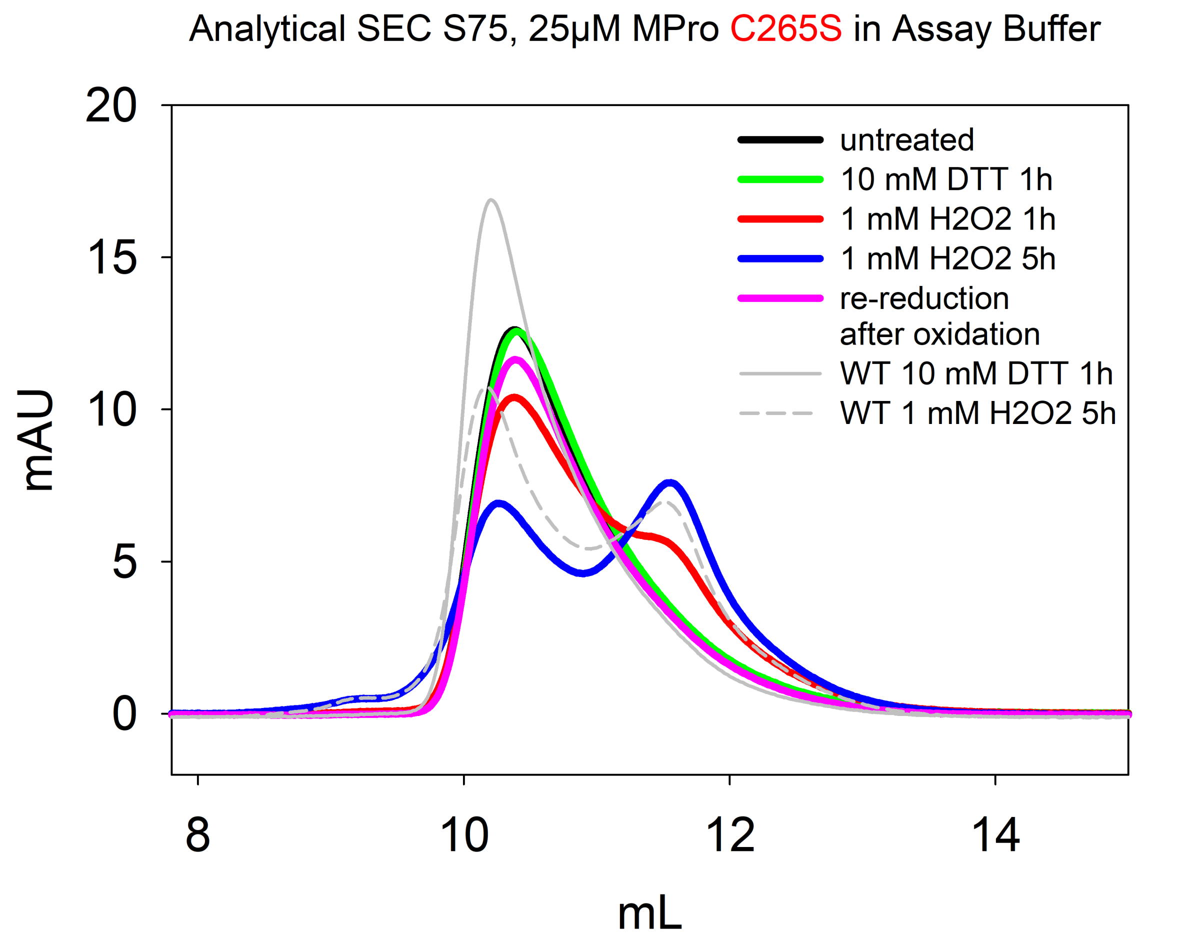
Analytical SEC S75, 25µM MPro C265S in Assay Buffer
20
15
10
5
0
mAU
8
10
12
14
mL
untreated
10 mM DTT 1h
1 mM H2O2 1h
1 mM H2O2 5h
re-reduction after oxidation
WT 10 mM DTT 1h
WT 1 mM H2O2 5h

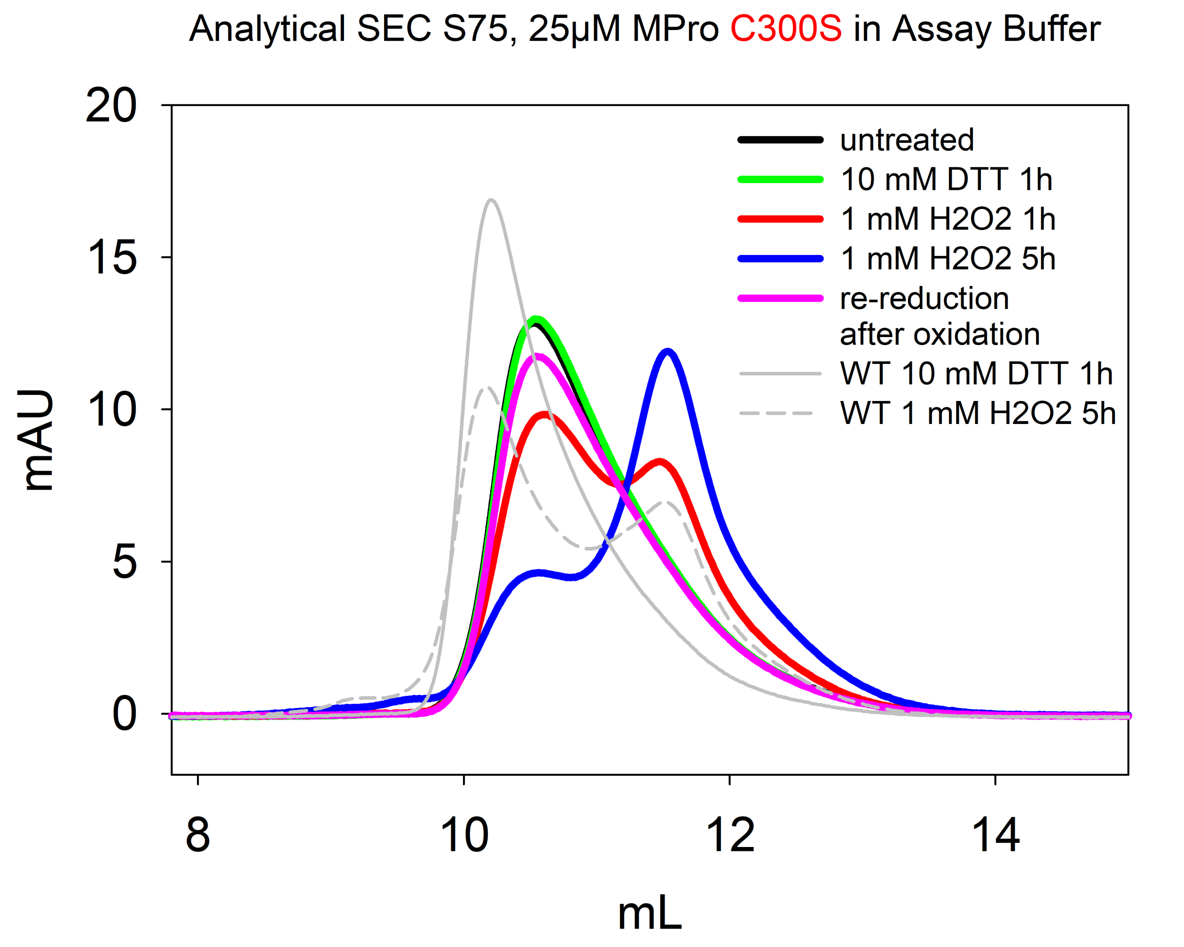

Analytical SEC S75, 25µM MPro C300S in Assay Buffer
20
15
10
5
0
mAU
8
10
12
14
mL
untreated
10 mM DTT 1h
1 mM H2O2 1h
1 mM H2O2 5h
re-reduction
after oxidation
WT 10 mM DTT 1h
WT 1 mM H2O2 5h

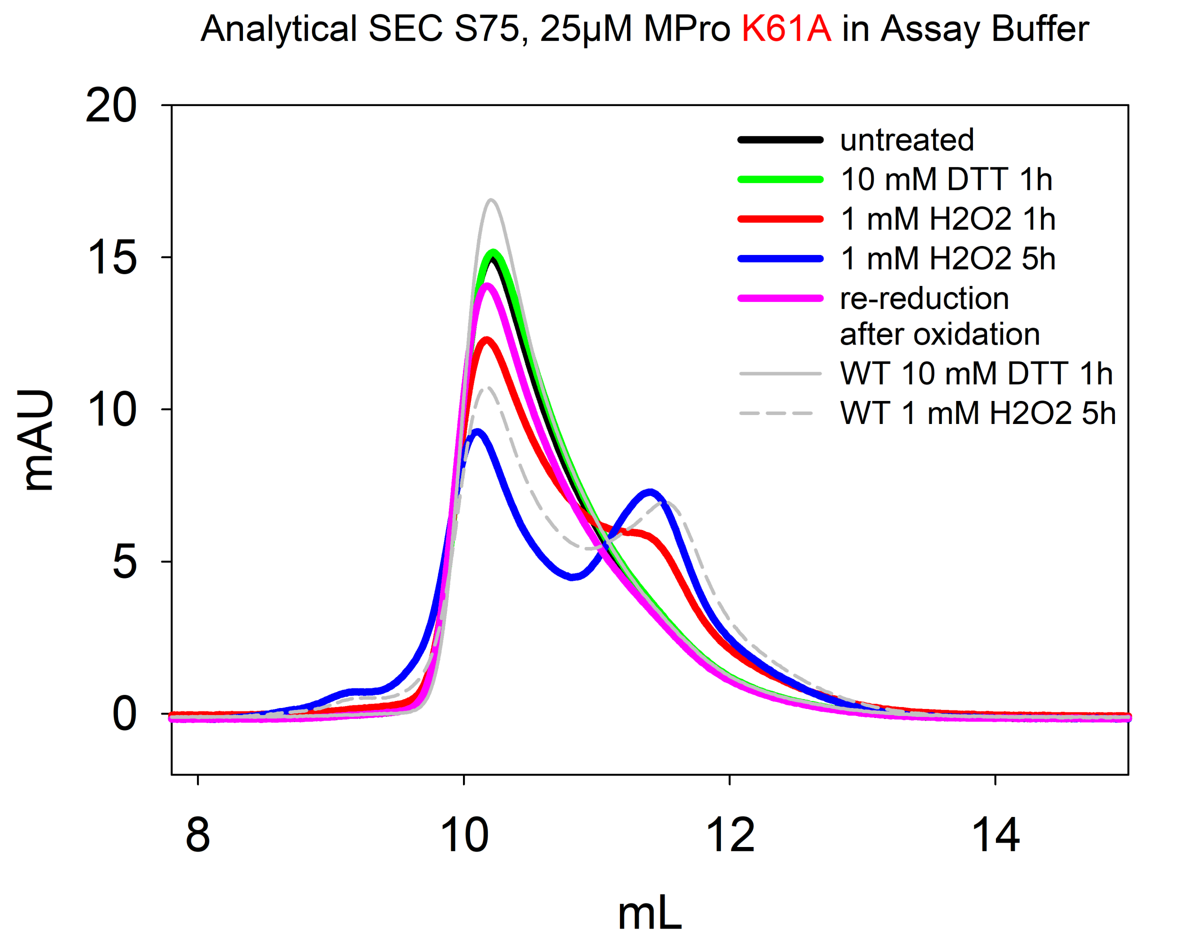
Analytical SEC S75, 25µM MPro K61A in Assay Buffer
20
15
10
5
0
mAU
8
10
12
14
mL
untreated
10 mM DTT 1h
1 mM H2O2 1h
1 mM H2O2 5h
re-reduction
after oxidation
WT 10 mM DTT 1h
WT 1 mM H2O2 5h

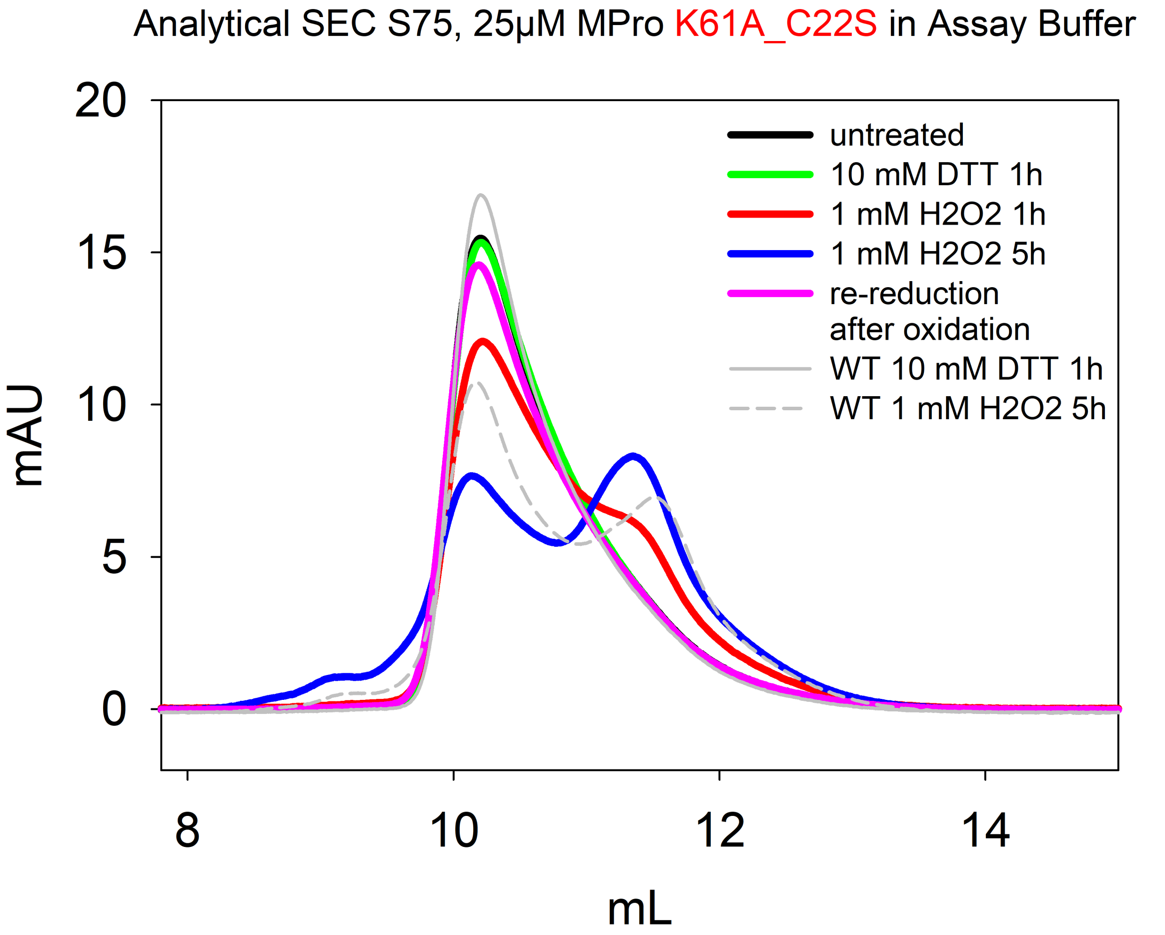
Analytical SEC S75, 25µM MPro K61A_C22S in Assay Buffer
mAU
mL
20
15
10
5
0
8
10
12
14
untreated
10 mM DTT 1h
1 mM H2O2 1h
1 mM H2O2 5h
re-reduction
after oxidation
WT 10 mM DTT 1h
WT 1 mM H2O2 5h

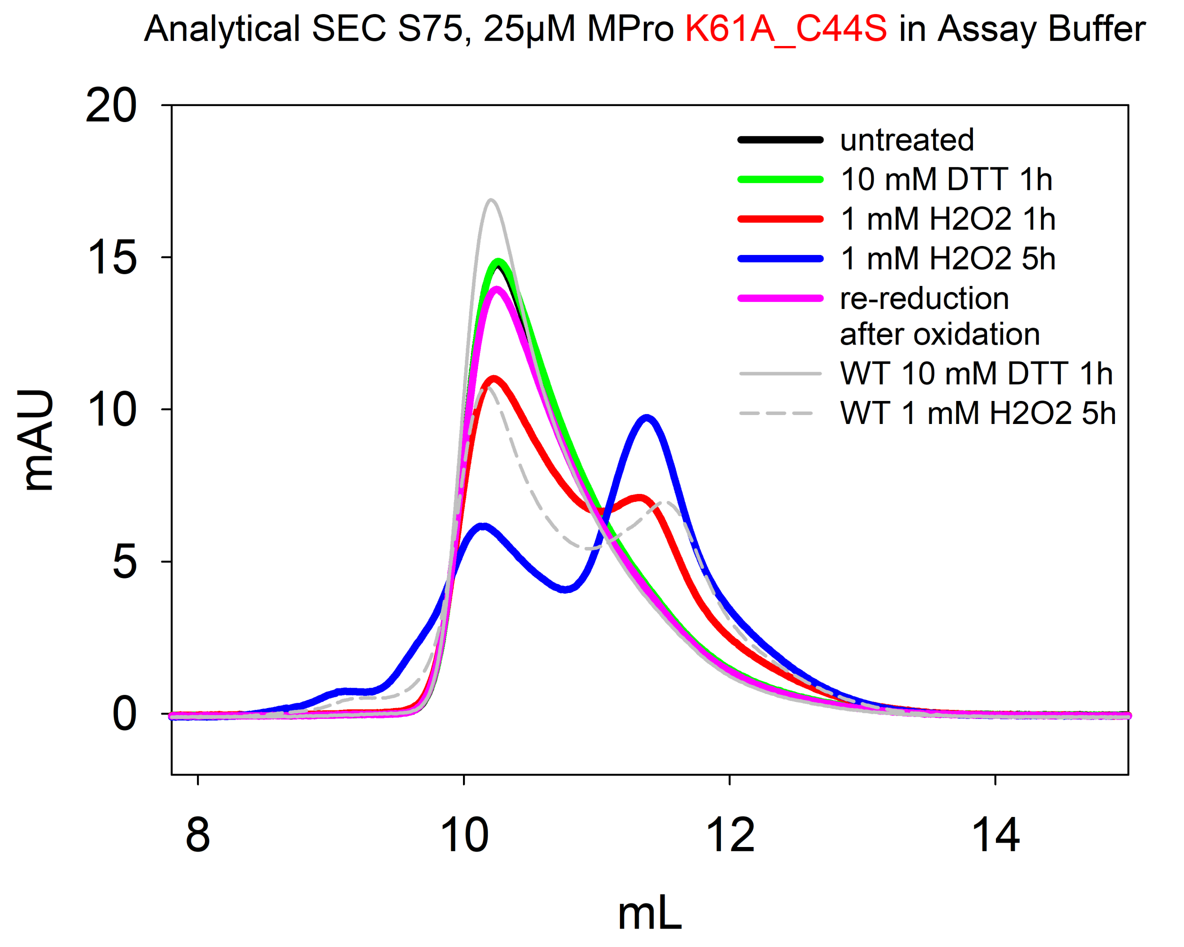
Analytical SEC S75, 25µM MPro K61A_C44S in Assay Buffer
untreated
10 mM DTT 1h
1 mM H2O2 1h
1 mM H2O2 5h
re-reduction after oxidation
WT 10 mM DTT 1h
WT 1 mM H2O2 5h
mAU
mL
20
15
10
5
0
8
10
12
14

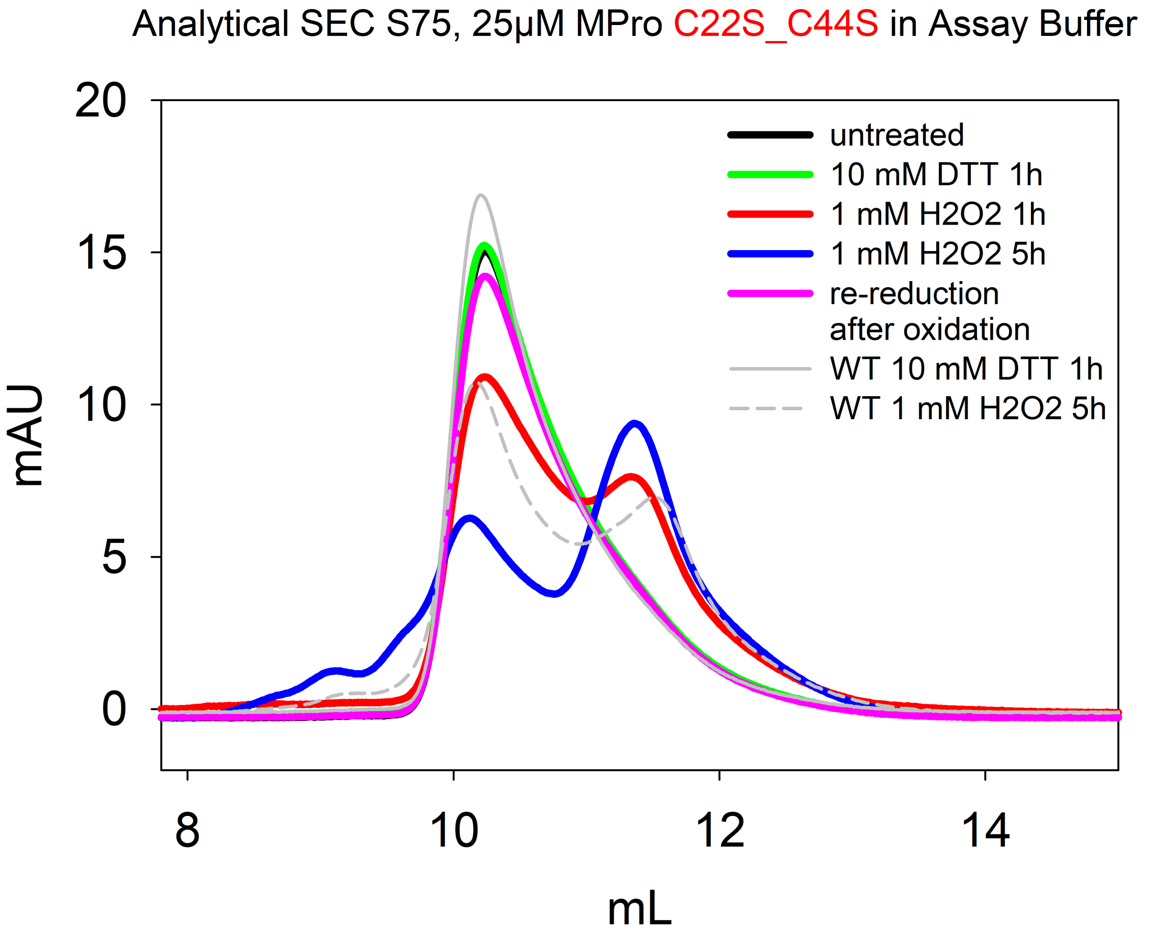

Analytical SEC S75, 25µM MPro C22S_C44S in Assay Buffer
mAU
20
15
10
5
0
8
10
12
14
mL
untreated
10 mM DTT 1h
1 mM H2O2 1h
1 mM H2O2 5h
re-reduction
after oxidation
WT 10 mM DTT 1h
WT 1 mM H2O2 5h

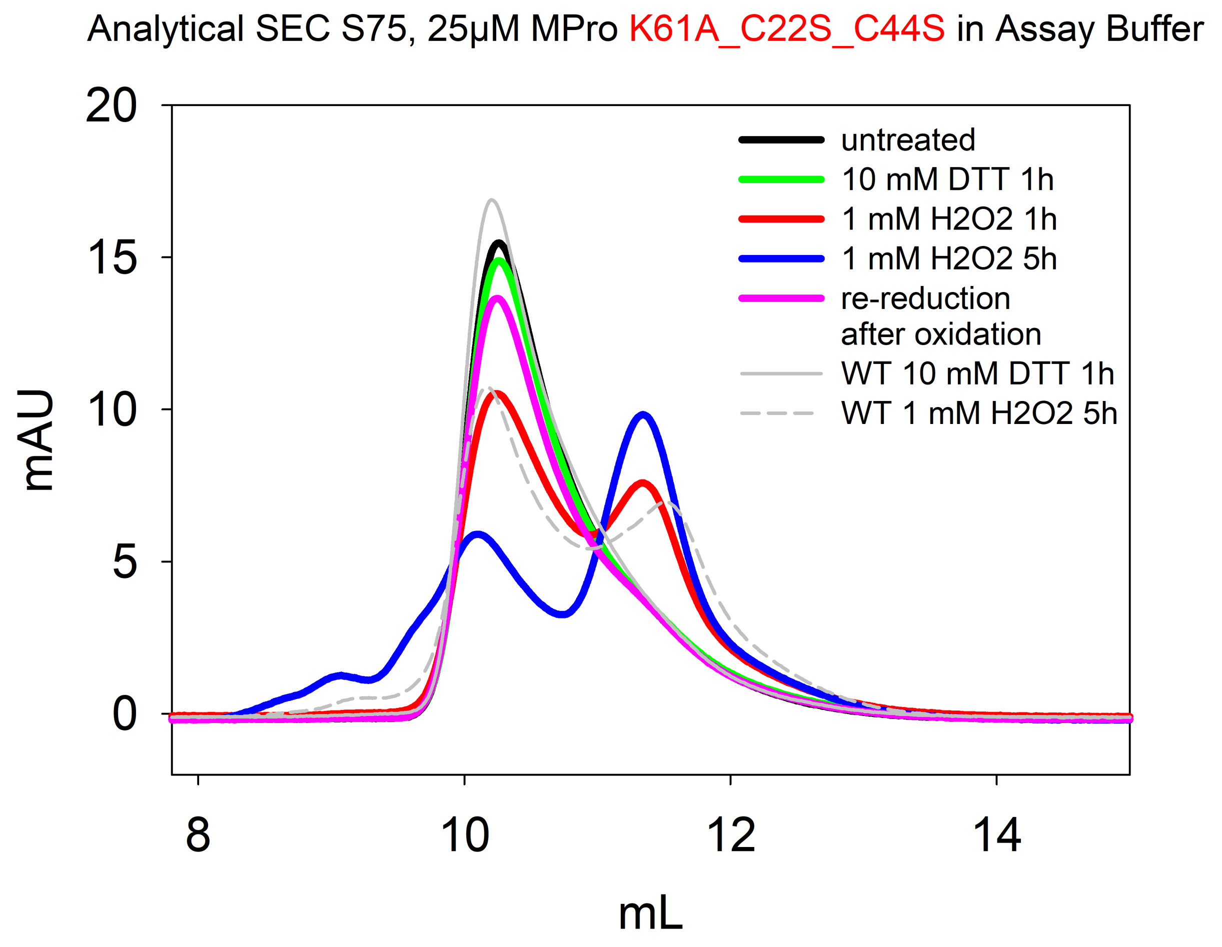

Analytical SEC S75, 25µM MPro K61A_C22S_C44S in Assay Buffer
20
15
10
5
0
mAU
8
10
12
14
mL
untreated
10 mM DTT 1h
1 mM H2O2 1h
1 mM H2O2 5h
re-reduction after oxidation
WT 10 mM DTT 1h
WT 1 mM H2O2 5h

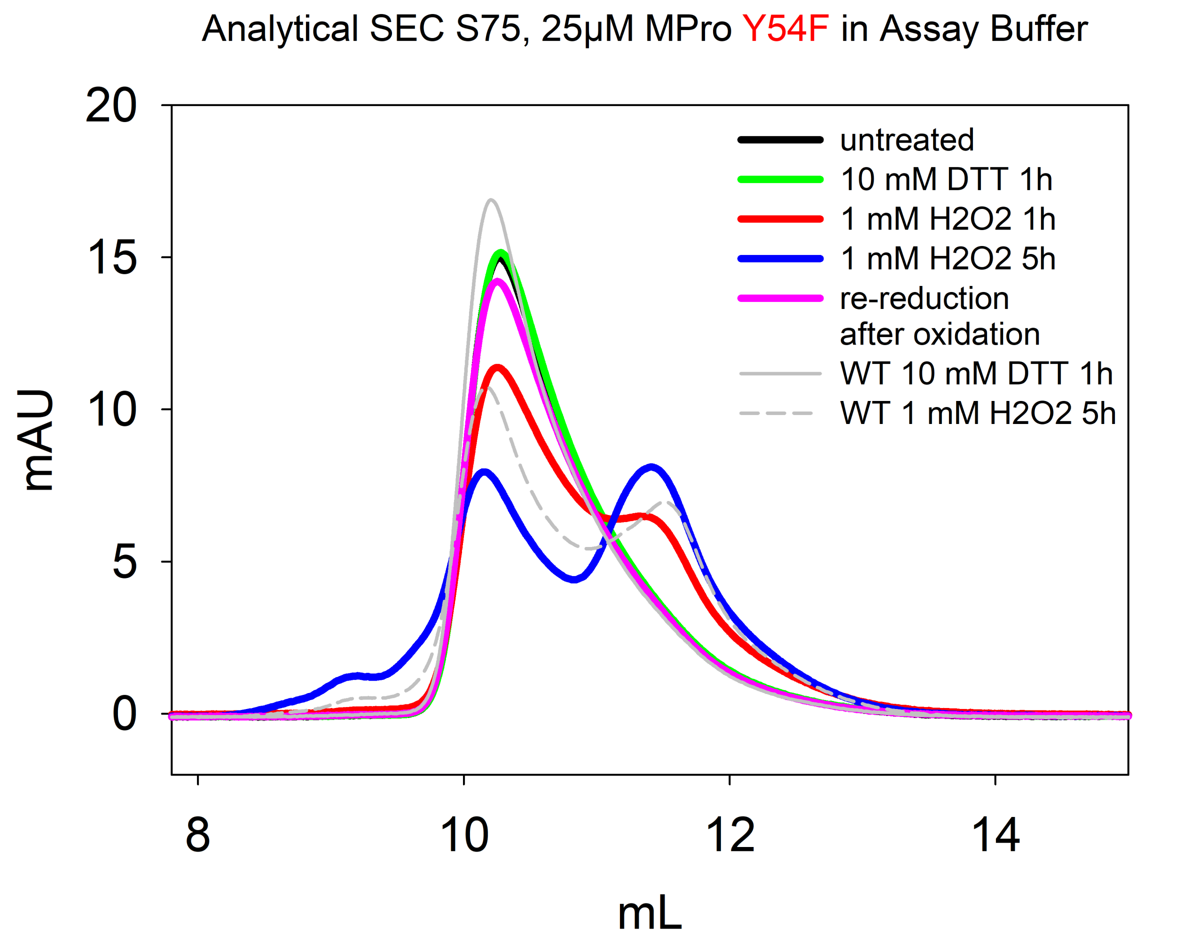
Analytical SEC S75, 25µM MPro Y54F in Assay Buffer
mAU
mL
20
15
10
5
0
8
10
12
14
untreated
10 mM DTT 1h
1 mM H2O2 1h
1 mM H2O2 5h
re-reduction
after oxidation
WT 10 mM DTT 1h
WT 1 mM H2O2 5h
