## Appendix 2 - CD spectra for "Redox regulation of the SARS-CoV-2 main protease provides new opportunities for drug design"

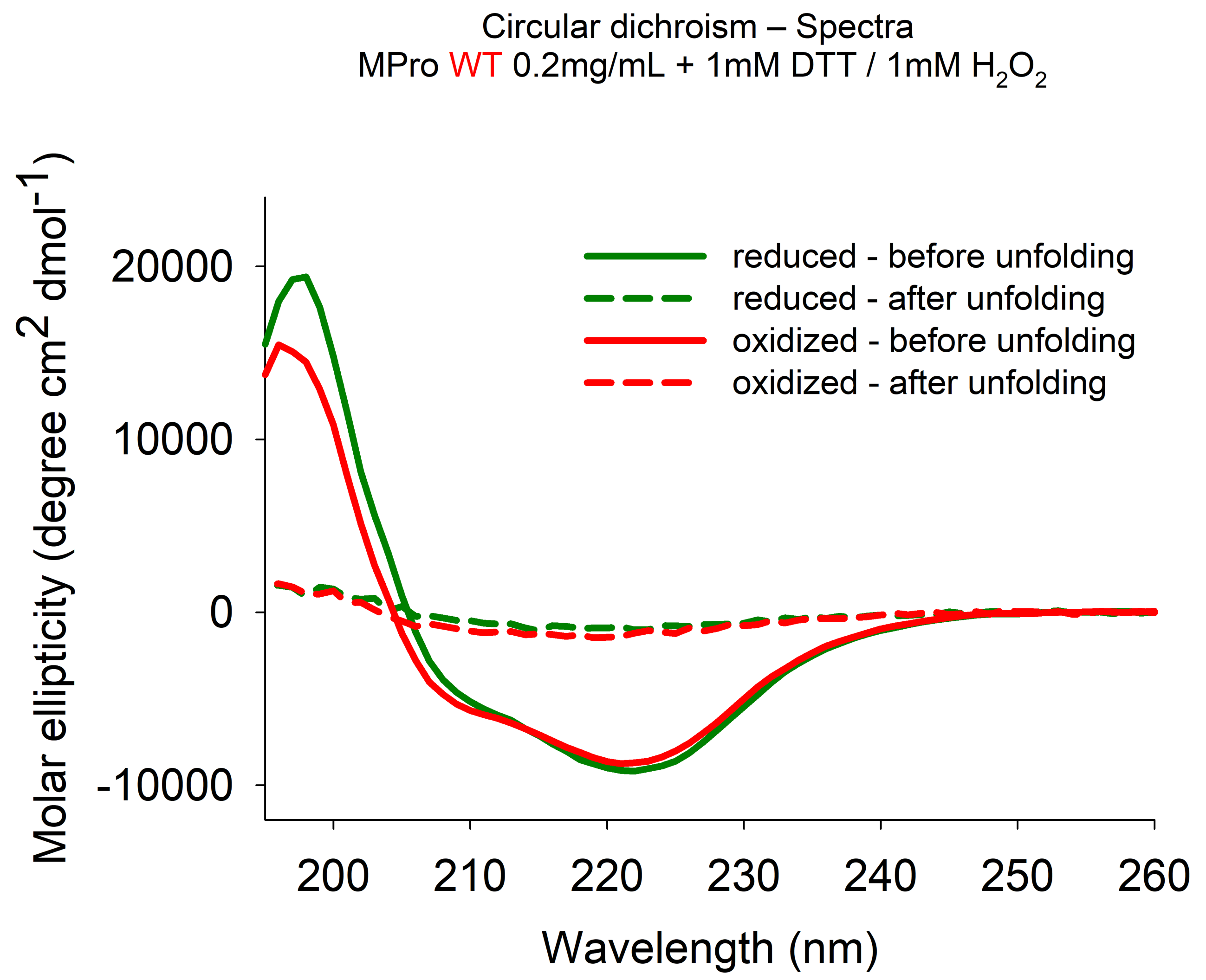
Circular dichroism – Spectra
MPro WT 0.2mg/mL + 1mM DTT / 1mM H2O2
Molar ellipticity (degree cm2 dmol-1)
20000
10000
0
-10000
200
210
220
230
240
250
260
Wavelength (nm)
reduced - before unfolding
reduced - after unfolding
oxidized - before unfolding
oxidized - after unfolding

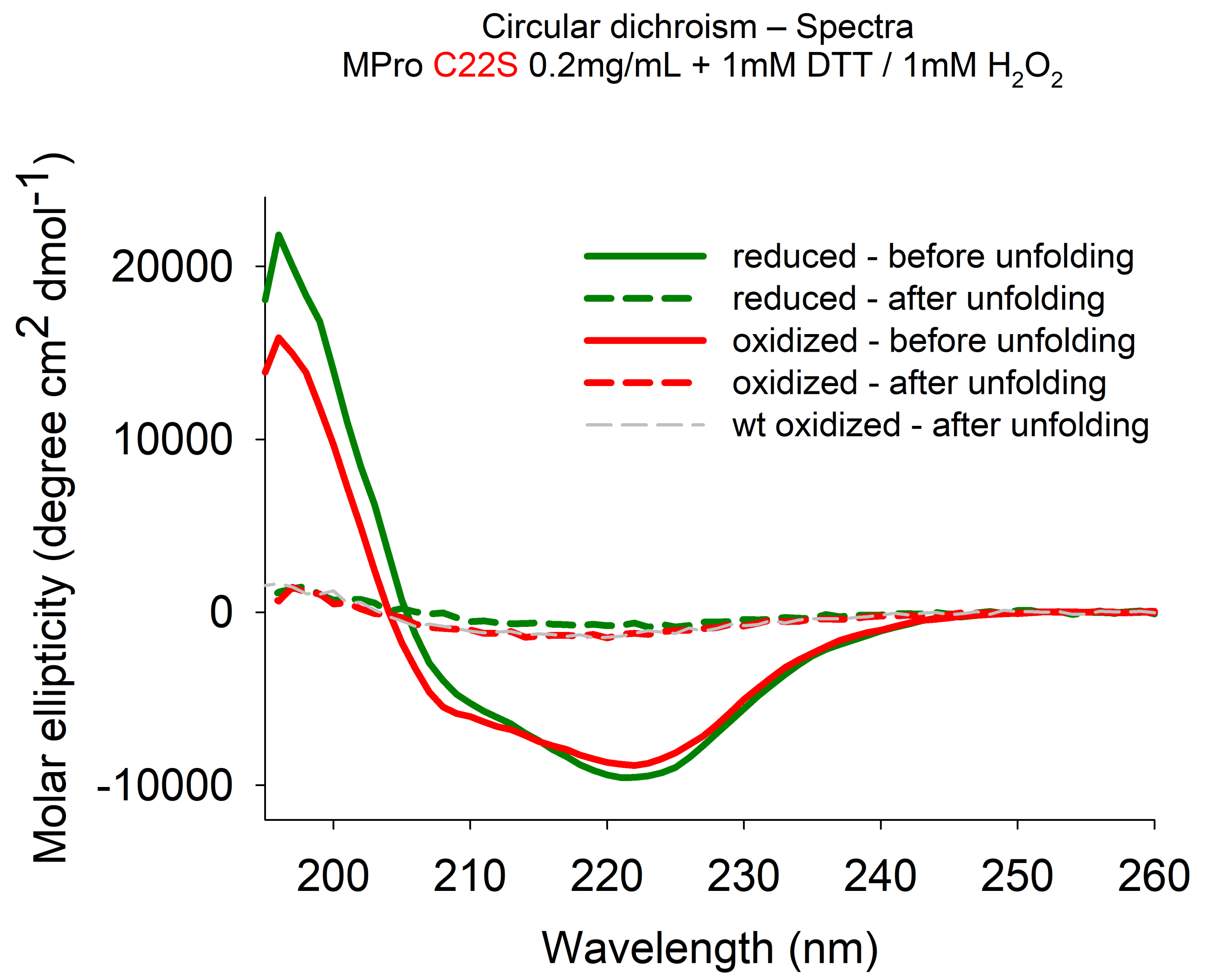
Circular dichroism – Spectra
MPro C22S 0.2mg/mL + 1mM DTT / 1mM $H_2O_2$
Molar ellipticity (degree cm$^2$ dmol$^{-1}$)
20000
10000
0
-10000
200
210
220
230
240
250
260
Wavelength (nm)
reduced - before unfolding
reduced - after unfolding
oxidized - before unfolding
oxidized - after unfolding
wt oxidized - after unfolding

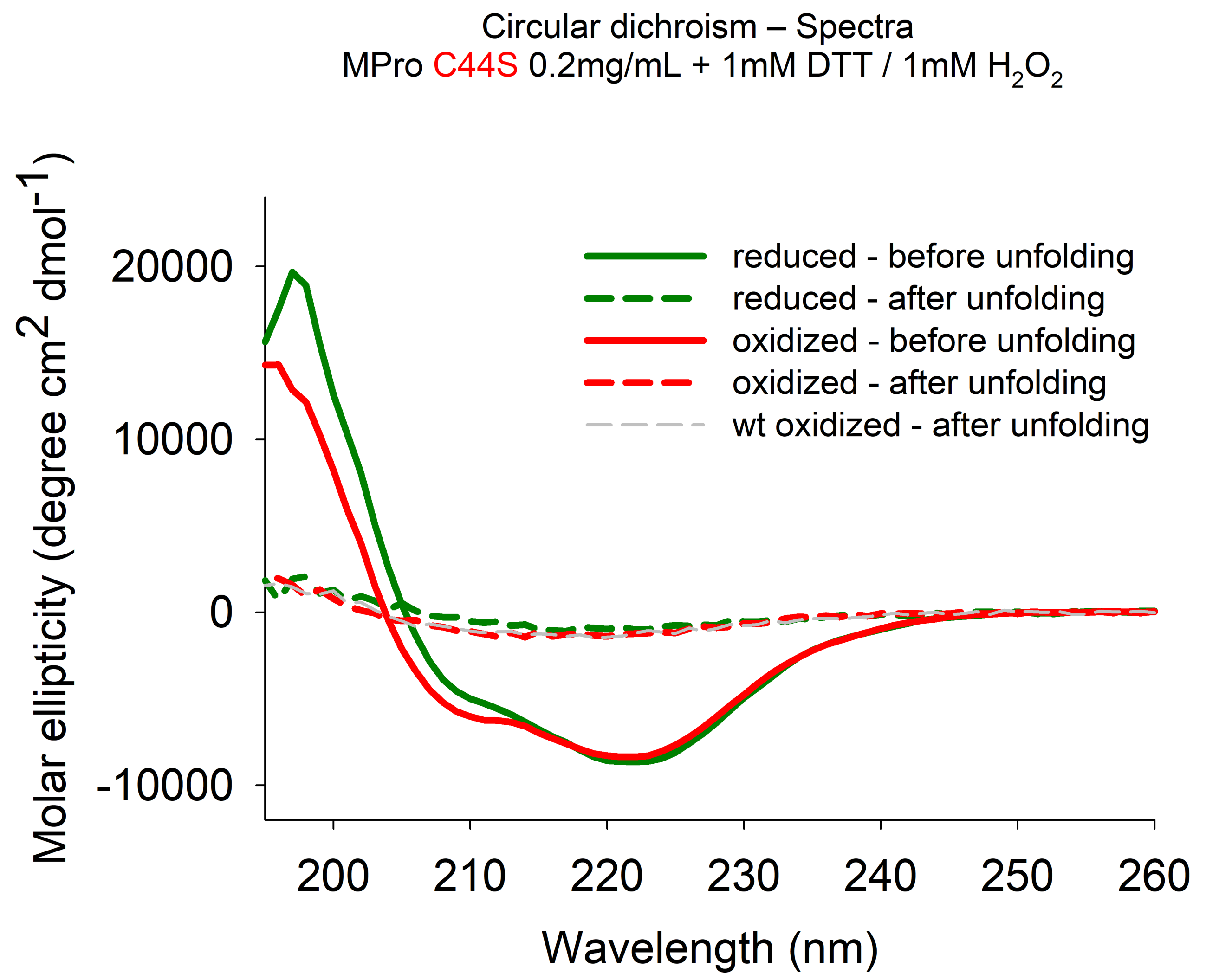
Circular dichroism – Spectra
MPro C44S 0.2mg/mL + 1mM DTT / 1mM $H_2O_2$
reduced - before unfolding
reduced - after unfolding
oxidized - before unfolding
oxidized - after unfolding
wt oxidized - after unfolding
Molar ellipticity (degree cm$^2$ dmol$^{-1}$)
20000
10000
0
-10000
200
210
220
230
240
250
260
Wavelength (nm)

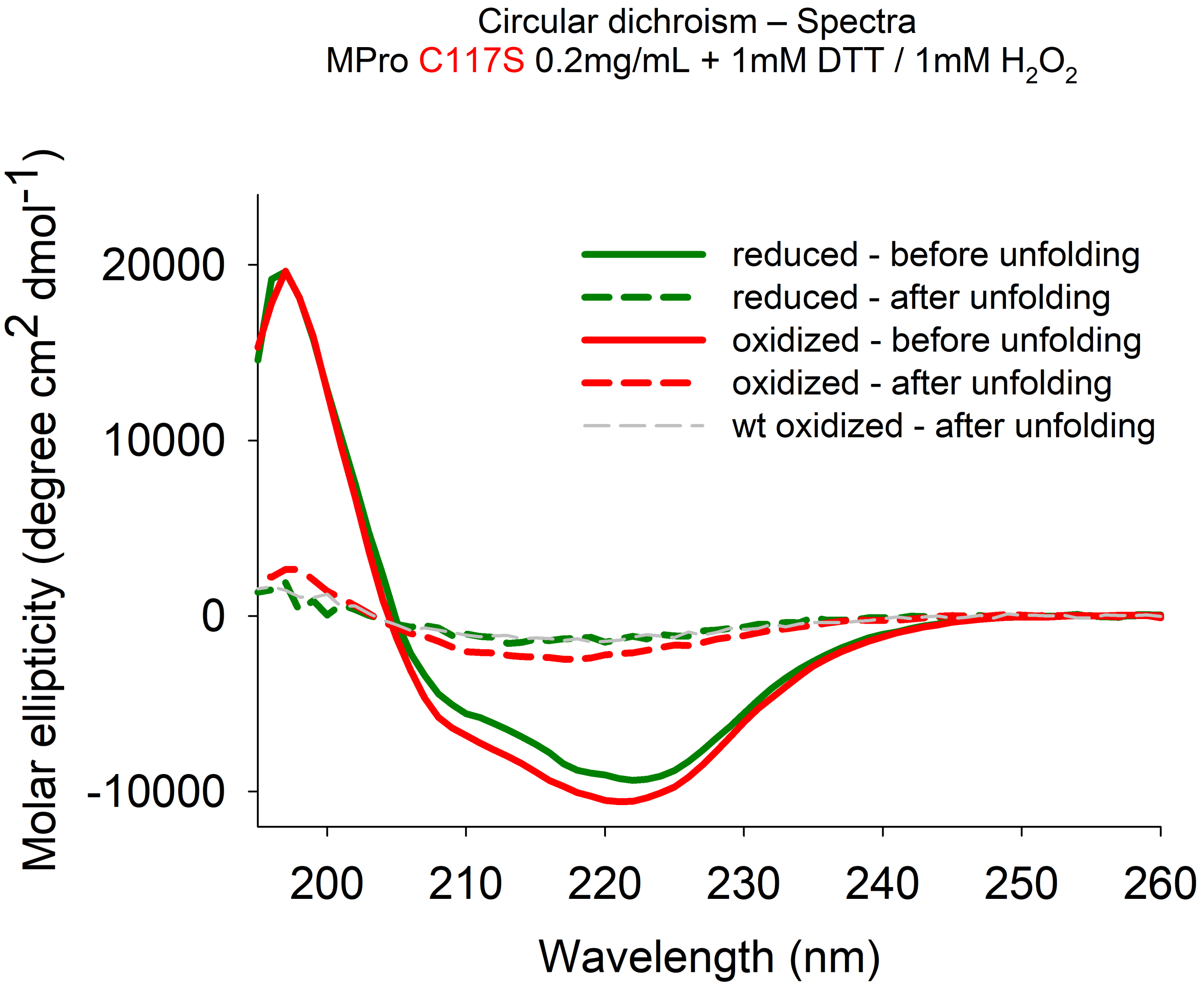

Circular dichroism – Spectra
MPro C117S 0.2mg/mL + 1mM DTT / 1mM H2O2
Molar ellipticity (degree cm2 dmol-1)
20000
10000
0
-10000
200
210
220
230
240
250
260
Wavelength (nm)
reduced - before unfolding
reduced - after unfolding
oxidized - before unfolding
oxidized - after unfolding
wt oxidized - after unfolding

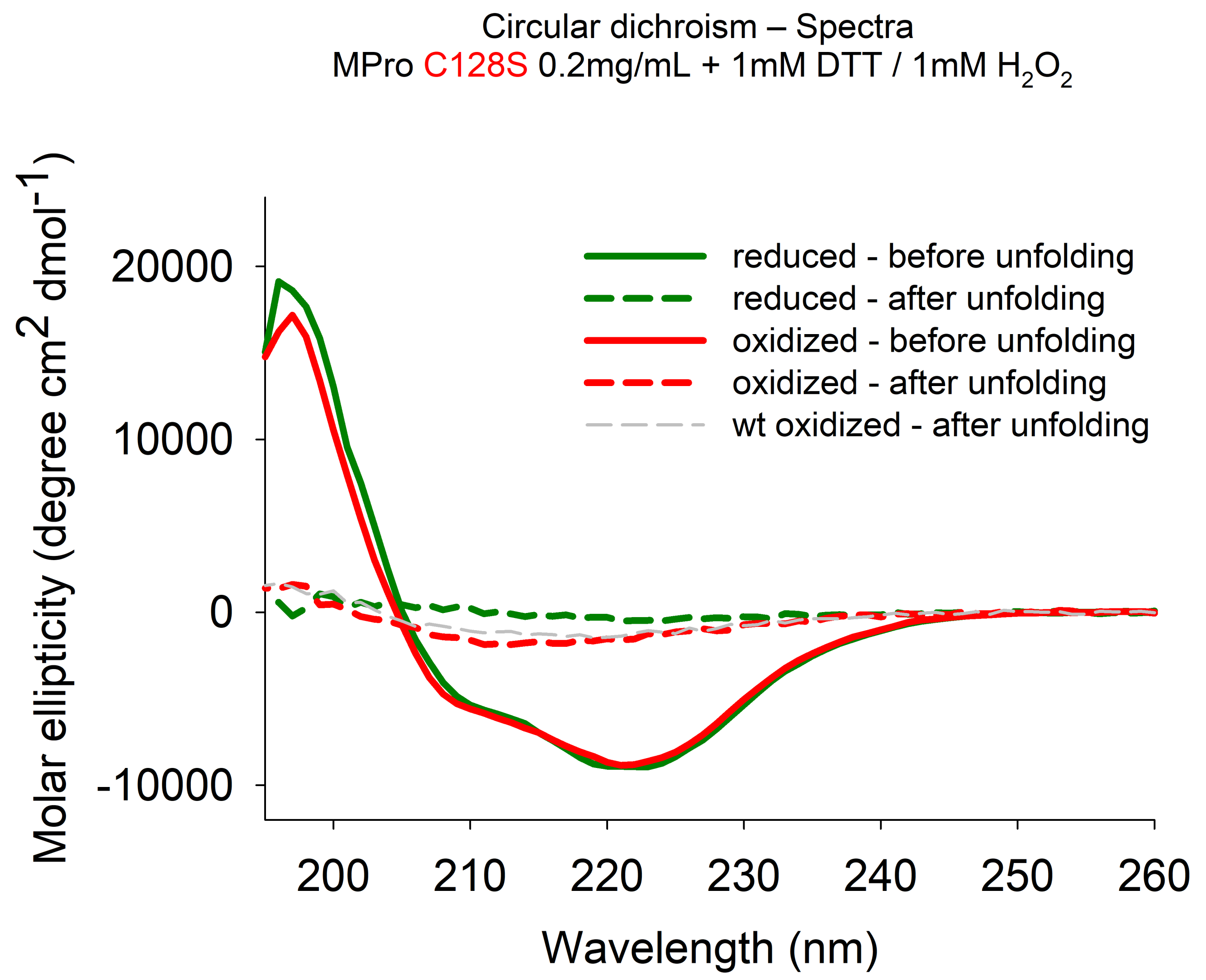

Circular dichroism – Spectra
MPro C128S 0.2mg/mL + 1mM DTT / 1mM $H_2O_2$
Molar ellipticity (degree cm$^2$ dmol$^{-1}$)
20000
10000
0
-10000
200
210
220
230
240
250
260
Wavelength (nm)
reduced - before unfolding
reduced - after unfolding
oxidized - before unfolding
oxidized - after unfolding
wt oxidized - after unfolding

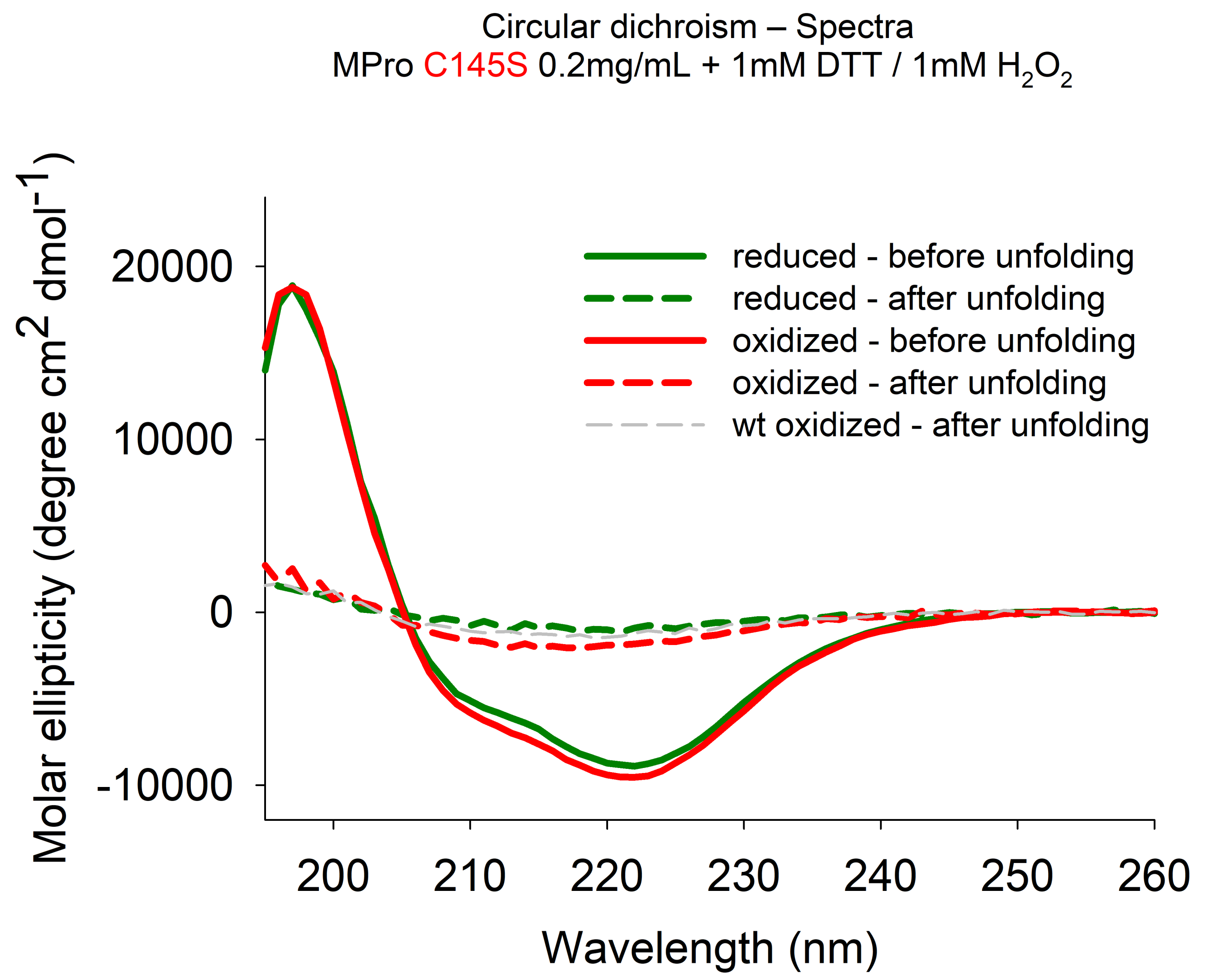

Circular dichroism – Spectra
MPro C145S 0.2mg/mL + 1mM DTT / 1mM $H_2O_2$
Molar ellipticity (degree $cm^2$ $dmol^{-1}$)
20000
10000
0
-10000
200
210
220
230
240
250
260
Wavelength (nm)
reduced - before unfolding
reduced - after unfolding
oxidized - before unfolding
oxidized - after unfolding
wt oxidized - after unfolding

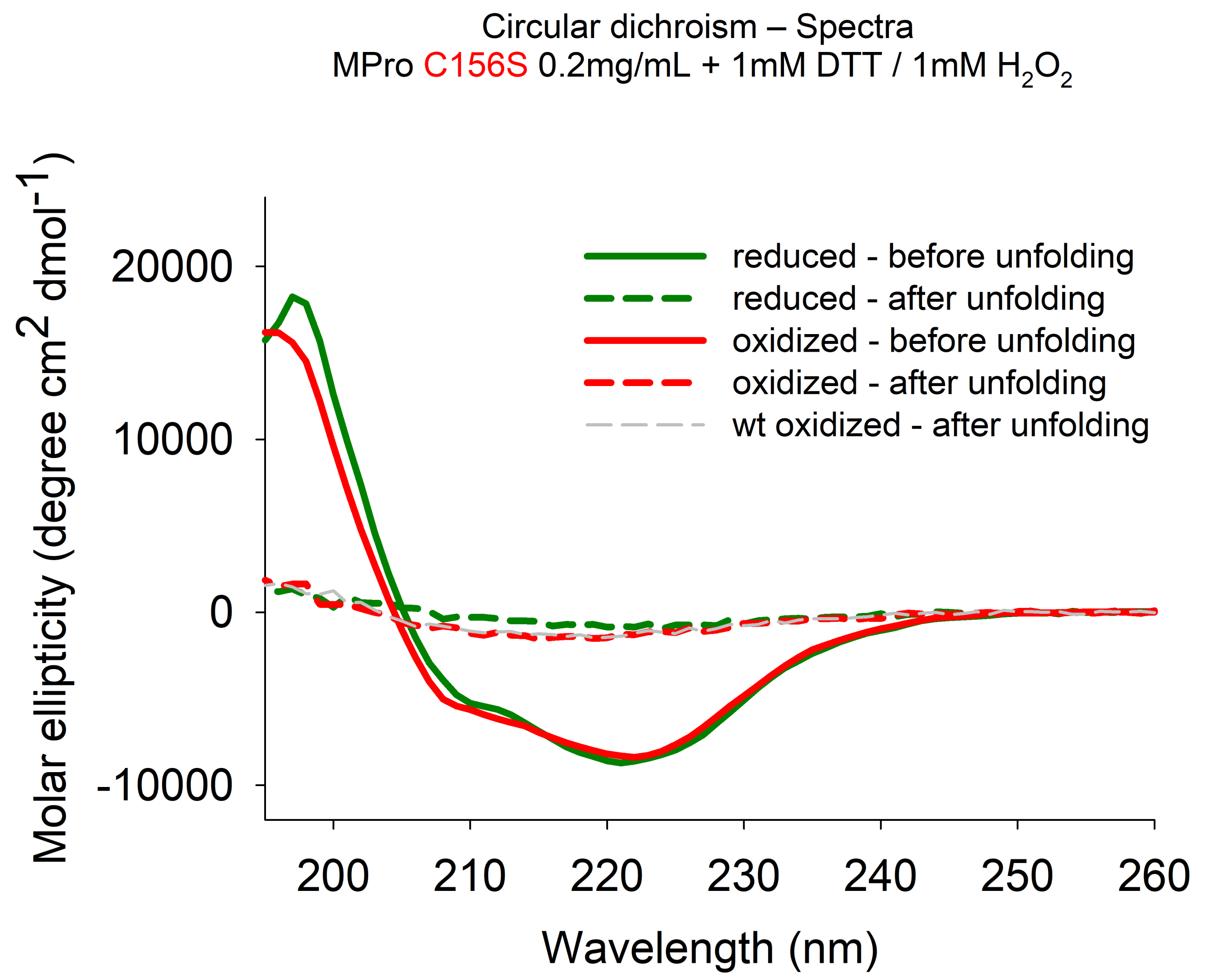
Circular dichroism – Spectra
MPro C156S 0.2mg/mL + 1mM DTT / 1mM $H_2O_2$
Molar ellipticity (degree $cm^2$ $dmol^{-1}$)
20000
10000
0
-10000
200
210
220
230
240
250
260
Wavelength (nm)
reduced - before unfolding
reduced - after unfolding
oxidized - before unfolding
oxidized - after unfolding
wt oxidized - after unfolding

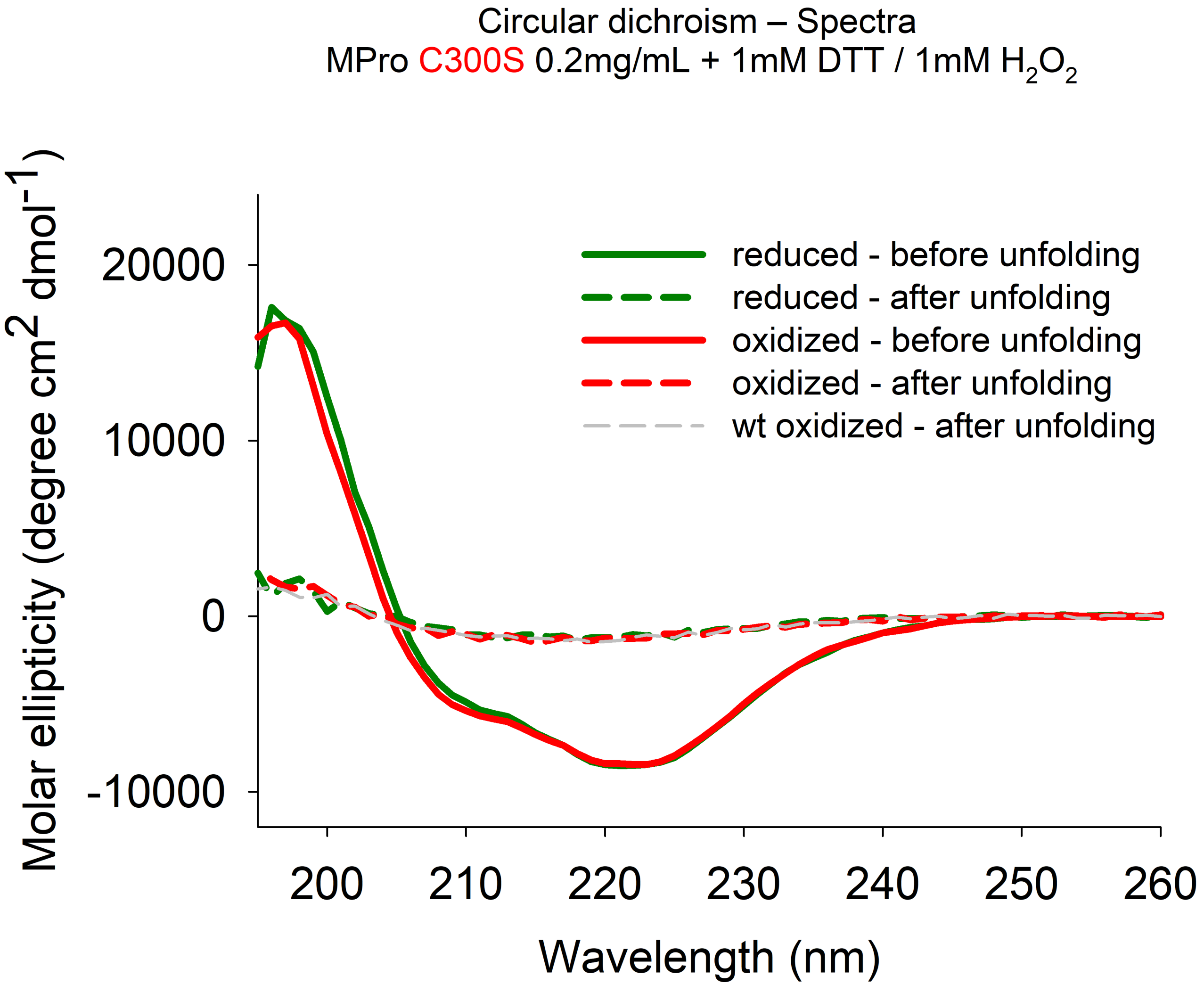

Circular dichroism – Spectra
MPro C300S 0.2mg/mL + 1mM DTT / 1mM $H_2O_2$
Molar ellipticity (degree cm$^2$ dmol$^{-1}$)
20000
10000
0
-10000
200
210
220
230
240
250
260
Wavelength (nm)
reduced - before unfolding
reduced - after unfolding
oxidized - before unfolding
oxidized - after unfolding
wt oxidized - after unfolding

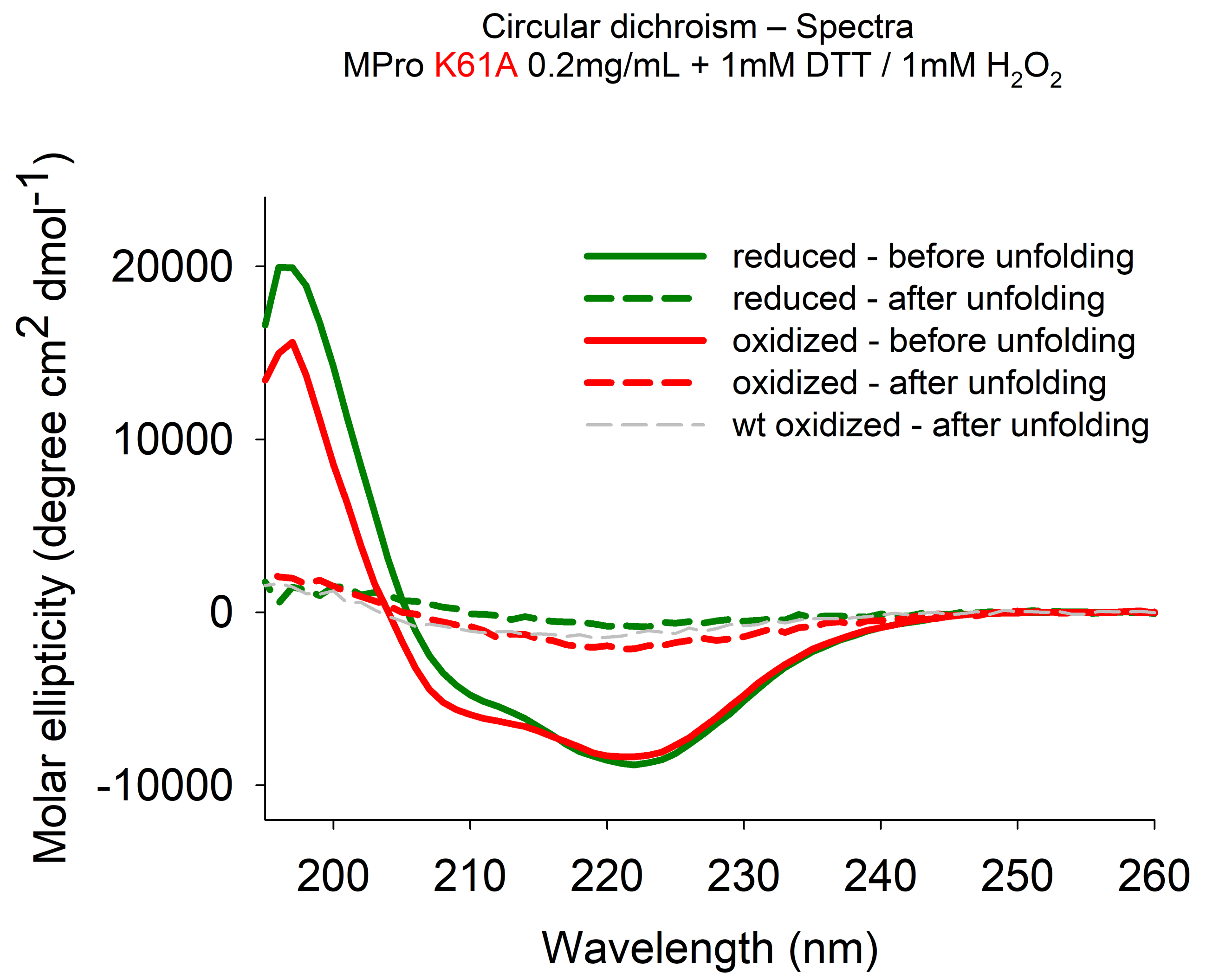
Circular dichroism – Spectra
MPro K61A 0.2mg/mL + 1mM DTT / 1mM $H_2O_2$
Molar ellipticity (degree cm$^2$ dmol$^{-1}$)
20000
10000
0
-10000
200
210
220
230
240
250
260
Wavelength (nm)
reduced - before unfolding
reduced - after unfolding
oxidized - before unfolding
oxidized - after unfolding
wt oxidized - after unfolding

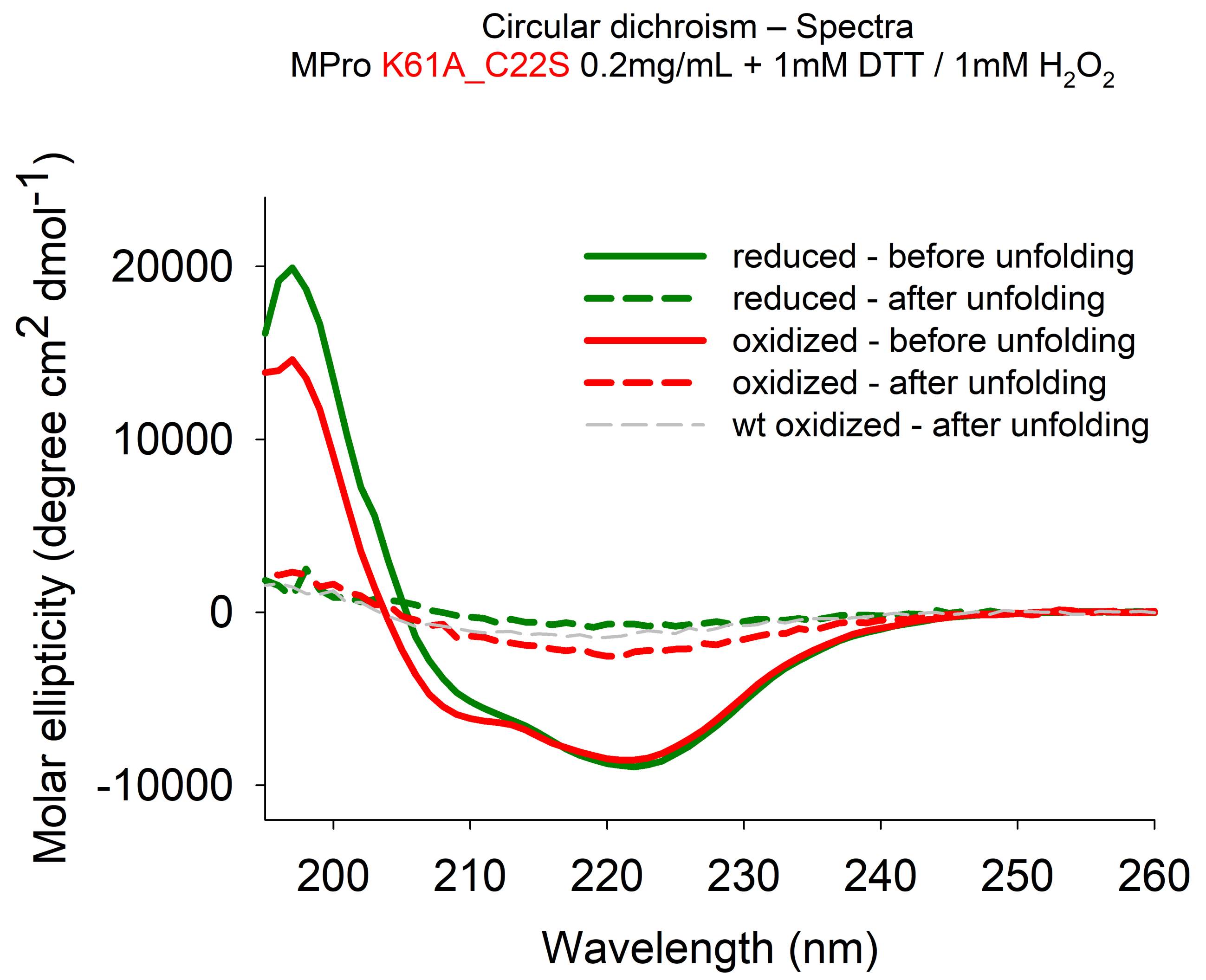
Circular dichroism – Spectra
MPro K61A_C22S 0.2mg/mL + 1mM DTT / 1mM $H_2O_2$
Molar ellipticity (degree $cm^2$ $dmol^{-1}$)
20000
10000
0
-10000
200
210
220
230
240
250
260
Wavelength (nm)
reduced - before unfolding
reduced - after unfolding
oxidized - before unfolding
oxidized - after unfolding
wt oxidized - after unfolding

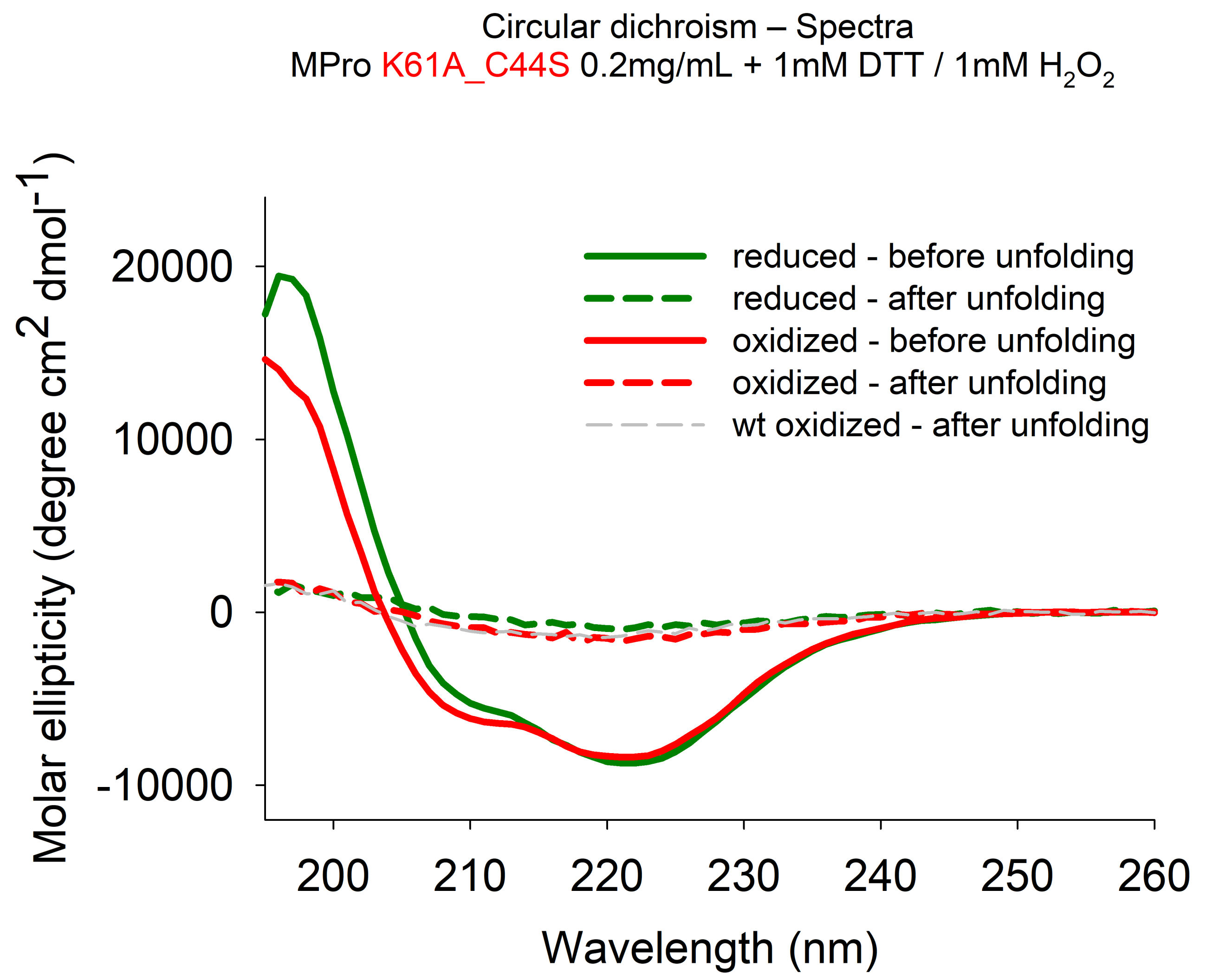

Circular dichroism – Spectra
MPro K61A_C44S 0.2mg/mL + 1mM DTT / 1mM H2O2
Molar ellipticity (degree cm2 dmol-1)
20000
10000
0
-10000
200
210
220
230
240
250
260
Wavelength (nm)
reduced - before unfolding
reduced - after unfolding
oxidized - before unfolding
oxidized - after unfolding
wt oxidized - after unfolding

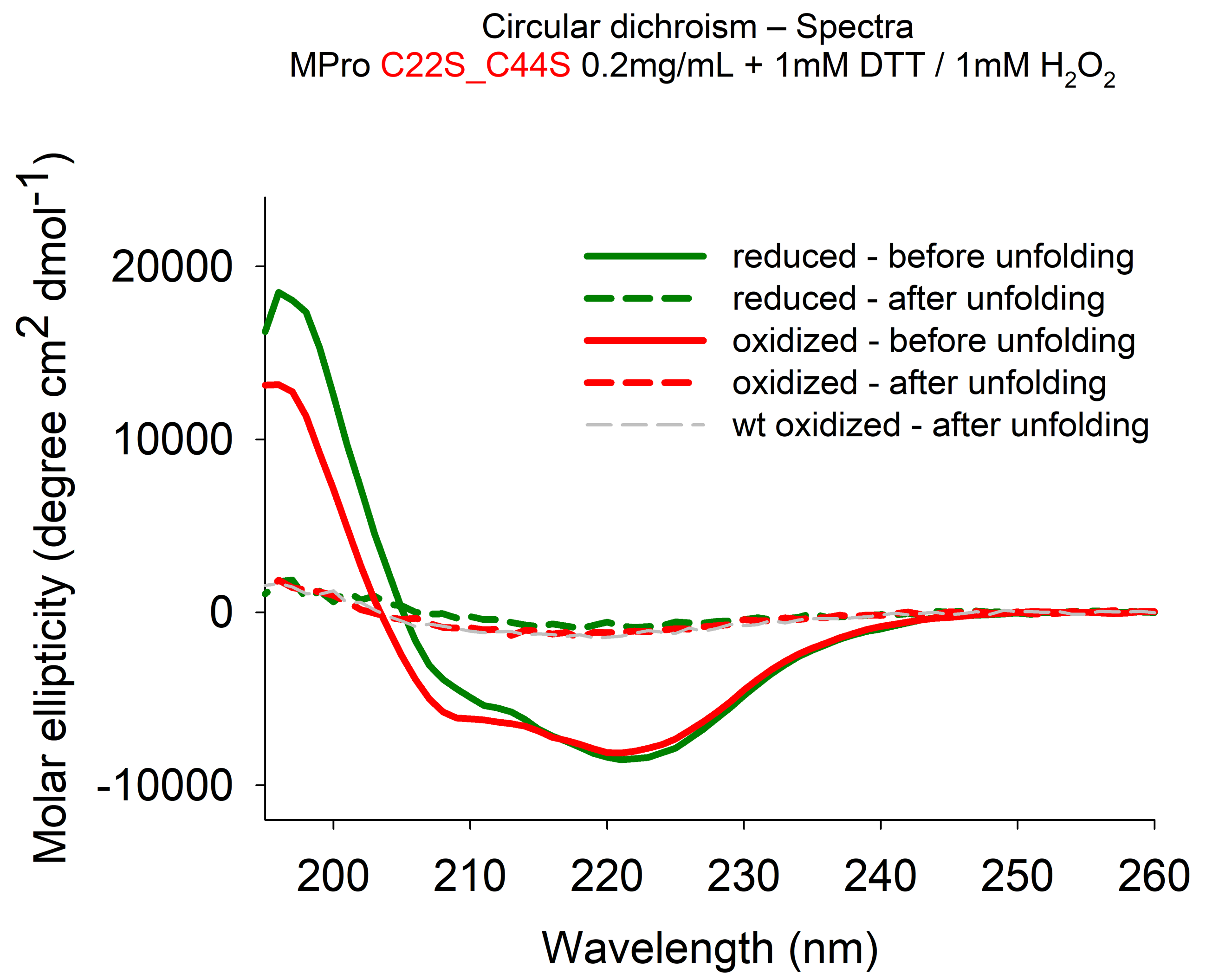

Circular dichroism – Spectra
MPro C22S_C44S 0.2mg/mL + 1mM DTT / 1mM $H_2O_2$
Molar ellipticity (degree cm$^2$ dmol$^{-1}$)
20000
10000
0
-10000
200
210
220
230
240
250
260
Wavelength (nm)
reduced - before unfolding
reduced - after unfolding
oxidized - before unfolding
oxidized - after unfolding
wt oxidized - after unfolding

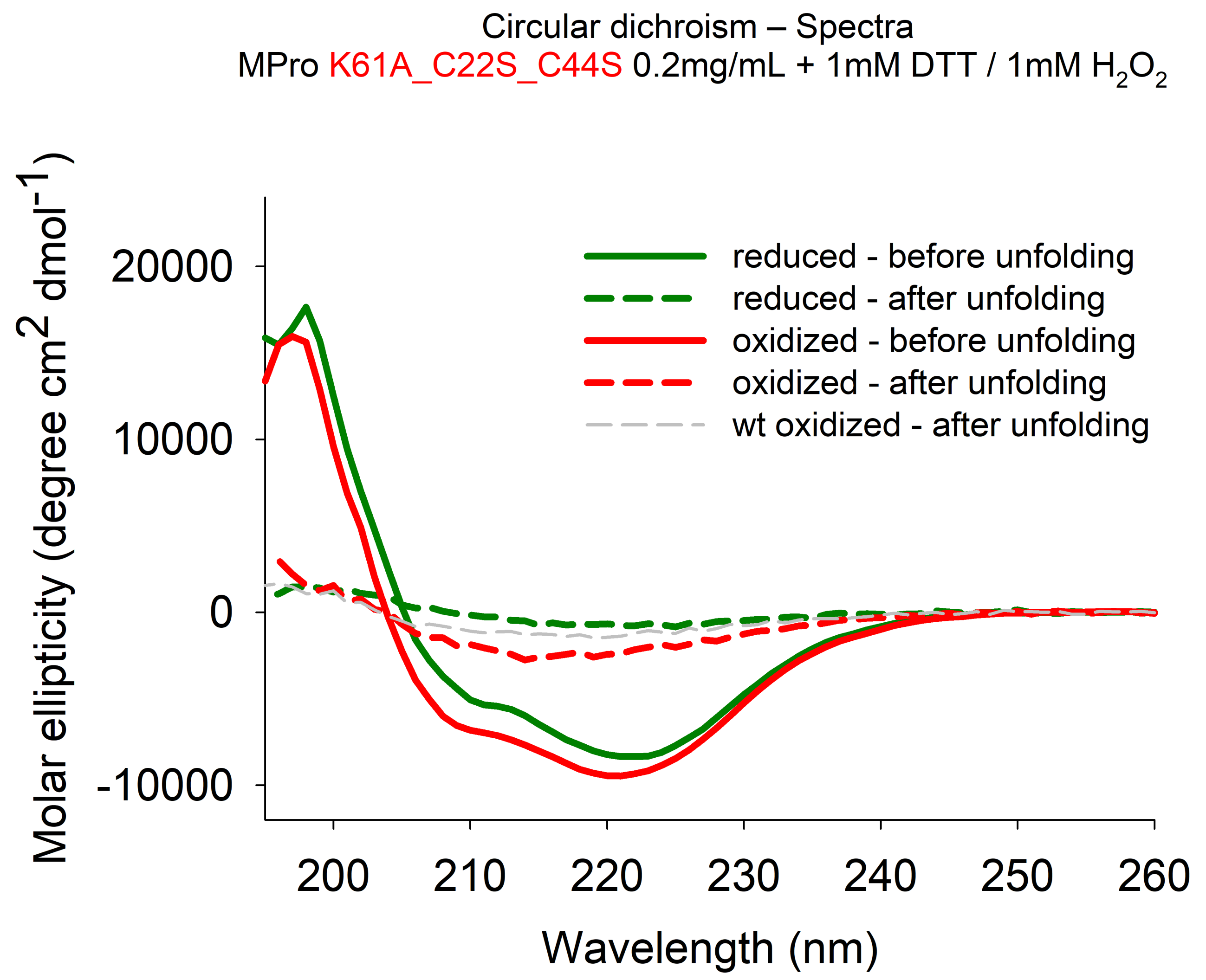

Circular dichroism – Spectra
MPro K61A_C22S_C44S 0.2mg/mL + 1mM DTT / 1mM $H_2O_2$
reduced - before unfolding
reduced - after unfolding
oxidized - before unfolding
oxidized - after unfolding
wt oxidized - after unfolding
Molar ellipticity (degree cm$^2$ dmol$^{-1}$)
20000
10000
0
-10000
200
210
220
230
240
250
260
Wavelength (nm)

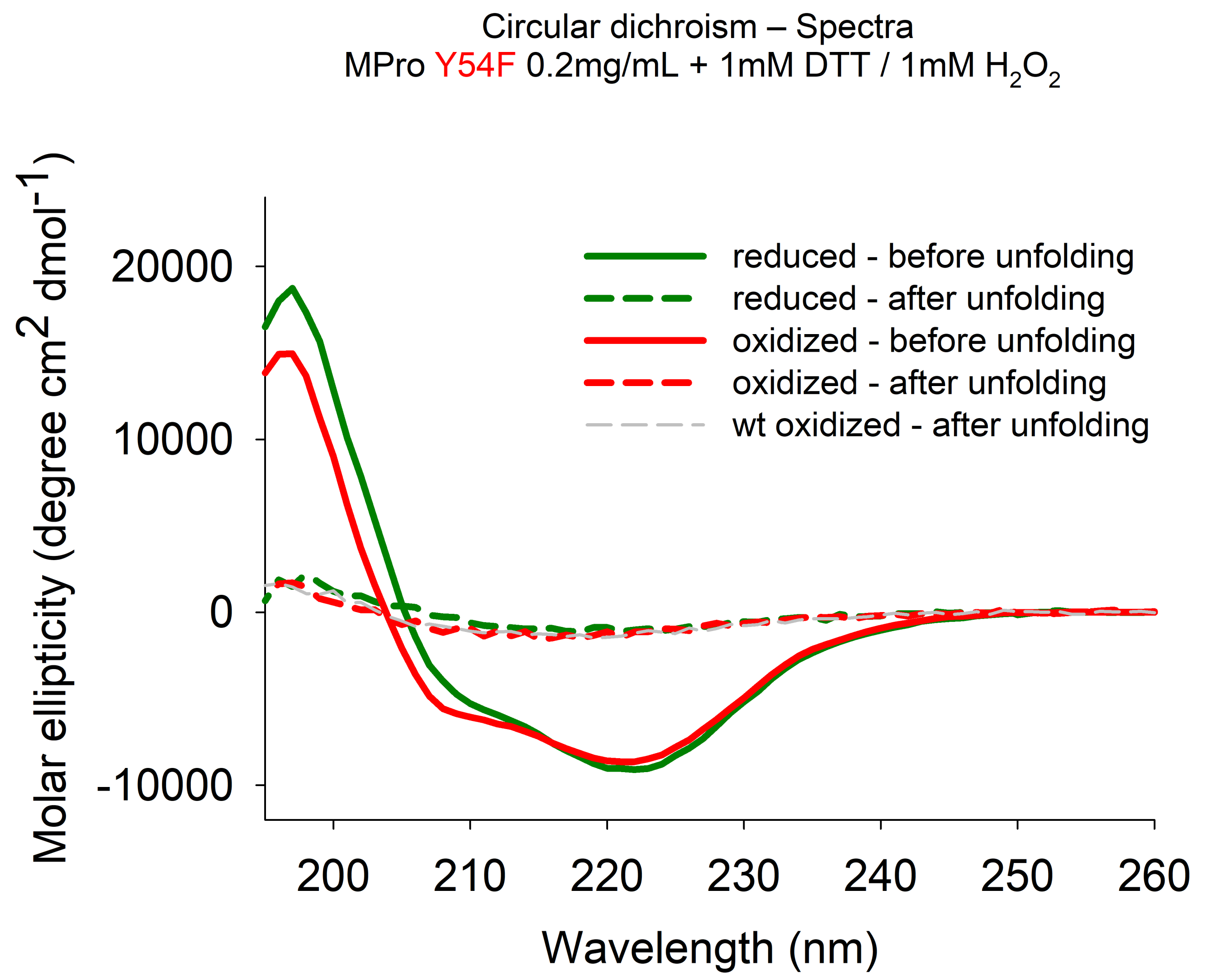

Circular dichroism – Spectra
MPro Y54F 0.2mg/mL + 1mM DTT / 1mM $H_2O_2$
Molar ellipticity (degree cm$^2$ dmol$^{-1}$)
20000
10000
0
-10000
200
210
220
230
240
250
260
Wavelength (nm)
reduced - before unfolding
reduced - after unfolding
oxidized - before unfolding
oxidized - after unfolding
wt oxidized - after unfolding
