## Appendix 3 - Unfolding data for "Redox regulation of the SARS-CoV-2 main protease provides new opportunities for drug design"

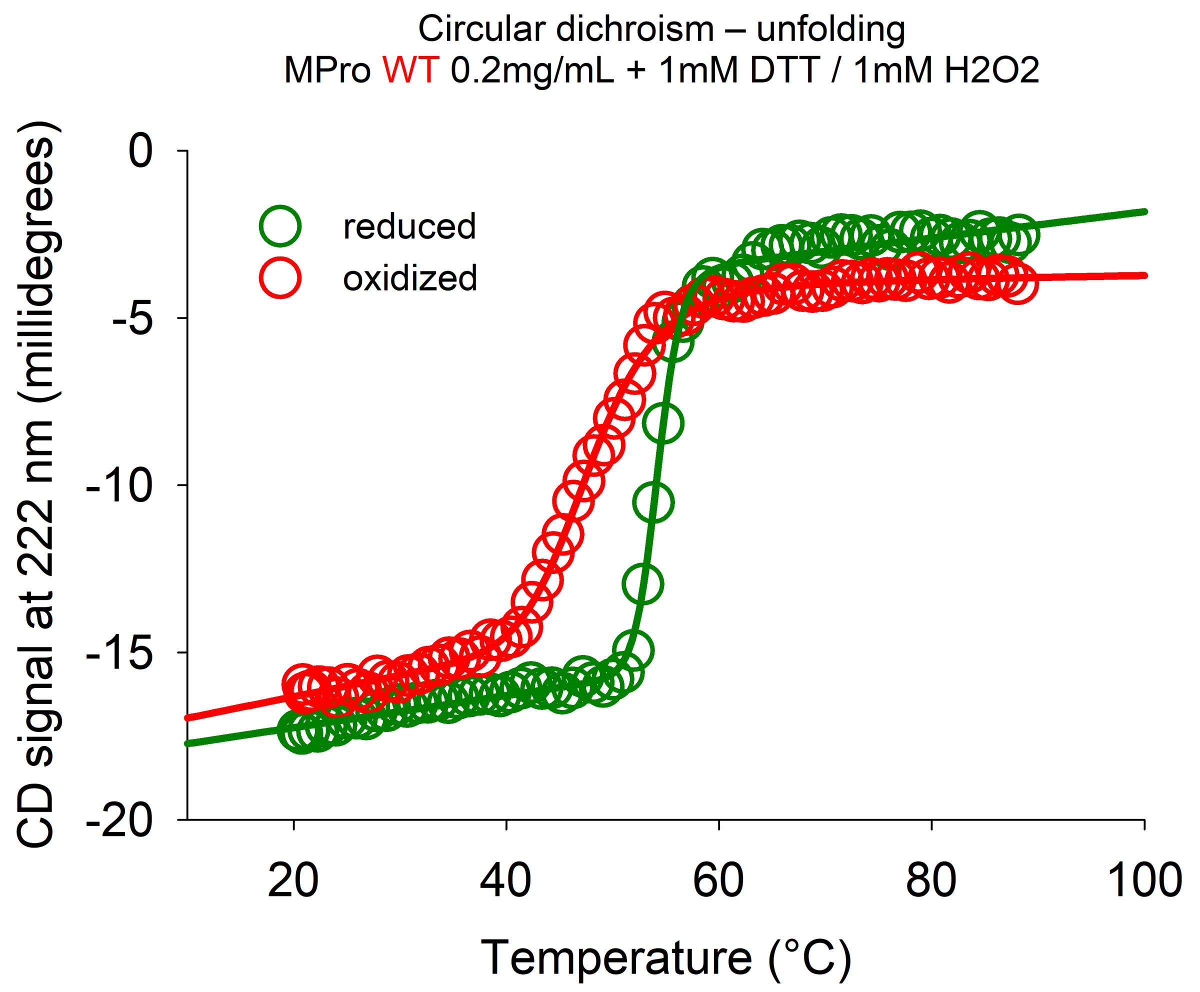
Circular dichroism – unfolding
MPro WT 0.2mg/mL + 1mM DTT / 1mM H2O2
reduced
oxidized
CD signal at 222 nm (millidegrees)
0
-5
-10
-15
-20
20
40
60
80
100
Temperature (°C)

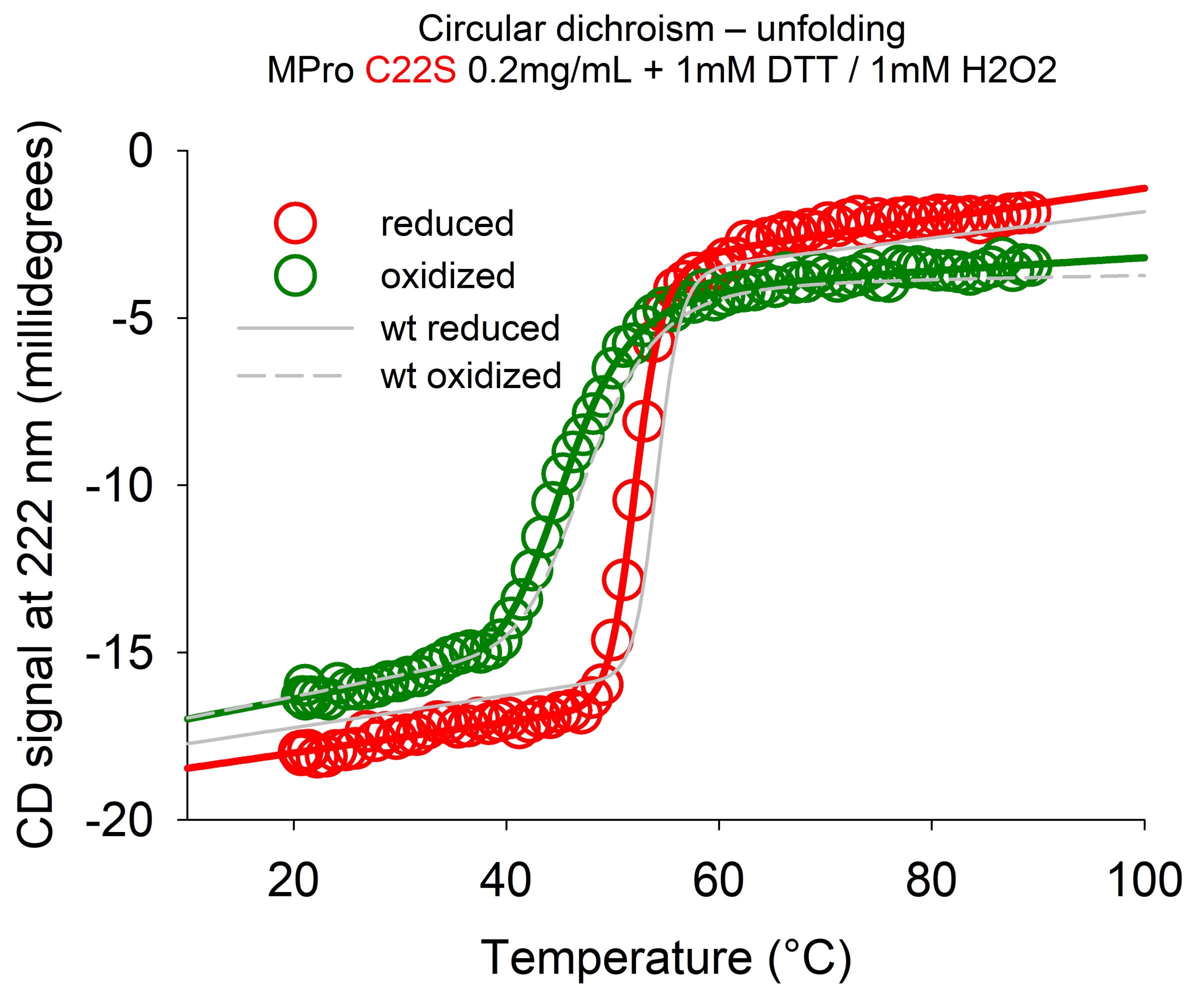
Circular dichroism – unfolding
MPro C22S 0.2mg/mL + 1mM DTT / 1mM H2O2
reduced
oxidized
wt reduced
wt oxidized
CD signal at 222 nm (millidegrees)
0
-5
-10
-15
-20
20
40
60
80
100
Temperature (°C)

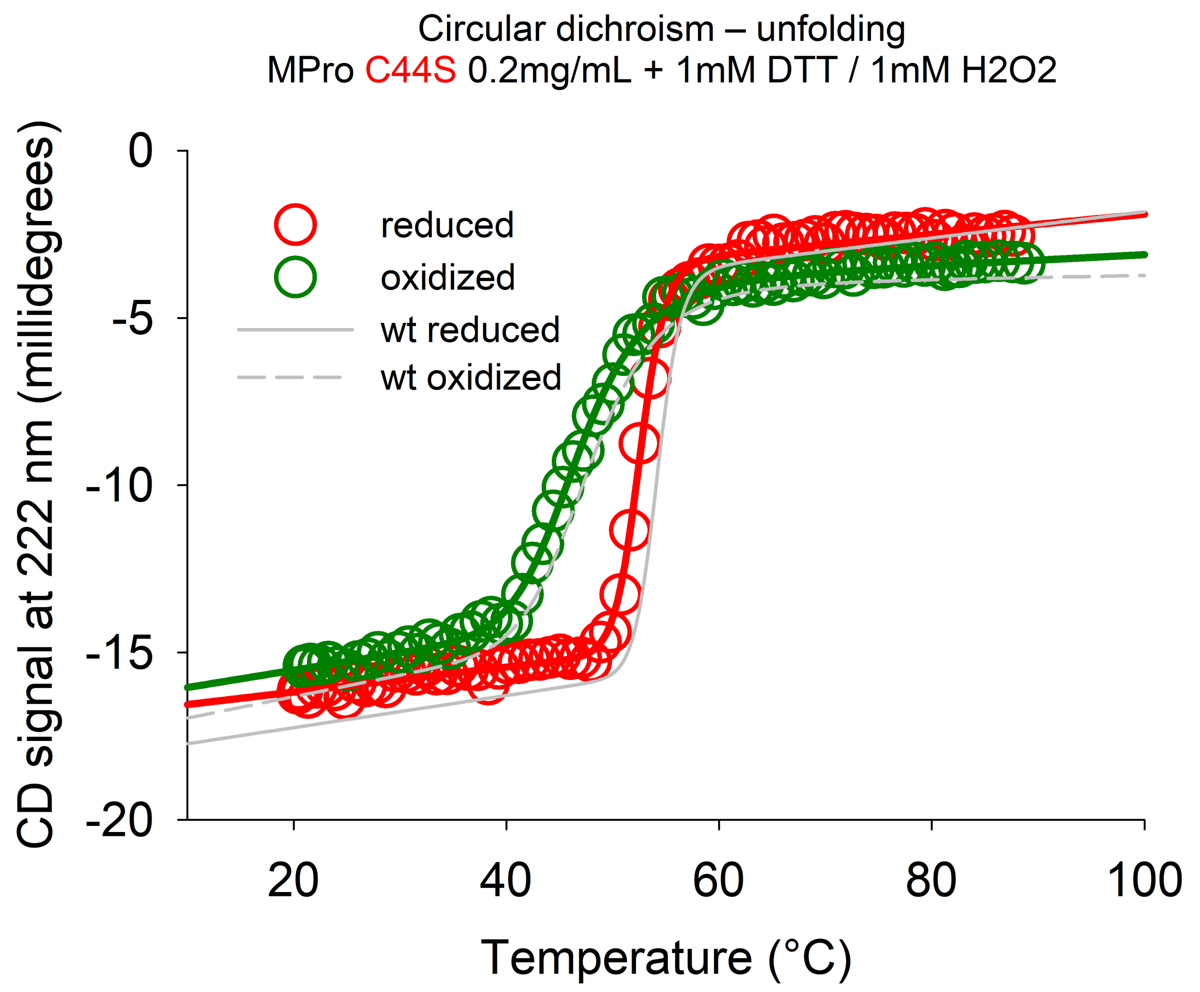
Circular dichroism – unfolding
MPro C44S 0.2mg/mL + 1mM DTT / 1mM H2O2
reduced
oxidized
wt reduced
wt oxidized
CD signal at 222 nm (millidegrees)
0
-5
-10
-15
-20
20
40
60
80
100
Temperature (°C)

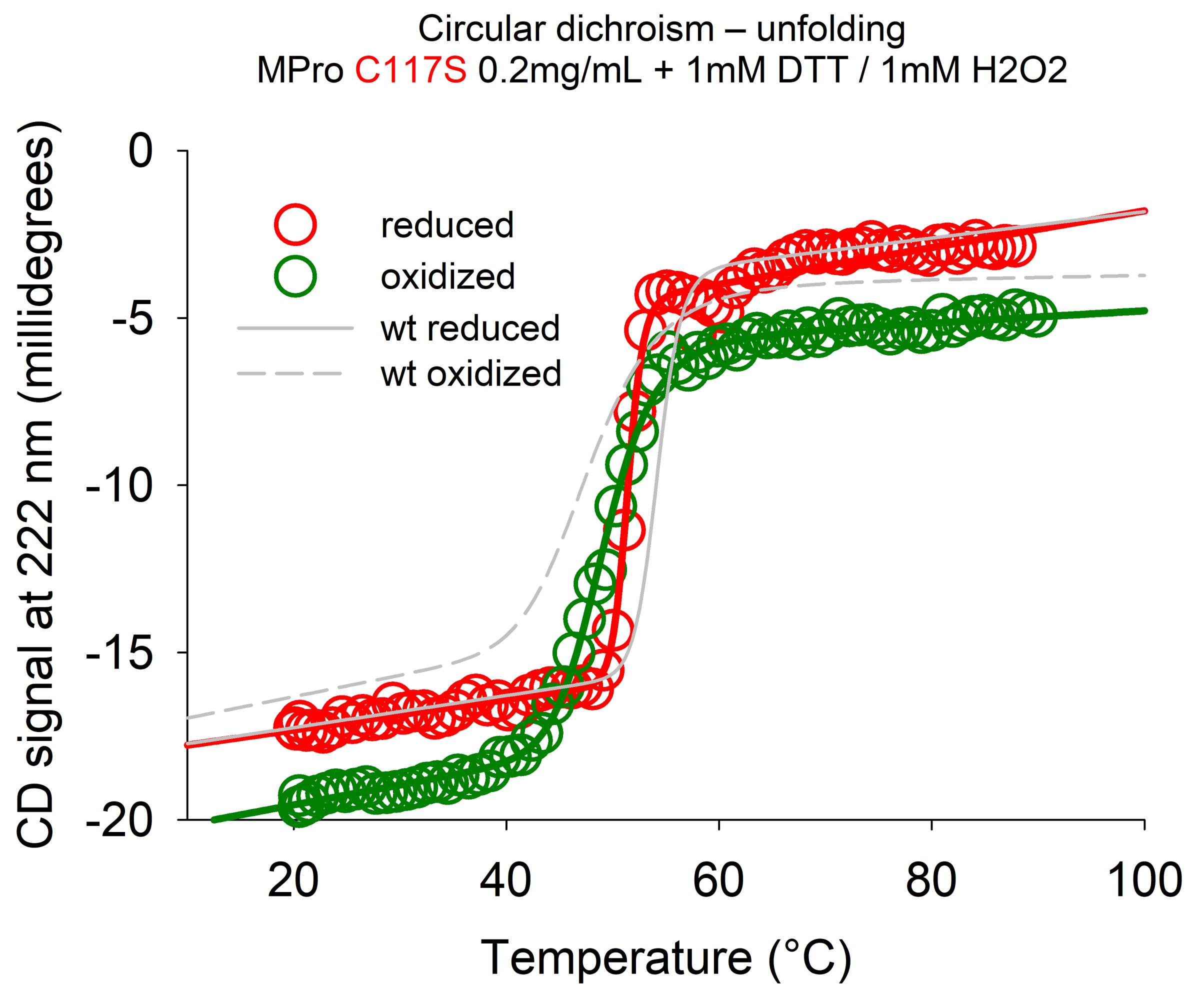
Circular dichroism – unfolding
MPro C117S 0.2mg/mL + 1mM DTT / 1mM H2O2
reduced
oxidized
wt reduced
wt oxidized
CD signal at 222 nm (millidegrees)
0
-5
-10
-15
-20
20
40
60
80
100
Temperature (°C)

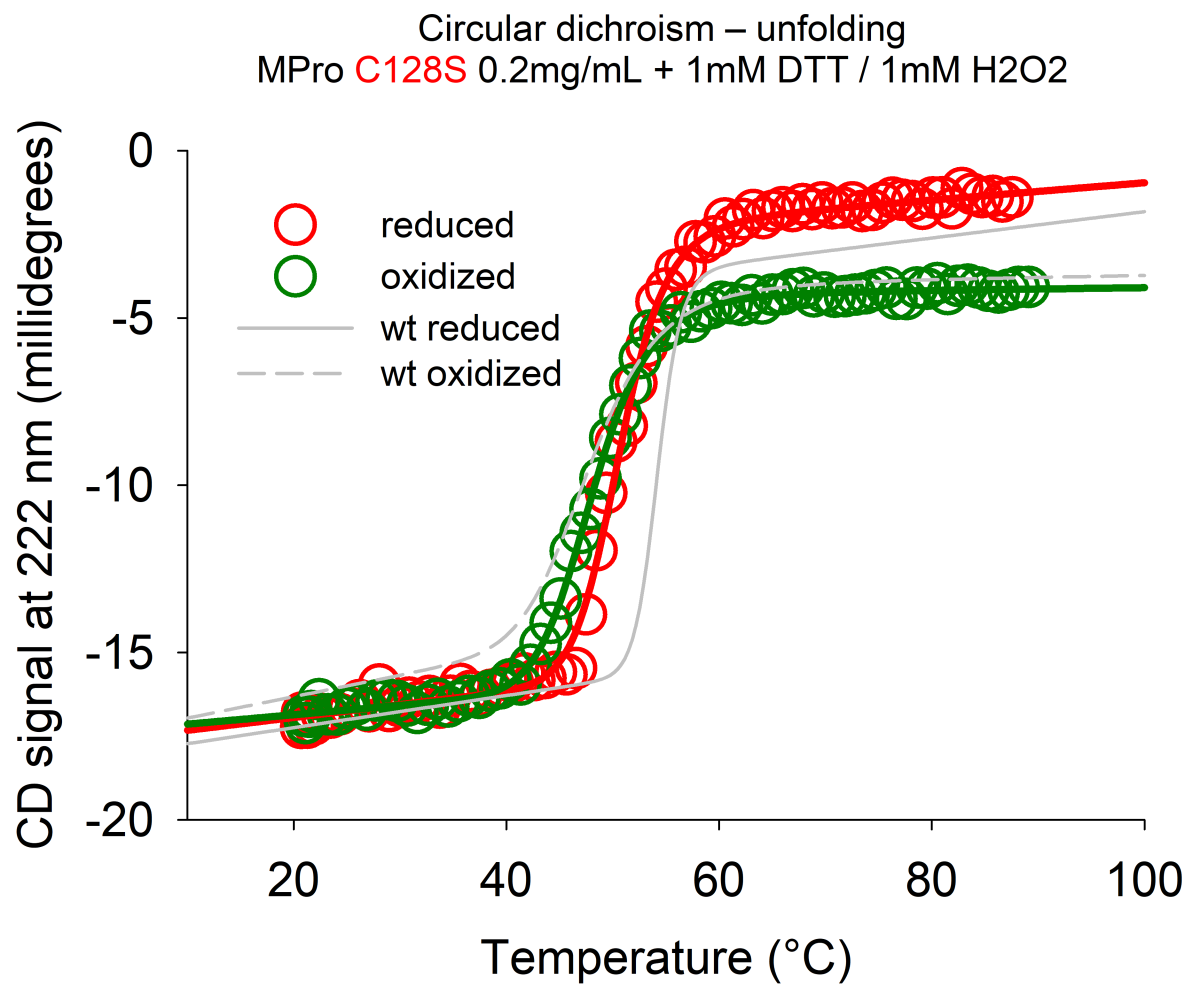
Circular dichroism – unfolding
MPro C128S 0.2mg/mL + 1mM DTT / 1mM H2O2
reduced
oxidized
wt reduced
wt oxidized
CD signal at 222 nm (millidegrees)
0
-5
-10
-15
-20
20
40
60
80
100
Temperature (°C)

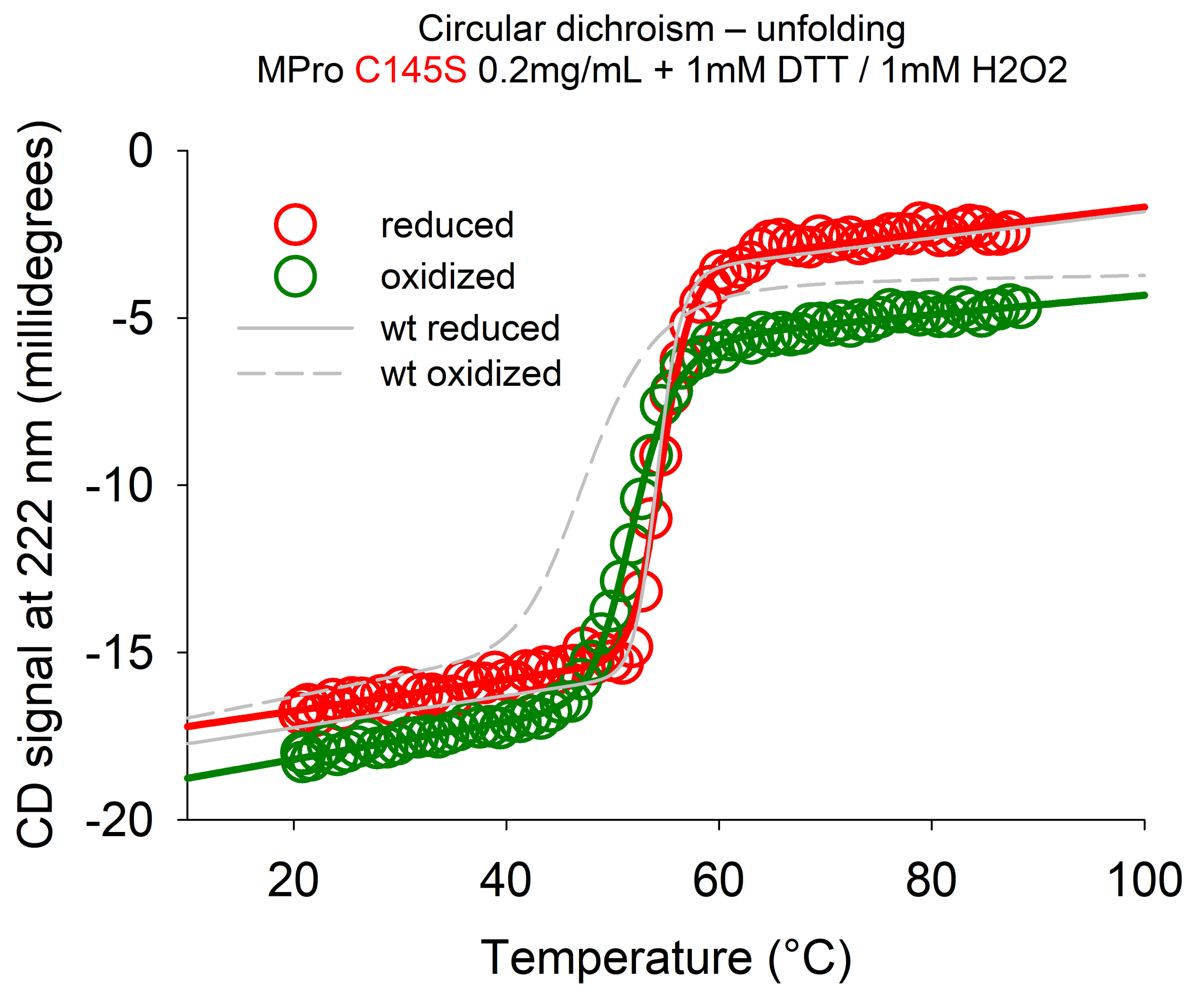

Circular dichroism – unfolding
MPro C145S 0.2mg/mL + 1mM DTT / 1mM H2O2
reduced
oxidized
wt reduced
wt oxidized
0
-5
-10
-15
-20
20
40
60
80
100
CD signal at 222 nm (millidegrees)
Temperature (°C)

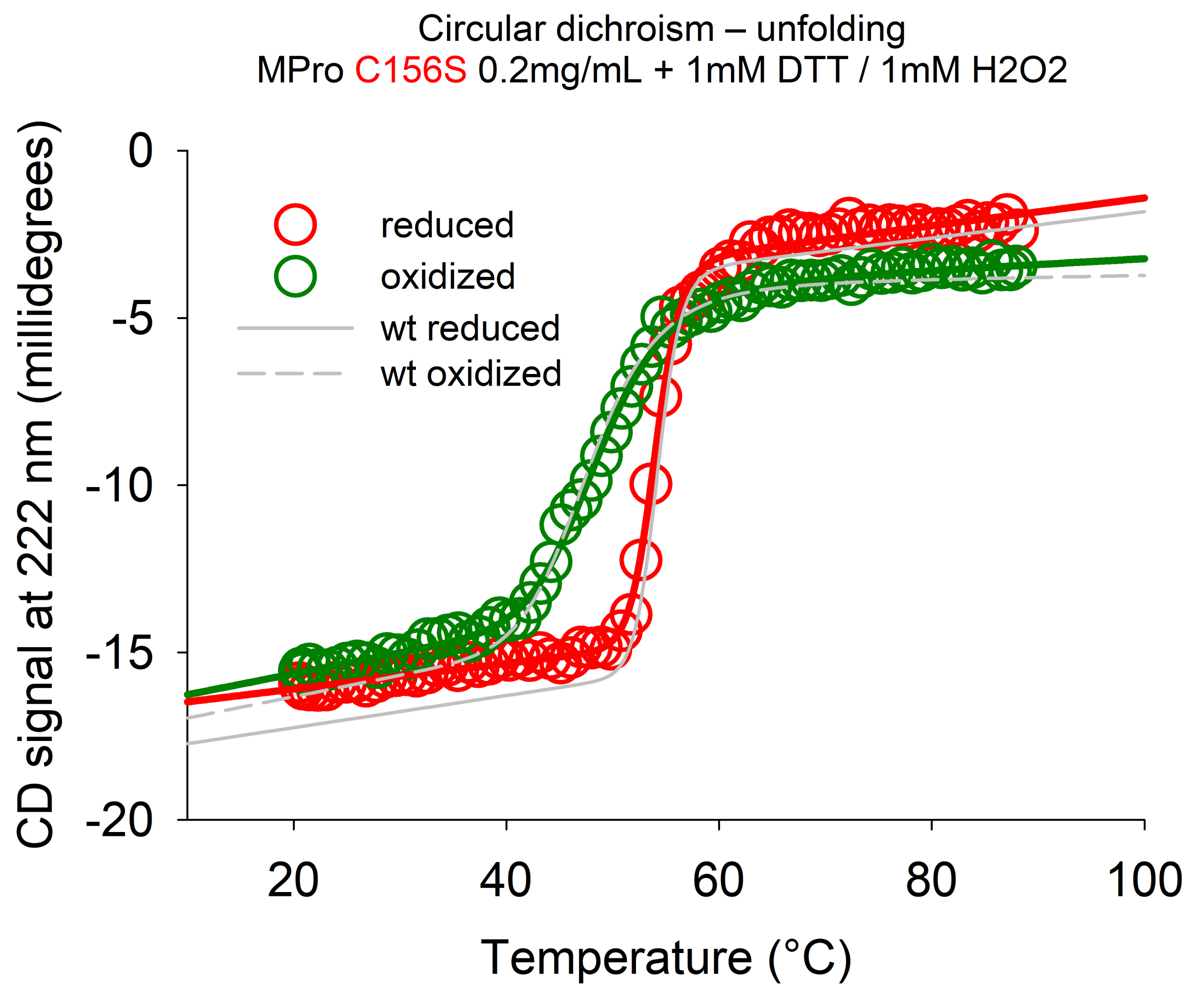
Circular dichroism – unfolding
MPro C156S 0.2mg/mL + 1mM DTT / 1mM H2O2
reduced
oxidized
wt reduced
wt oxidized
CD signal at 222 nm (millidegrees)
0
-5
-10
-15
-20
20
40
60
80
100
Temperature (°C)

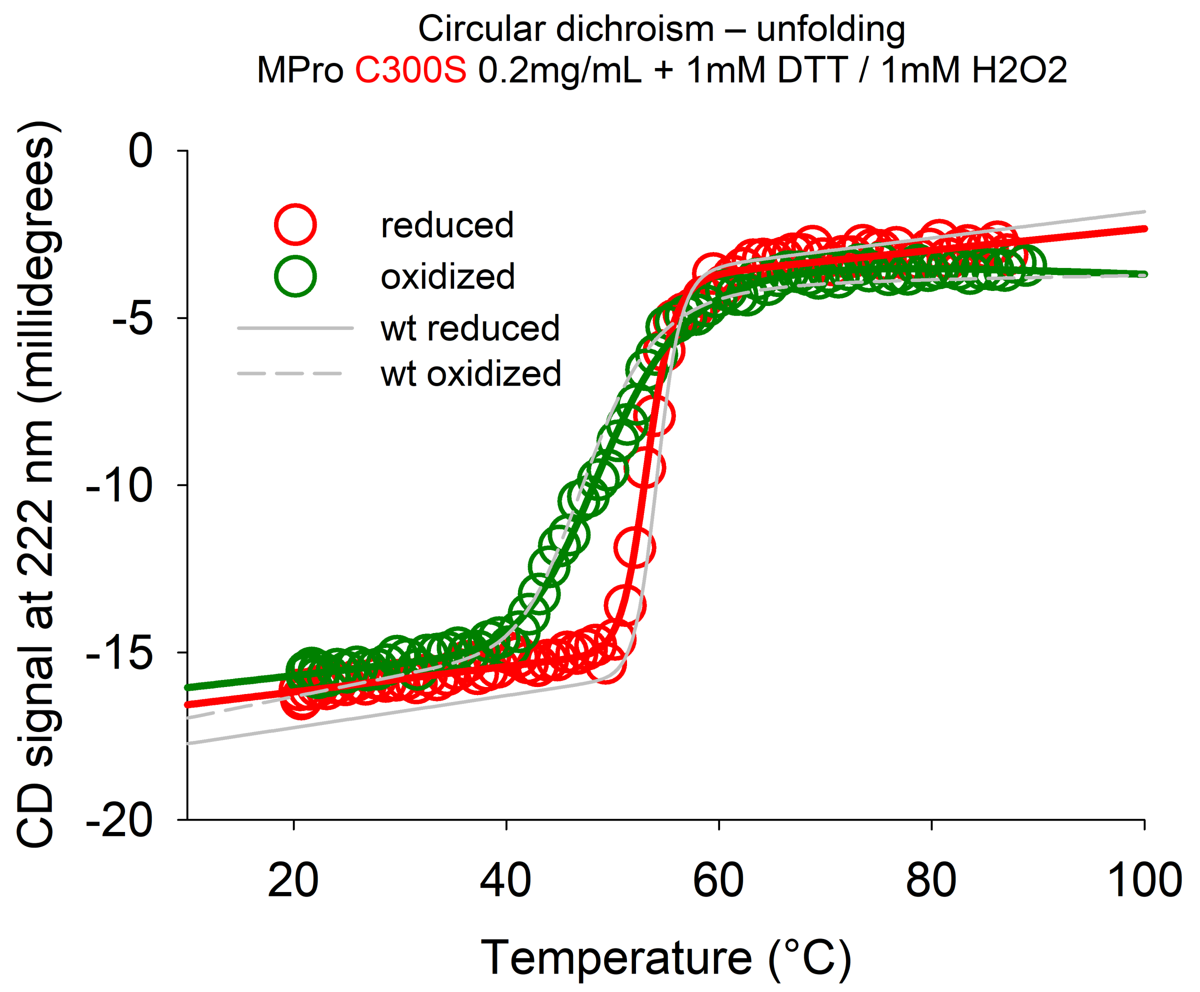
Circular dichroism – unfolding
MPro C300S 0.2mg/mL + 1mM DTT / 1mM H2O2
reduced
oxidized
wt reduced
wt oxidized
0
-5
-10
-15
-20
20
40
60
80
100
CD signal at 222 nm (millidegrees)
Temperature (°C)

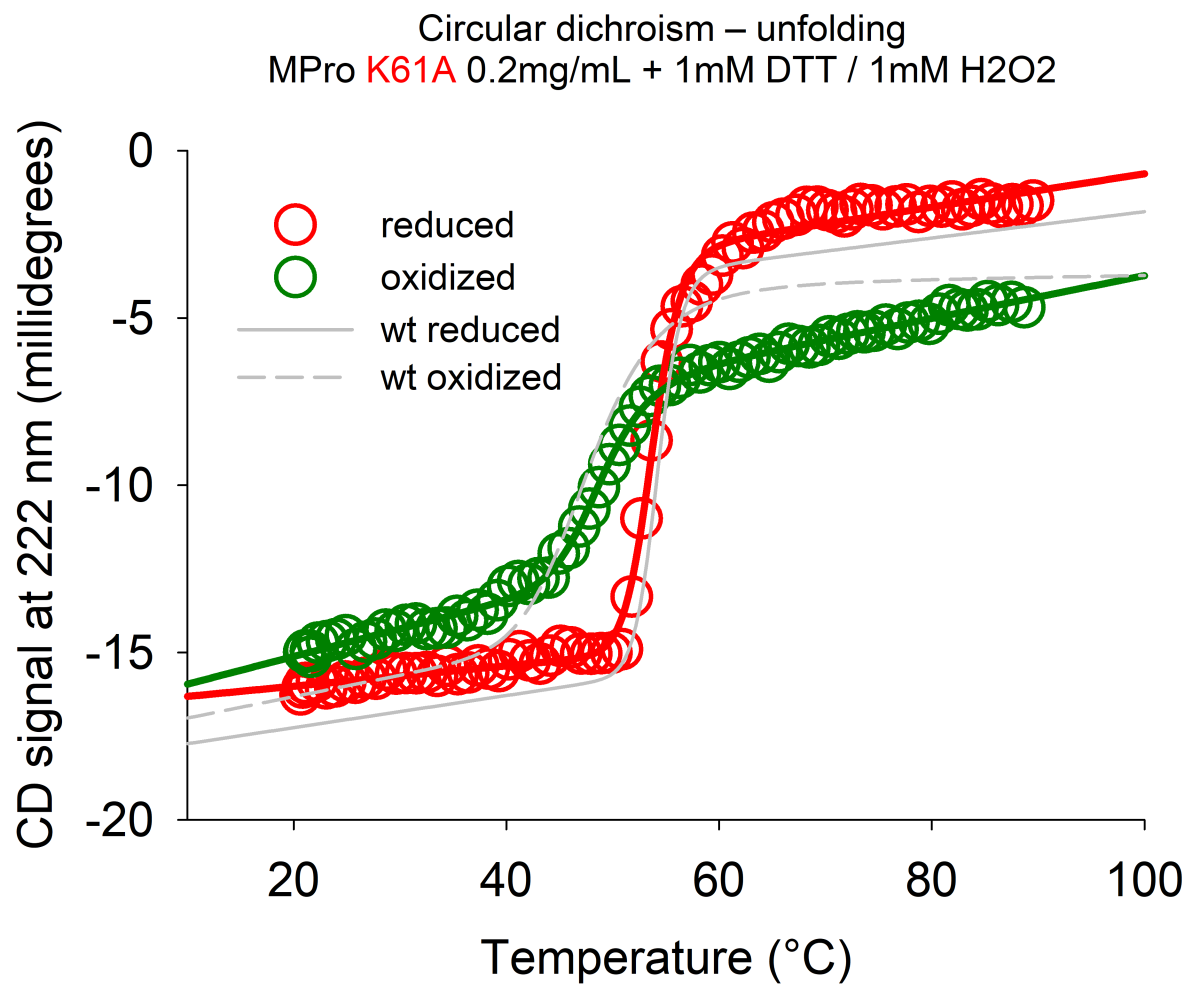

Circular dichroism – unfolding
MPro K61A 0.2mg/mL + 1mM DTT / 1mM H2O2
CD signal at 222 nm (millidegrees)
0
-5
-10
-15
-20
20
40
60
80
100
Temperature (°C)
reduced
oxidized
wt reduced
wt oxidized

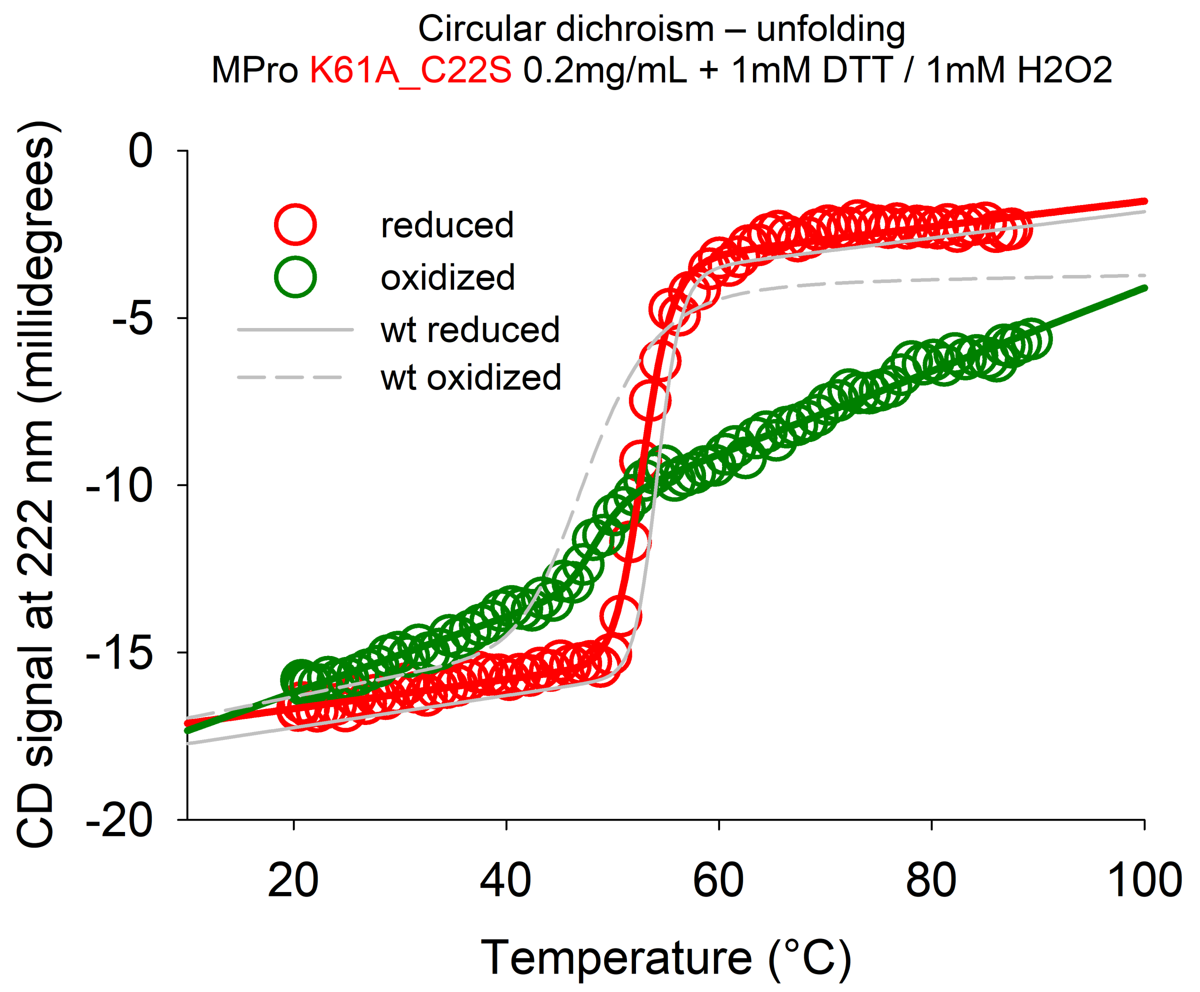
Circular dichroism – unfolding
MPro K61A_C22S 0.2mg/mL + 1mM DTT / 1mM H2O2
reduced
oxidized
wt reduced
wt oxidized
CD signal at 222 nm (millidegrees)
0
-5
-10
-15
-20
20
40
60
80
100
Temperature (°C)

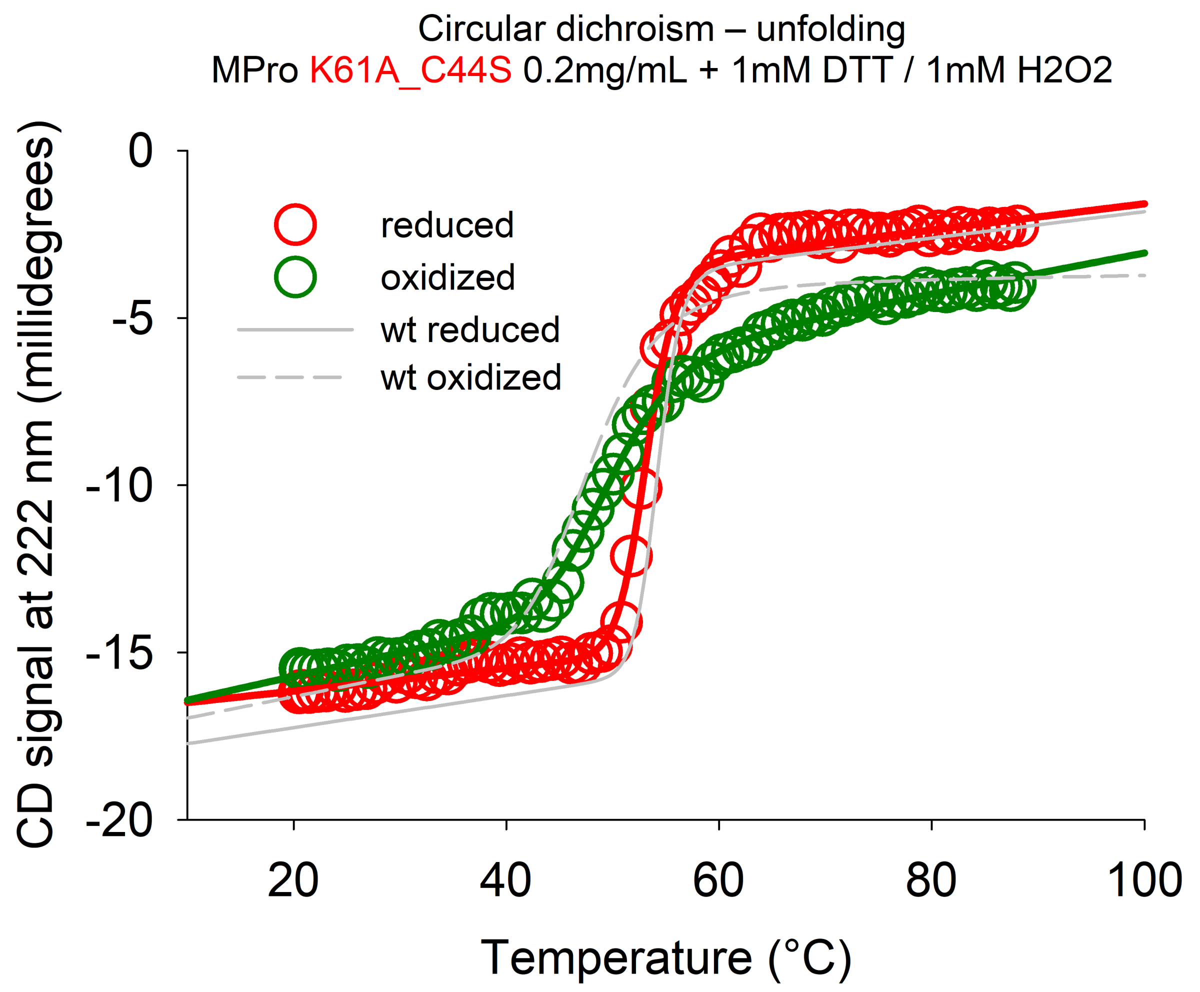
Circular dichroism – unfolding
MPro K61A_C44S 0.2mg/mL + 1mM DTT / 1mM H2O2
reduced
oxidized
wt reduced
wt oxidized
CD signal at 222 nm (millidegrees)
0
-5
-10
-15
-20
20
40
60
80
100
Temperature (°C)

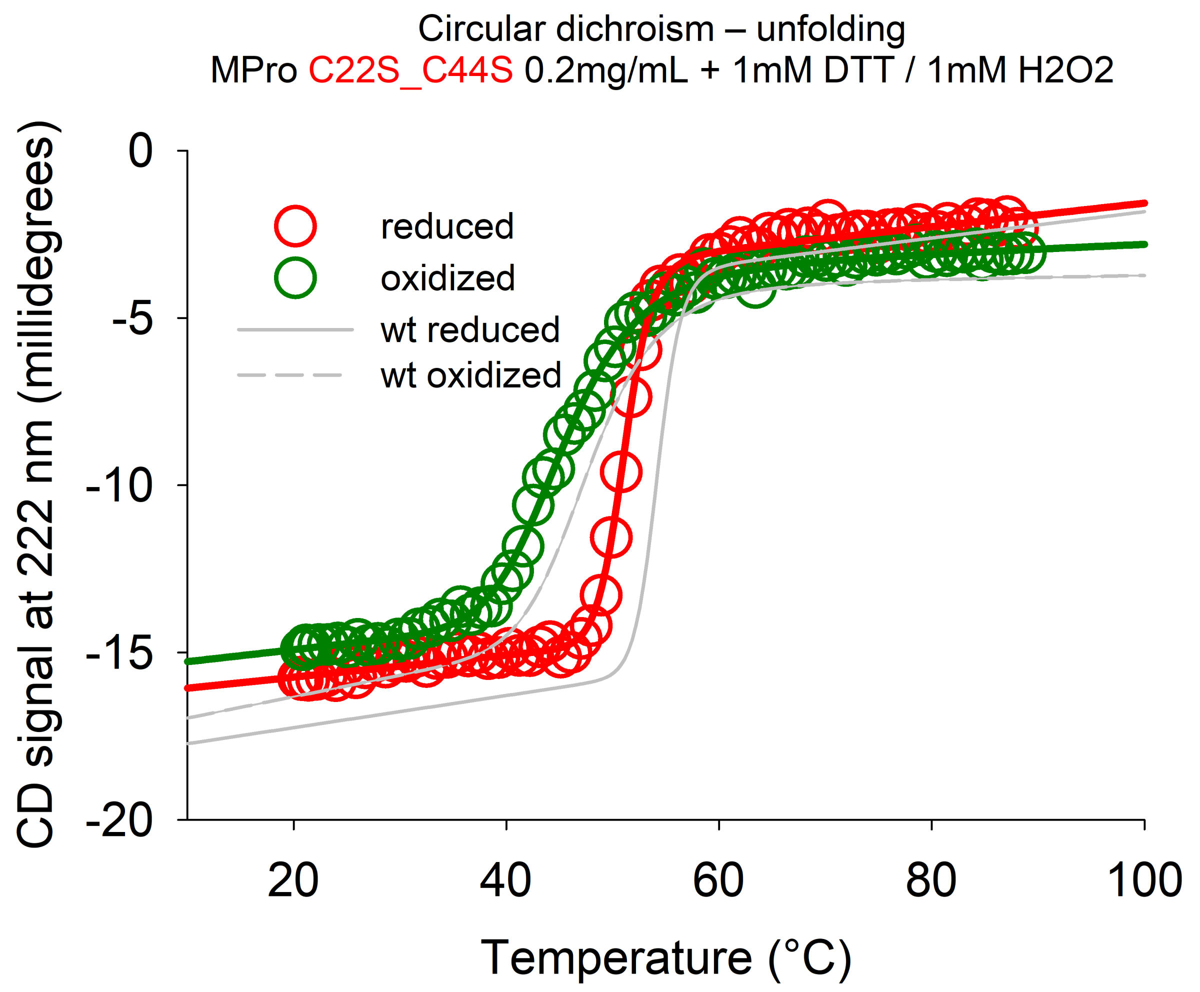
Circular dichroism – unfolding
MPro C22S_C44S 0.2mg/mL + 1mM DTT / 1mM H2O2
reduced
oxidized
wt reduced
wt oxidized
CD signal at 222 nm (millidegrees)
0
-5
-10
-15
-20
20
40
60
80
100
Temperature (°C)

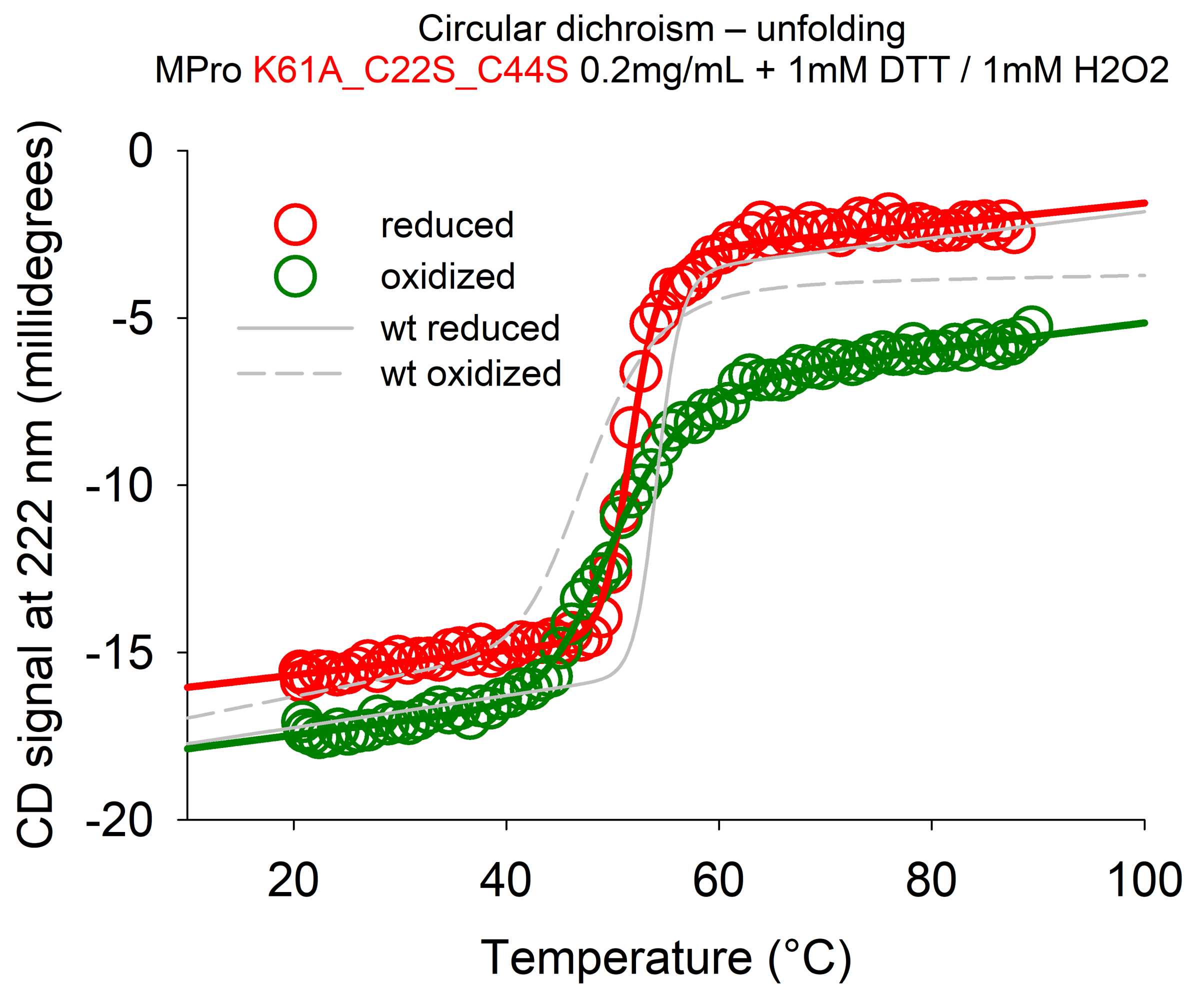

Circular dichroism – unfolding
MPro K61A_C22S_C44S 0.2mg/mL + 1mM DTT / 1mM H2O2
reduced
oxidized
wt reduced
wt oxidized
CD signal at 222 nm (millidegrees)
0
-5
-10
-15
-20
20
40
60
80
100
Temperature (°C)

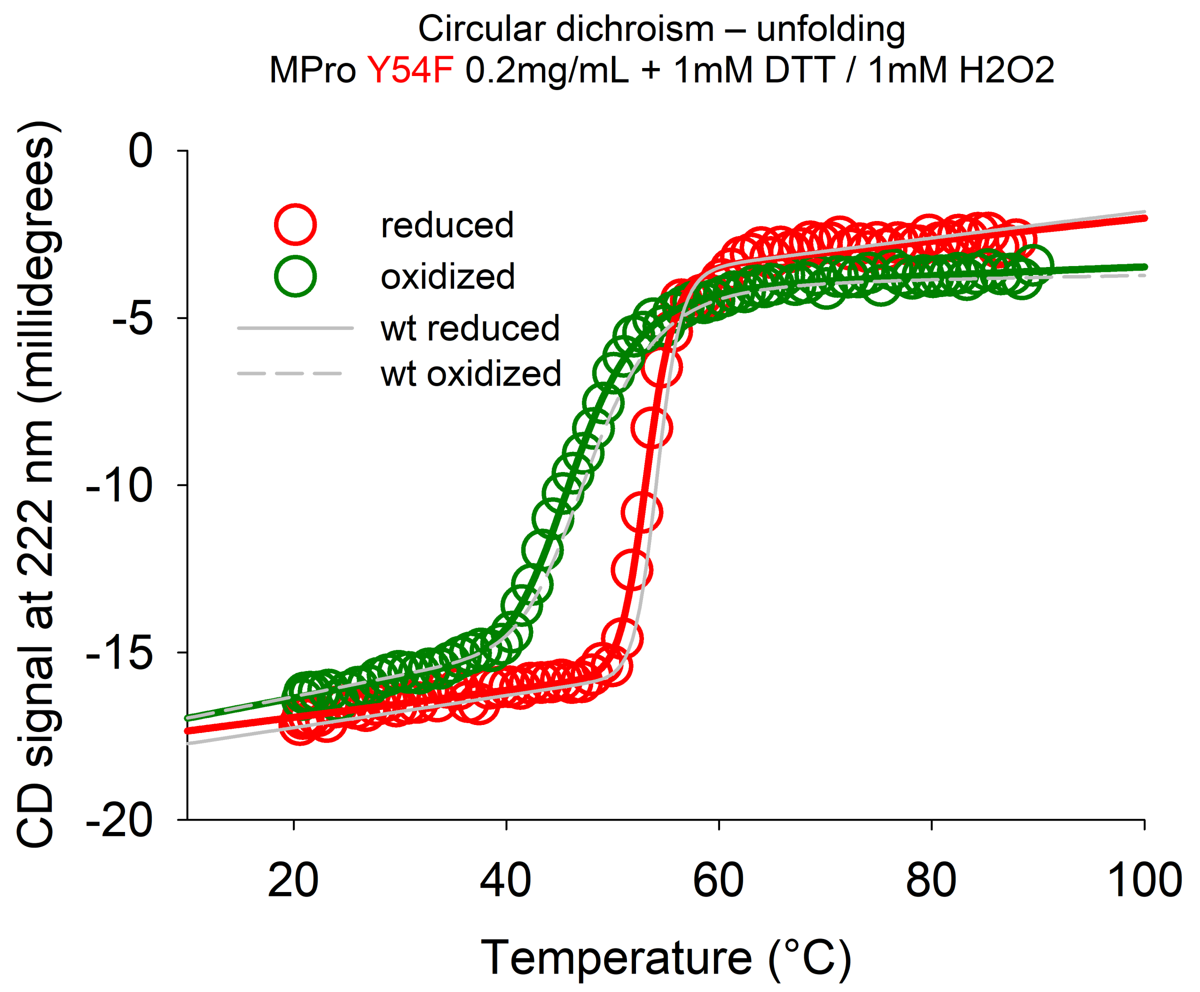

Circular dichroism – unfolding
MPro Y54F 0.2mg/mL + 1mM DTT / 1mM H2O2
reduced
oxidized
wt reduced
wt oxidized
0
-5
-10
-15
-20
20
40
60
80
100
CD signal at 222 nm (millidegrees)
Temperature (°C)
