## Appendix 4 - Crosslinking experiments for "Redox regulation of the SARS-CoV-2 main protease provides new opportunities for drug design"

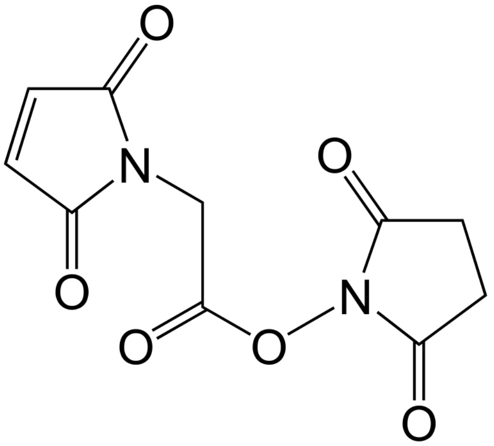

**MAH**
**(Maleimidoacetic acid N-hydroxysuccinimide ester)**

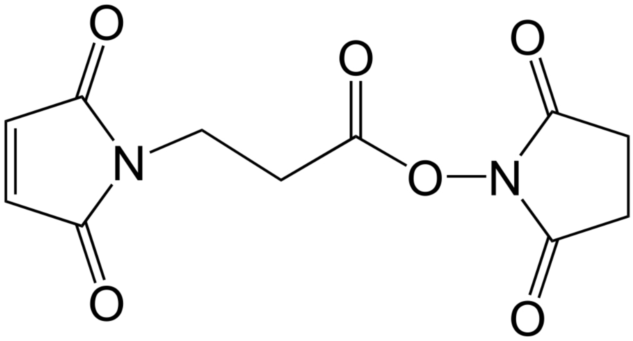

**MPH**
**(3-(Maleimido)propionic acid N-hydroxysuccinimide ester)**

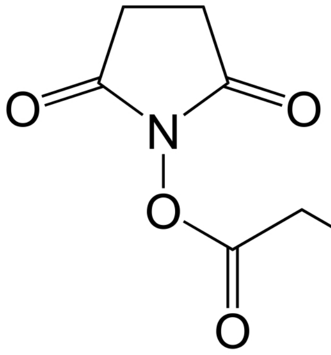

**IANH**
**(Iodoacetic acid N-hydroxysuccinimide ester)**

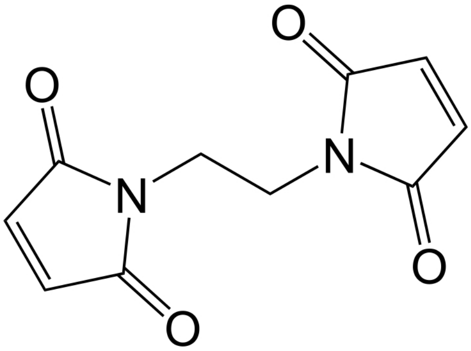

**BMOE**
**(bis-maleimidoethane)**

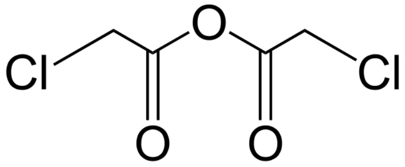

**CAH**
**(Chloroacetic anhydride)**

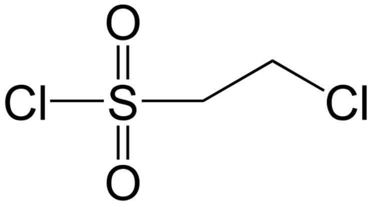

**CSC**
**(2-Chloroethanesulfonyl chloride)**

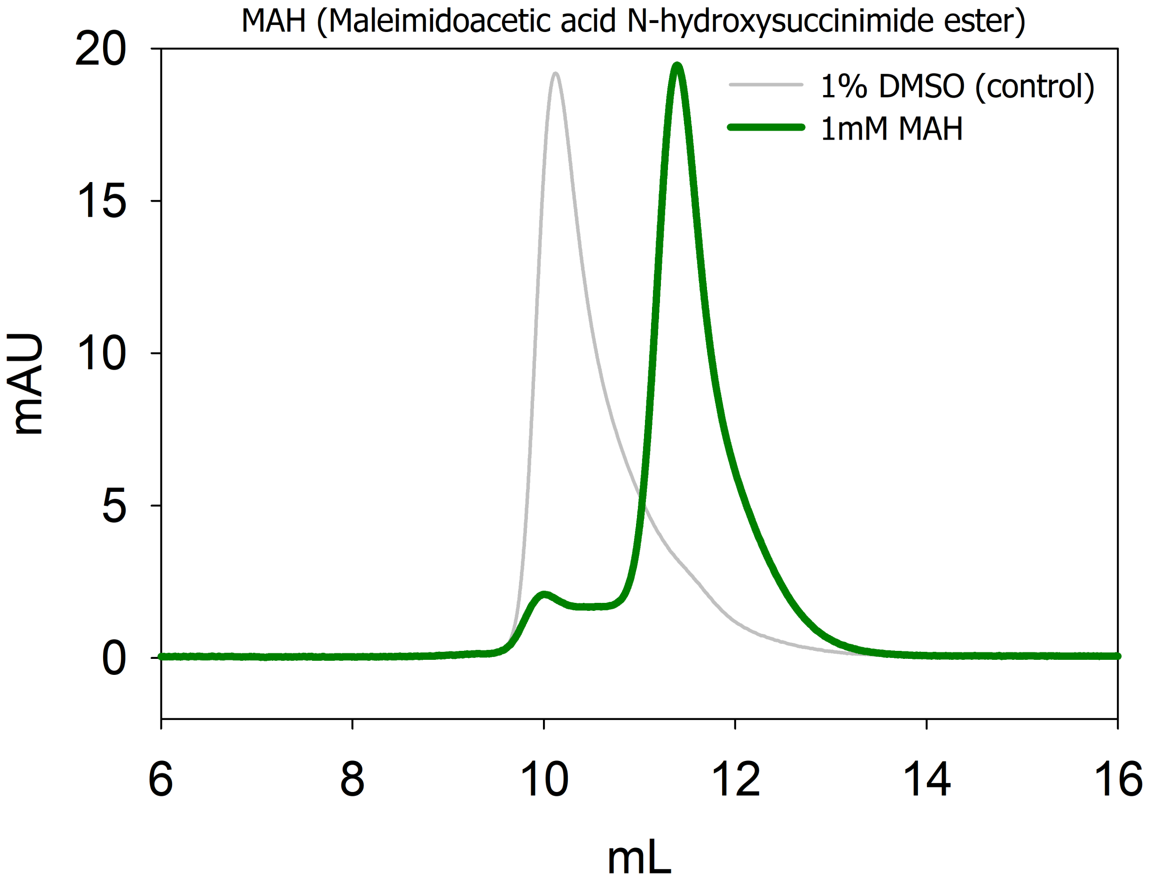
MAH (Maleimidoacetic acid N-hydroxysuccinimide ester)
1% DMSO (control)
1mM MAH
mAU
20
15
10
5
0
6
8
10
12
14
16
mL

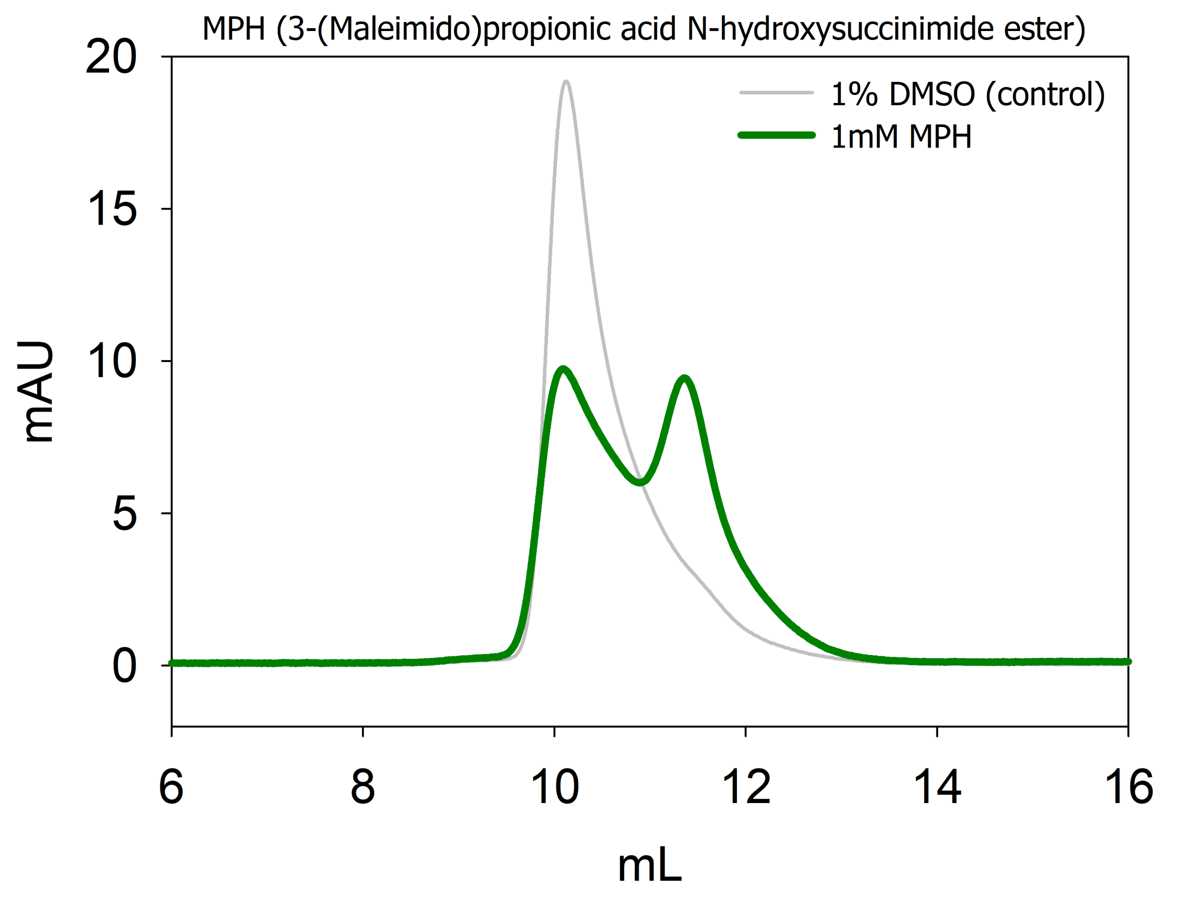
MPH (3-(Maleimido)propionic acid N-hydroxysuccinimide ester)
1% DMSO (control)
1mM MPH
mAU
20
15
10
5
0
6
8
10
12
14
16
mL

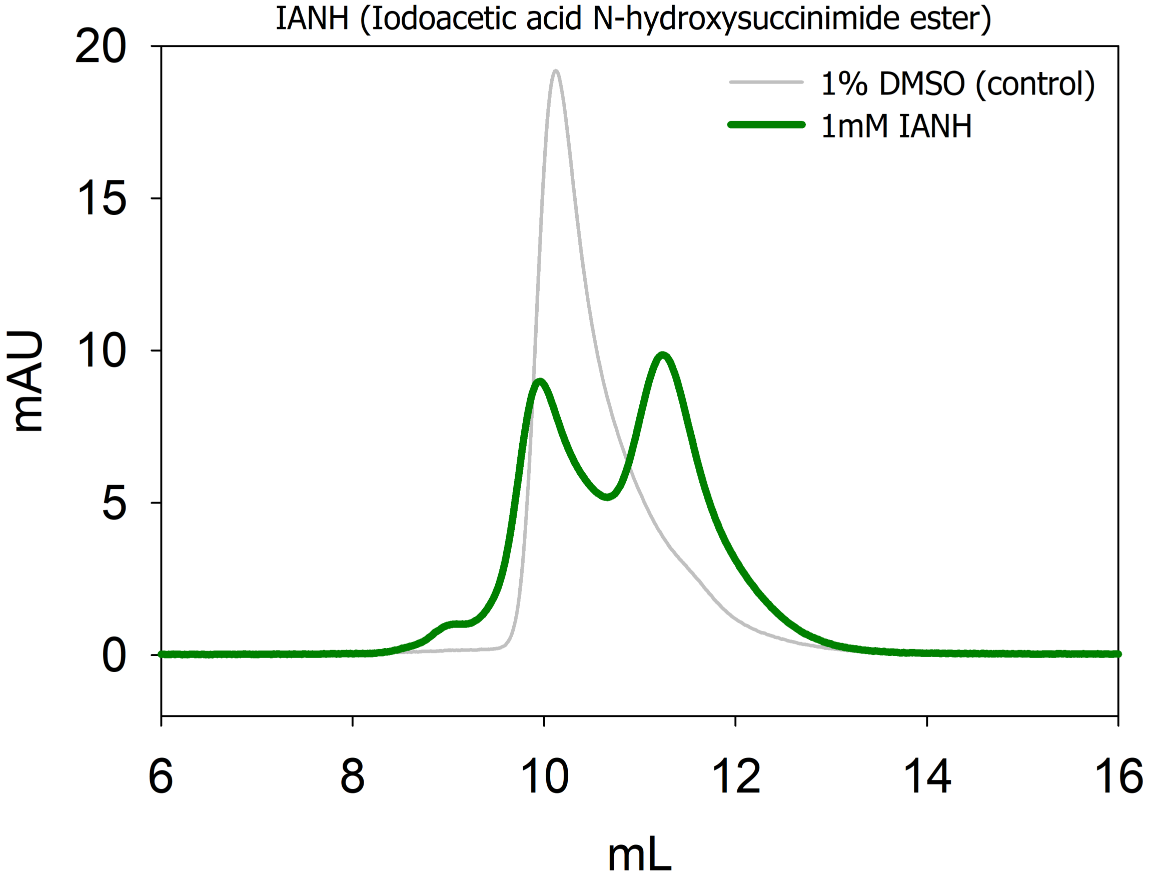
IANH (Iodoacetic acid N-hydroxysuccinimide ester)
1% DMSO (control)
1mM IANH
mAU
20
15
10
5
0
6
8
10
12
14
16
mL

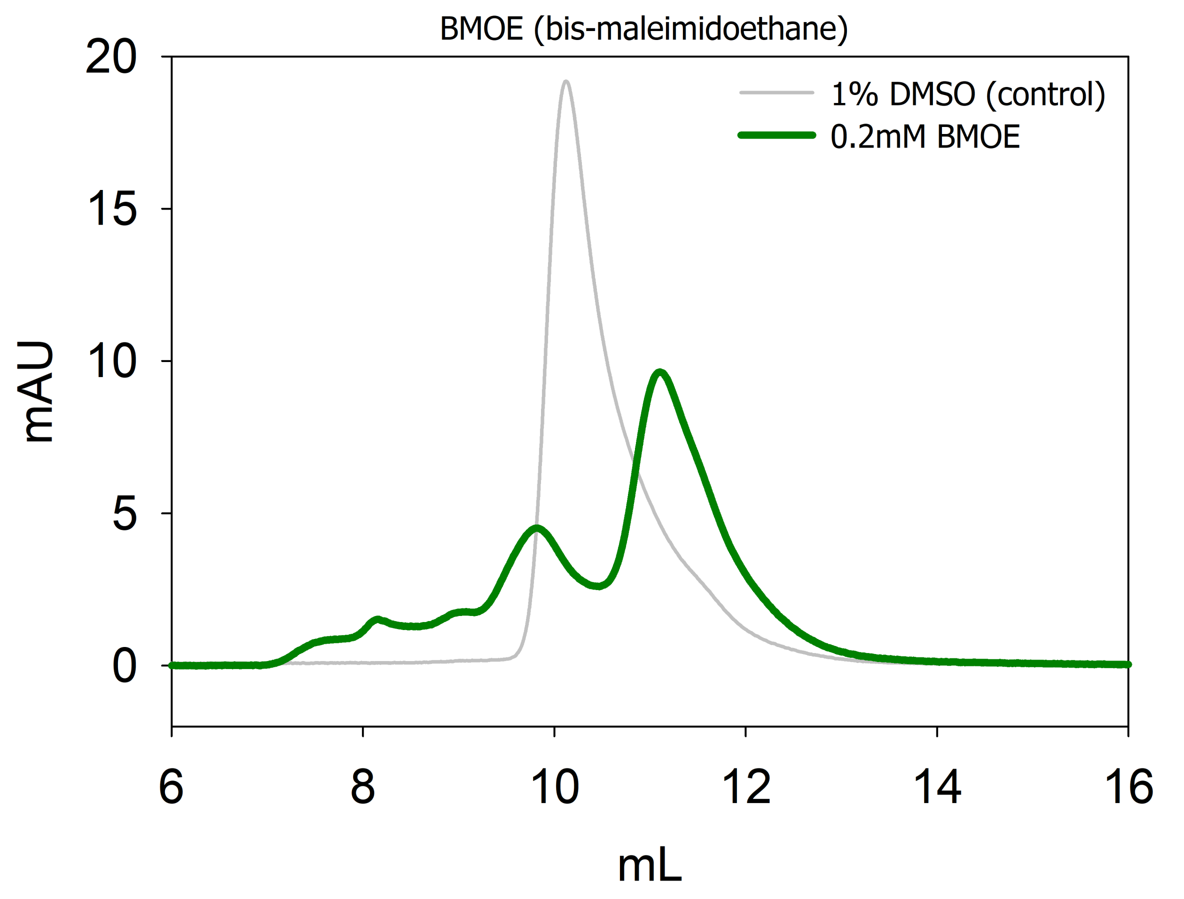
BMOE (bis-maleimidoethane)
1% DMSO (control)
0.2mM BMOE
mAU
20
15
10
5
0
6
8
10
12
14
16
mL

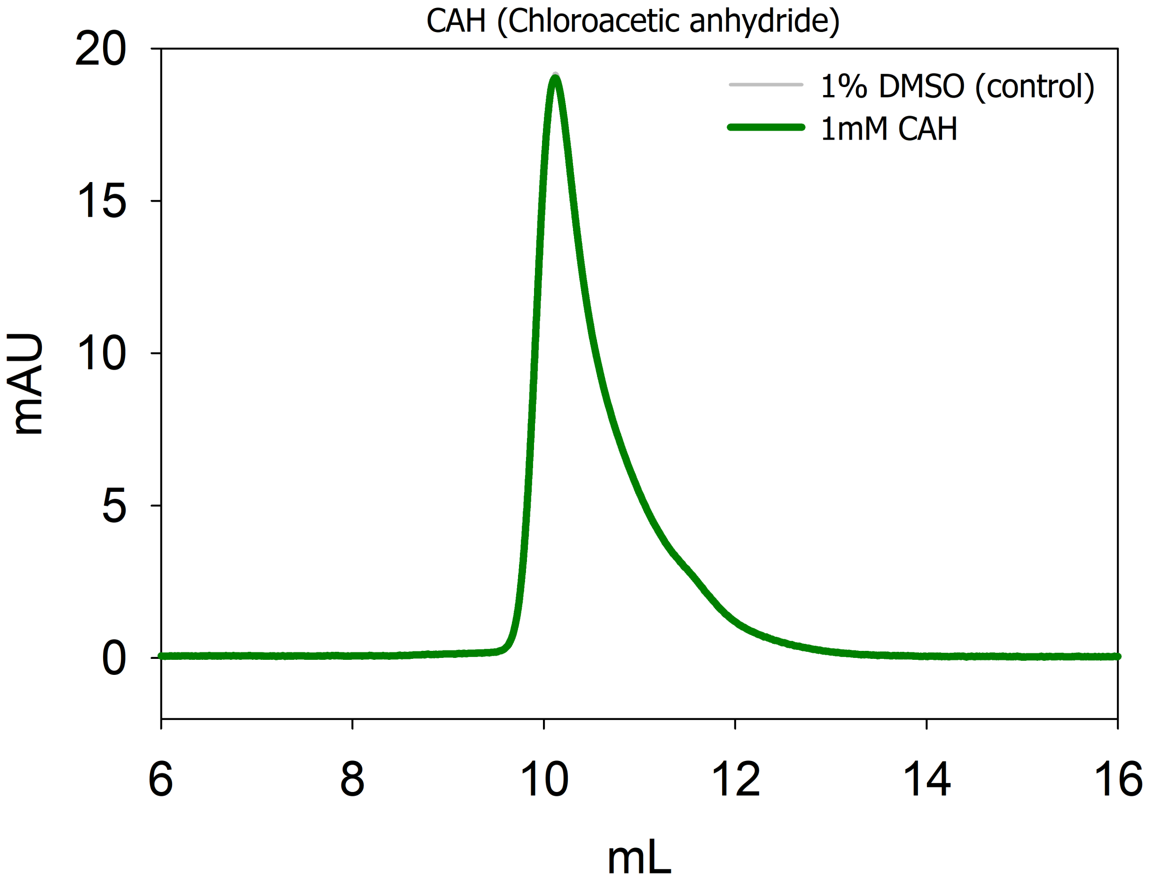
CAH (Chloroacetic anhydride)
1% DMSO (control)
1mM CAH
mAU
mL
20
15
10
5
0
6
8
10
12
14
16

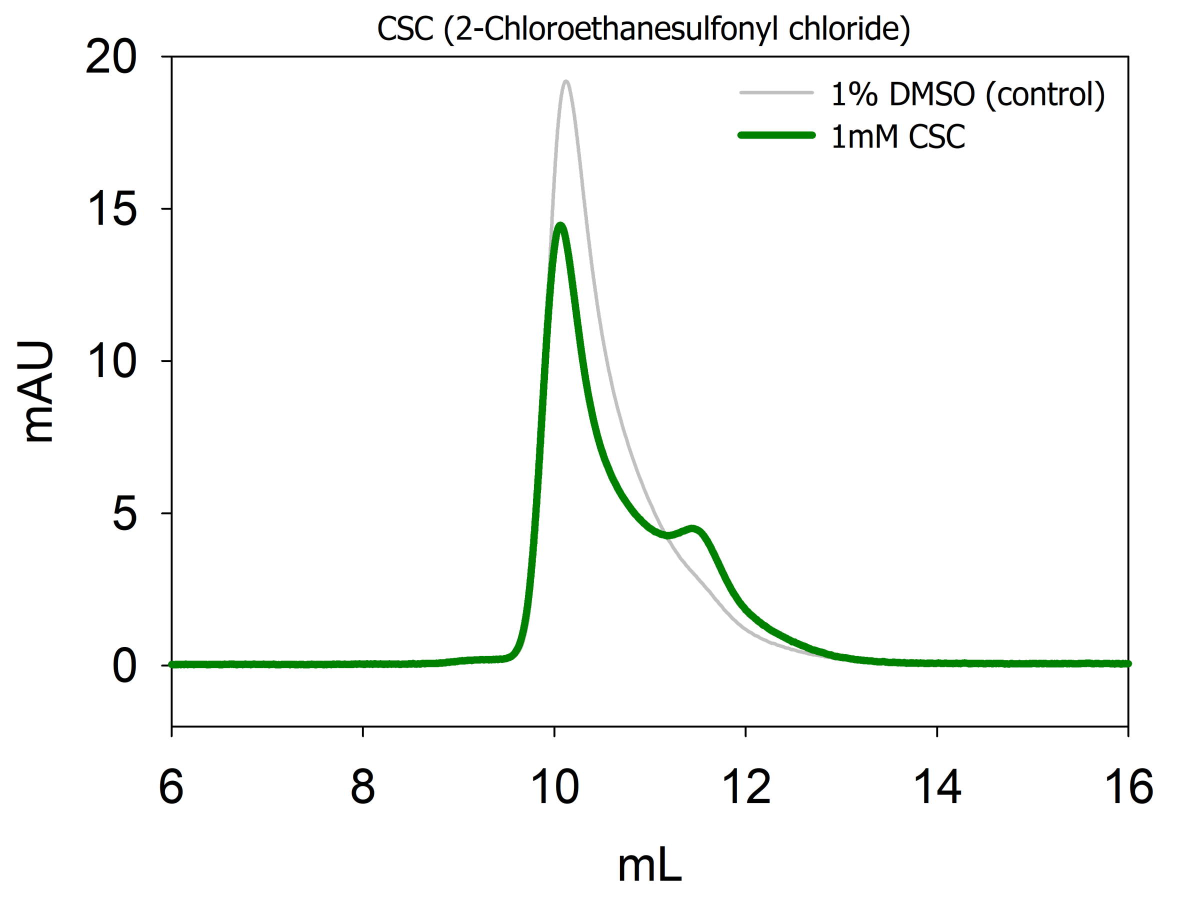
CSC (2-Chloroethanesulfonyl chloride)
1% DMSO (control)
1mM CSC
mAU
20
15
10
5
0
6
8
10
12
14
16
mL
